# Assessing the quality of comparative genomics data and results with the *cogeqc* R/Bioconductor package

**DOI:** 10.1101/2023.04.14.536860

**Authors:** Fabricio Almeida-Silva, Yves Van de Peer

## Abstract

Comparative genomics has become an indispensable part of modern biology due to the advancements in high-throughput sequencing technologies and the accumulation of genomic data in public databases. However, the quality of genomic data and the choice of parameters used in software tools used for comparative genomics can greatly impact the accuracy of results. To address these issues, we present *cogeqc*, an R/Bioconductor package that provides researchers with a toolkit to assess genome assembly and annotation quality, orthogroup inference, and synteny detection. The package offers context-guided assessments of assembly and annotation statistics by comparing observed statistics to those of closely-related species on NCBI. To assess orthogroup inference, *cogeqc* calculates a protein domain-aware orthogroup score that aims at maximizing the number of shared protein domains within the same orthogroup. The assessment of synteny detection consists in representing anchor gene pairs as a synteny network and analyzing its graph properties, such as clustering coefficient, node count, and scale-free topology fit. The application of cogeqc to real data sets allowed for an evaluation of multiple parameter combinations for orthogroup inference and synteny detection, providing researchers with guidelines to aid in the selection of the most appropriate tools and parameters for their specific data.

## Introduction

Comparative genomics has revolutionized our understanding of (the genomic basis of) biology. The advent of high-throughput sequencing technologies and the availability of genomic data from an ever-growing number of species have transformed this field and opened up new avenues for discovery (Alföldi & Lindblad-Toh, 2013; Sun et al., 2021; Yousaf et al., 2021). Over the years, large-scale genome comparisons have resulted in numerous groundbreaking discoveries, providing critical insights into the evolution of life (Koonin, 2003; Li et al., 2021; Prüfer et al., 2014; Van de Peer et al., 2017), the genetic basis of species differences and adaptation to changing environments (Konstantinidis & Tiedje, 2005; Wan et al., 2021; Yuan et al., 2021), and the evolution of novel traits (Palfalvi et al., 2020; M. Wu et al., 2018). As a result, comparative genomics analyses have become an integral part of modern biology, being routinely used in the life sciences.

The advancements made in sequencing technologies have led to a sharp increase in the amount of genomic data available in public databases, presenting a wealth of information for comparative genomics research. However, unfortunately, much is of poor quality (Feron & Waterhouse, 2022; Marks et al., 2021). Common analyses in comparative genomics, such as genome-wide synteny analysis and gene orthology detection, rely on high-quality data to accurately identify similarities and differences between genomes (Liu et al., 2018; P. Wang & Wang, 2022). Poor quality data, on the other hand, can lead to incorrect or unreliable results, making it difficult to draw meaningful conclusions about the evolution of species and their genomes. Therefore,careful quality control and validation of genomics data is key before such data can be used effectively for comparative genomics research.

In addition to data quality, results obtained from comparative genomics software can be greatly influenced by the choice of parameters used in the analyses. Different software tools offer a wide range of parameters for users to choose from, and these parameters can significantly impact the results obtained (Buchfink et al., 2021; Emms & Kelly, 2019). Default parameters in software tools are typically optimized for particular (usually gold standard) data sets, which means that they may not be suitable for other data sets that have different characteristics, such as greater heterogeneity, or lower quality. Thus, having a set of tools to assess the quality of results under different combinations of parameters can help improve the accuracy and reliability of the results.

Here, we present *cogeqc*, an R/Bioconductor package that can be used as a toolkit for assessing genome assembly and annotation statistics, orthogroup inference, and synteny detection. The package offers context-guided assessments of assembly and annotation statistics by comparing observed values to those of closely-related species on the National Center for Biotechnology Information (NCBI), while gene space completeness can be assessed with best universal single-copy orthologs (BUSCOs). The orthogroup inference assessment uses a protein domain-aware orthogroup score to maximize the number of shared protein domains within the same orthogroup. Finally, the assessment of synteny detection relies on representing anchor pairs as a synteny network and analyzing its graph properties. The application of *cogeqc* to real datasets allowed for an evaluation of multiple parameter combinations for orthogroup inference and synteny detection, providing researchers with guidelines to aid in the selection of the most appropriate parameters for their specific data.

## Materials and Methods

### Implementation

*cogeqc* is part of the Bioconductor ecosystem and, as such, can be easily integrated with other Bioconductor packages. Input data types are either base R or core Bioconductor classes (*e*.*g*., *DNAStringSet* and *AAStringSet* objects for DNA and protein sequences, respectively). For integration with external software tools (*i*.*e*., *BUSCO* (Simão et al., 2015) and OrthoFinder (Emms & Kelly, 2019)), we provide users with functions to read and parse their output for downstream analyses in *cogeqc*.

### Assessing genome assembly and annotation statistics

We propose a context-guided assessment of assembly and annotation statistics that consists in comparing observed values for common metrics (e.g., genome size, contiguity measures, number of genes, etc.) with those of closely-related species on the National Center for Biotechnology Information (NCBI). For a particular taxon, the function *get_genome_stats()* extracts summary assembly and annotation statistics for all genomes on NCBI via the Datasets REST API (https://www.ncbi.nlm.nih.gov/datasets/) and returns a data frame with information on 35 variables, such as assembly level, scaffold and contig contiguity measures, number of coding and non-coding genes, and submitter data. In addition to the statistics already provided by the NCBI, the output data frame includes a variable ‘CC ratio’ representing the ratio of the number of contigs to the number of chromosome pairs, which has recently been proposed by (P. Wang & Wang, 2022) as a contiguity measure that compensates for the flaws of N50/L50 and allows cross-species comparisons.

Additionally, users can create a data frame containing assembly and annotation statistics for their own genome projects and pass it to the *compare_genome_stats()* function along with the output of *get_genome_stats()*. This function will add the user-provided statistics to a distribution of reference statistics from NCBI quality-checked genomes and report the percentile and rank of observed values in the distribution. Observed statistics can also be visually compared with reference statistics in publication-ready plots created by the function *plot_genome_stats()* (Fig. 1A). Such context-guided assessments are particularly useful in cases when assembly statistics seem problematic (*e*.*g*., genome size is too large, number of genes is too small), but which are in fact due to genomic features of particular taxa. For instance, adaptation to unusual habitats are often associated with massive gene loss, as observed for parasitic (Xu et al., 2021), carnivorous (Palfalvi et al., 2020), and aquatic plants (An et al., 2019). Likewise, genomes with higher transposable element contents are typically larger (Michael, 2014). Thus, comparing observed statistics with those from closely-related genomes can help distinguish assembly and annotation issues from expected biological phenomena.

**Fig. 1.**
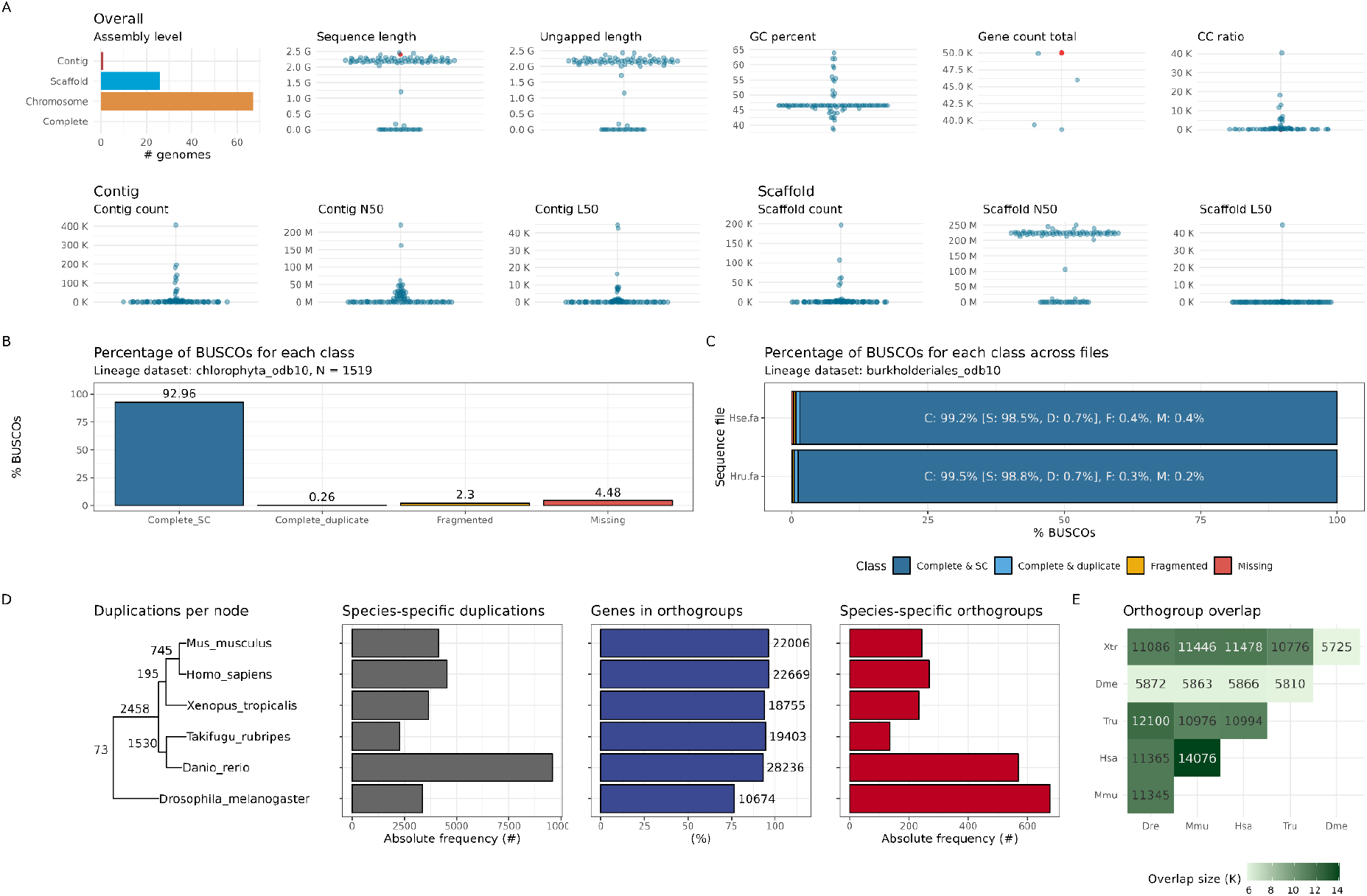
Summary of publication-ready plots that can be created with graphical functions in *cogeqc*. **A**. Summary assembly and annotation statistics for all *Zea mays* genomes on the NCBI obtained with the function *plot_genome_stats()*. Red data points in the “Sequence length” and “Gene count total” panels represent simulated observed values passed by the user. **B**. BUSCO summary statistics for a single genome obtained with the function *plot_busco()*. The data used in this figure is a BUSCO output for the *Ostreococcus tauri* genome. **C**. BUSCO summary statistics for multiple genomes obtained with the function *plot_busco()*. The data used in this figure is a BUSCO output in batch mode for the bacteria *Herbaspirillum seropedicae* and *Herbaspirillum rubrisubalbicans*. **D**. Summary comparative genomics statistics from OrthoFinder obtained with the function *plot_orthofinder_stats()*. **E**. Heatmap of orthogroup overlap for pairwise species comparisons as obtained with the function *plot_og_overlap()*. All data used to create the figures are distributed with the package as example data sets.

### Assessing gene space completeness

To assess gene space completeness, *cogeqc* relies on the identification of best universal single-copy orthologs (BUSCOs) (Simão et al., 2015). The function *run_busco()* is a wrapper that takes a sequence as input (either as a FASTA file or as *AA/DNA/RNAStringSet* objects), runs BUSCO from the R session, and returns a data frame with the frequency of complete (duplicate and single copy), fragmented, and missing BUSCOs. Users can also input the path to a directory containing multiple FASTA files, so *run_busco()* will run BUSCO in batch mode and return a data frame with summary BUSCO scores for all files. Alternatively, if users have already run BUSCO through the command line, they can use the function *read_busco()* to read and parse the BUSCO output file as a data frame. Finally, the function *plot_busco()* can be used to create publication-ready summary plots for both single-genome and batch modes (Fig. 1B and 1C).

### Assessing orthogroup inference

We developed a protein domain-aware orthogroup assessment score that aims to maximize the number of shared protein domains within the same orthogroup while minimizing the number of different orthogroups containing the same protein domain. The rationale for such approach is that genes that share the same protein domain are expected to have evolved from a common ancestor, so they should be assigned to the same orthogroup. Formally, orthogroup scores are calculated as:

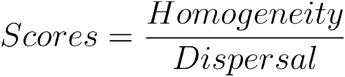

The numerator, *homogeneity*, is the mean Sorensen-Dice index for all pairwise combinations of genes in an orthogroup. The Sorensen-Dice index measures how similar two genes are in terms of the protein domains they have, and it ranges from 0 to 1, with 0 meaning that a gene pair does not share any protein domain, and 1 meaning that it shares all protein domains. Formally:

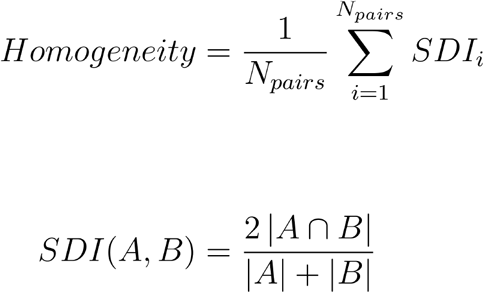

where A and B are the set of protein domains associated with genes A and B. Hence, an 1orthogroup with score 1 would have all genes with the exact same protein domains, while an orthogroup with score 0 would have a different protein domain for each gene. As individual genes in a gene family can lose domains and gain new ones, orthogroup scores can take any value from 0 to 1.

The denominator, *dispersal*, aims to correct for overclustering (*i*.*e*., orthogroup assignments that break ‘true’ gene families into an artificially large number of smaller subfamilies), and it describes the mean number of orthogroups containing the same protein domain normalized by the number of orthogroups. Formally:

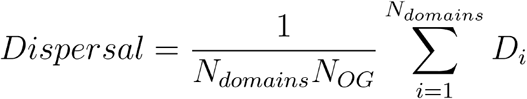

where *N*_*OG*_, is the number of orthogroups, and *D*_*i*_, is the number of orthogroups containing the protein domain *i*. This term penalizes orthogroup assignments with the same protein domains in multiple orthogroups. We acknowledge that the presence of the same protein domain in multiple orthogroups can occur due to convergent evolution, but since convergent evolution of protein domains is rare (Gough, 2005), we assume that such patterns indicate overclustering of gene families.

With *cogeqc*, orthogroups inferred with OrthoFinder (Emms & Kelly, 2019) can be read and parsed as a data frame with the function *read_orthogroups()*. The function *assess_orthogroups()* takes this data frame as input and calculates orthogroup scores for each species, as well as mean and median scores for each orthogroup. To ensure a higher accuracy in orthogroup assignments, we recommend running OrthoFinder with different combinations of parameters and comparing the distributions of orthogroup scores in each run to select the best. Alternatively, if reference and reliable orthogroup assignments exist, users can compare their predicted orthogroups with reference orthogroups by using the function *compare_orthogroups()*, which can show the percentage of reference orthogroups that are preserved in predicted orthogroups.

Finally, comparative genomics statistics obtained with OrthoFinder can be read as a list of data frames with the function *read_orthofinder_stats()*. This list can be passed as input to graphical functions that create publication-ready plots summarizing statistics, such as *plot_orthofinder_stats(), plot_og_overlap()*, and *plot_og_sizes()* (Fig. 1D and 1E).

### Assessing synteny detection

We propose a network-based assessment of synteny (or collinearity, used here as synonyms) detection that consists in representing synteny relationships as a graph (*i*.*e*., a synteny network) and analyzing topological properties of the graph to assess its quality. To infer synteny networks for mammalian and angiosperm genomes, (Zhao & Schranz, 2019) have run a synteny detection algorithm with multiple combinations of parameters and selected the best combination based on the clustering coefficient and number of nodes of each network. Ideally, a synteny network should have a large number of nodes (*i*.*e*., anchor pairs, duplicated genes retained from a large-scale duplication event) and a high clustering coefficient.

However, there is often a trade-off between the number of nodes and the clustering coefficient, with larger networks being more sparse, and smaller networks being more densely connected. To account for this trade-off, we use the product of the clustering coefficient and the number of nodes to assess networks. Additionally, as synteny networks and biological networks in general tend to be scale-free (*i*.*e*., the degree distribution follows a power-law distribution) (Barabási, 2009; Barabasi & Oltvai, 2004; Ravasz et al., 2002; Venancio et al., 2009; Zhao & Schranz, 2019), we added a term to the network score formula that considers how well the network fits a scale-free topology. Formally, the score of a synteny network is calculated as:

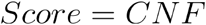

where *C* is the network’s clustering coefficient, *N* is the number of nodes, and *F* is the coefficient of determination (R^2^) for the scale-free topology fit.

Synteny networks can be inferred with the R package syntenet (Almeida-Silva et al., 2023). The score of a synteny network can be calculated with the function *assess_synnet()*, which returns a data frame with the network’s score and the observed values for each term of the formula above (*C, N*, and *F*). If users have multiple networks stored in a list, the function *assess_synnet_list()* can calculate scores for multiple networks at once.

### Benchmark data

Chlorophyta genomes for the BUSCO assessments were obtained from the Pico-PLAZA 3.0 database, and Fabaceae genomes for the synteny detection benchmark were obtained from PLAZA 5.0 (Van Bel et al., 2022). Brassicaceae genomes for the orthogroup assessments were obtained from PLAZA 5.0, Phytozome v13, BRAD, and CoGe (Cheng et al., 2011; Goodstein et al., 2012; Lyons et al., 2008; Van Bel et al., 2022). Enrichment analyses were performed with the Bioconductor package clusterprofiler (T. Wu et al., 2021).

## Results and Discussion

### Assessing the completeness of Chlorophyta genomes

To demonstrate the usage of the functions to assess gene space completeness, we obtained genome sequences for all Chlorophyta genomes on Pico-PLAZA 3.0 (*N* = 16) and calculated their BUSCO scores (Supplementary Text S1). All genomes were stored in the same directory and the function *run_busco()* was used to run BUSCO in batch mode using the lineage dataset *chlorophyta_odb10*. BUSCO scores were visualized with the function *plot_busco()* (Fig. 2). We observed that Chlorophyta genomes on Pico-PLAZA are highly complete, with >90% complete BUSCOs (Fig. 2). However, an exception is the alga *Helicosporidium sp*. (Trebouxiophyceae), with only 65.3% complete BUSCOs.

**Fig. 2.**
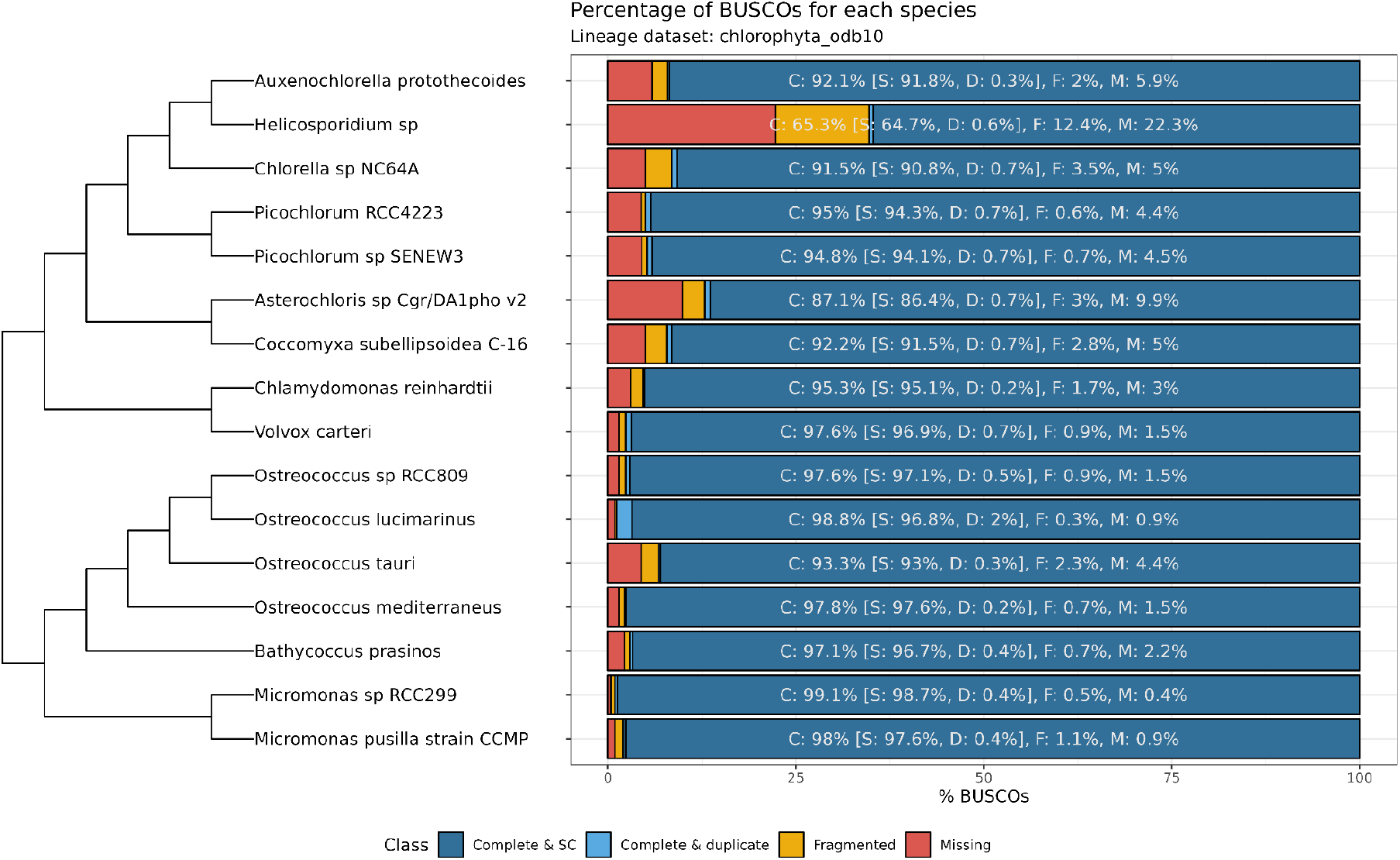
Species tree and BUSCO scores for Chlorophyta genomes in Pico-PLAZA 3.0. Scores were calculated using the function *run_busco()* to run BUSCO in batch mode using the chlorophyta_odb10 as lineage data set. Except for *Helicosporidium sp*., all species have highly complete genomes, as demonstrated by their percentages of complete BUSCOs. Figures were generated by combining the output of the functions *plot_species_tree()* and *plot_busco()*, and code to recreate it is available in Supplementary Text S1.

### Orthogroup assignments in different public databases perform equally well

We used the protein domain-aware orthogroup assessment implemented in *cogeqc* to assess orthogroup assignments in public databases, namely PLAZA Dicots 5.0 (Van Bel et al., 2022), OrthoDB (Kuznetsov et al., 2023), eggNOG (Hernández-Plaza et al., 2023), and HOGENOM (Penel et al., 2009). As the species composition varies across databases, we used *Arabidopsis thaliana* as a representative species to assess orthogroups. For each database, orthogroups were filtered to include only *Arabidopsis* genes, and scores were calculated with the function *calculate_H()* using InterPro domain annotation obtained from PLAZA Dicots 5.0 (Supplementary Text S2).

We observed that eggNOG orthogroups have lower scores than orthogroups from all other databases (Mann-Whitney U test, *P* <0.01). HOGENOM orthogroup scores are higher than OrthoDB scores, but lower than PLAZA. Finally, PLAZA orthogroup scores are higher than all other databases (Supplementary Text S2, section 3). However, although differences were significantly different (Mann-Whitney U test, *P* <0.01), Wilcoxon effect sizes (r) were small, with the difference between eggNOG and HOGENOM being the only one with r > 0.1 (Fig. 3A). The small effect sizes suggest that the observed differences could be due to large sample sizes, as small *P*-values can be obtained if the sample size is large enough, even when differences are negligible (Sullivan & Feinn, 2012). Thus, despite some small differences, we conclude that all databases perform equally well in their orthogroup assignments, as their distributions of homogeneity scores were highly similar.

**Fig. 3.**
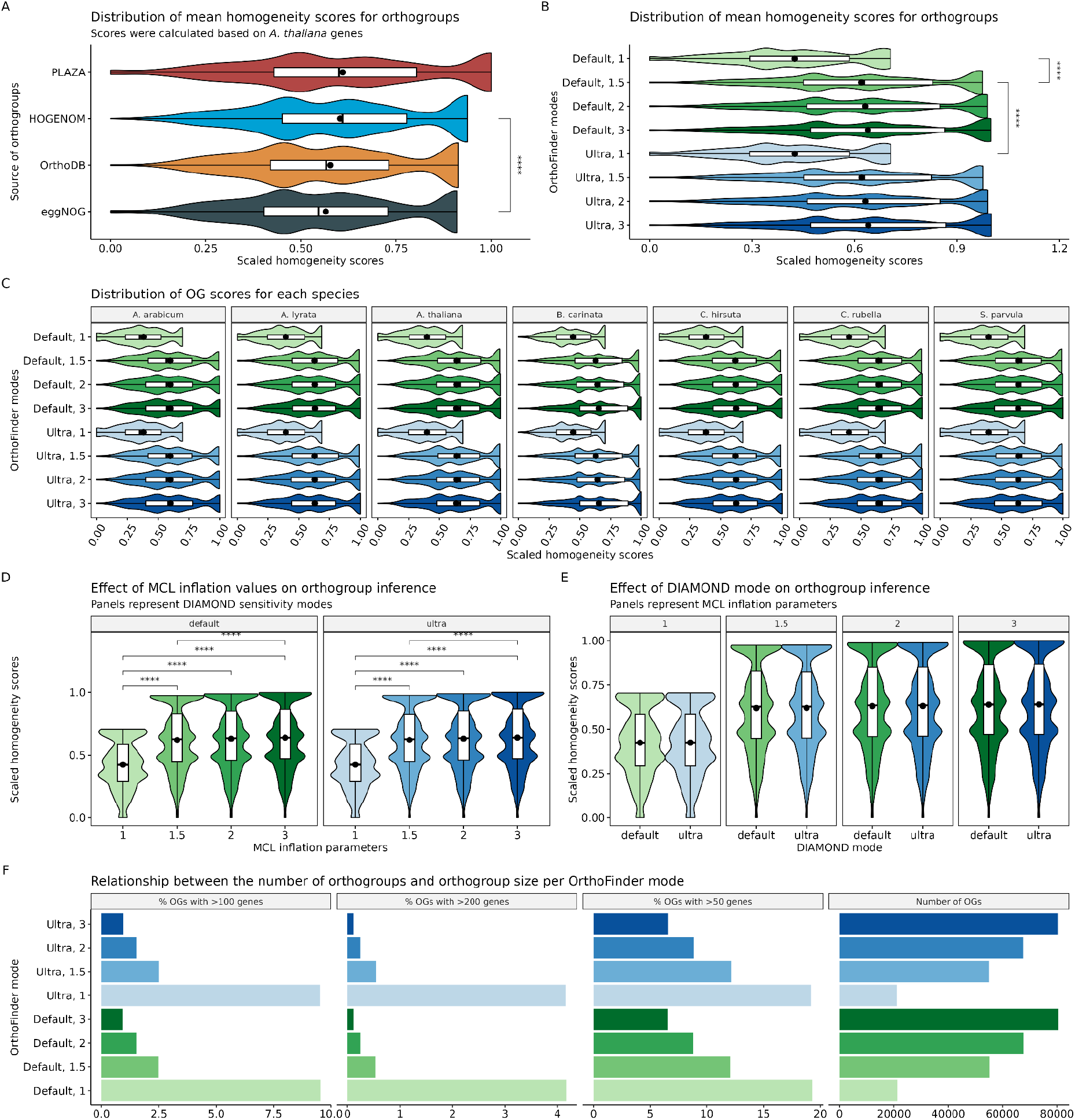
Assessment of orthogroup inference in public databases and for a Brassicaceae data set. **A**. Distribution of orthogroup scores for each database. Comparisons with Wilcoxon effect sizes ≥0.1 are highlighted, with asterisks representing significance levels. **B**. Distribution of mean orthogroup scores for each OrthoFinder run. Comparisons between the default OrthoFinder mode (standard DIAMOND, *mcl* = 1.5) and other runs with Wilcoxon effect sizes ≥0.1 are highlighted. **C**. Distribution of orthogroups scores obtained when considering each species individually. Distributions have the same shape regardless of the species choice. **D**. Comparison of the distributions of orthogroup scores for OrthoFinder runs with the same DIAMOND mode, but different *mcl* values. Significant differences (P <0.05, Mann-Whitney U test) are highlighted. **E**. Comparison of the distributions of orthogroup scores for OrthoFinder runs with the same *mcl* value, but different DIAMOND modes. **F**. Bar plots displaying the relationship between orthogroup count and size for each OrthoFinder run. Green and blue bars/distributions represent OrthoFinder runs with the default and ultra-sensitive DIAMOND modes, respectively. *, P ≤0.05. **, P ≤0.01, **. ***, P ≤0.001. ****, P ≤0.0001.

### Assessing orthogroup inference under multiple combinations of OrthoFinder parameters

To infer orthogroups from a set of proteomes for *N* species, OrthoFinder relies on similarity searches with DIAMOND (Buchfink et al., 2021), followed by normalization of bit scores by gene length, and graph-based clustering using Markov clustering (MCL). By default, OrthoFinder runs DIAMOND in default mode, which is faster, but less accurate than its ultra-sensitive mode. To cluster genes into orthogroups, an MCL inflation parameter of 1.5 is used by default, with lower values resulting in a smaller number of large clusters (*i*.*e*., low granularity), and higher values resulting in a greater number of small clusters (*i*.*e*., high granularity). To date, little is known about how changing the DIAMOND mode and MCL inflation parameter might influence the accuracy of orthogroup inference.

To investigate this issue, we ran OrthoFinder on a Brassicaceae data set (*N* = 25) with eight combinations of parameters by changing the DIAMOND mode (standard vs ultra-sensitive mode) and the Markov clustering inflation (*mcl* = 1, 1.5, 2, and 3). Orthogroup scores for each OrthoFinder run were obtained with the function *assess_orthogroups()* (Supplementary Text S3). A global comparison of the distributions of orthogroup scores shows that using an MCL inflation parameter of 1 dramatically reduces homogeneity scores as compared to every other *mcl* value (Mann-Whitney U test, P < 0.01; Fig 3B). Orthogroup scores for the default OrthoFinder mode (standard DIAMOND, *mcl* = 1.5) are much larger than runs with *mcl = 1*, both with standard and ultra-sensitive DIAMOND mode (Mann-Whitney U test, P < 0.001, effect size > 0.3; Fig. 3B). To test for a possible bias resulting from the species choice, we inspected the distributions of orthogroup scores by OrthoFinder mode for each species separately. We observed that the species choice does not affect scores, as revealed by similar distribution shapes for all species (Fig. 3C).

Furthermore, we analyzed the effects of changing arguments for each parameter separately (*i*.*e*., same DIAMOND mode with different *mcl* values, and vice versa) to understand their individual relevance to orthogroup scores. We observed that increasing *mcl* values leads to significantly higher orthogroup scores, with scores following the order 3 > 2 > 1.5 > 1, but the difference is only large between *mcl* values of 1 and other values (Mann-Whitney U test, P < 0.001, r >0.3; Fig. 3D). Wilcoxon effect sizes for the comparisons between *mcl* values of 1.5 and above are small (r <0.1; Tables 3 and 4 in Supplementary Text S3), suggesting that significant differences could be due to large sample sizes (*N* > 16,000). Likewise, comparisons of orthogroup scores for OrthoFinder runs with different DIAMOND modes revealed significant, but negligible differences (P <0.05, r <0.04; Fig. 3E; Table 5 in Supplementary Text S3), indicating that changing DIAMOND modes has little impact in orthogroup scores and, hence, should not be a concern in orthogroup detection. Thus, we recommend running OrthoFinder with the standard DIAMOND mode, because it offers a 100-fold increase in speed compared to the ultra-sensitive mode (Buchfink et al., 2021).

### Guidelines for using OrthoFinder to study gene family evolution

We showed that running OrthoFinder with the standard DIAMOND mode is a better approach, because it is dramatically faster without any effect on orthogroup scores (Fig. 3E). However, recommendations on which *mcl* value to use were unclear, as differences in orthogroup scores are very small for runs with *mcl* ≥1.5 (Fig. 3D). To provide readers with clearer guidelines, we compared OrthoFinder runs in terms of orthogroup scores, number of orthogroups, and orthogroup sizes. As expected, we observed that increasing *mcl* values leads to a greater number of orthogroups, but of smaller size (Fig. 3F). When comparing OrthoFinder runs with *mcl* values of 3 and 1.5 (default), we observed a 2.7-fold increase in the percentage of orthogroups with ≥100 genes for the latter.

Most of the models to study gene family evolution rely on phylogenetic birth-and-death processes (BDPs), such as those implemented in CAFE 5 (Mendes et al., 2020), Count (Csuros, 2010), and DeadBird (Zwaenepoel & Van de Peer, 2020). In such models, large variances in gene copy number can lead to non-informative parameter estimates, and a common rule of thumb consists in removing orthogroups with ≥100 genes (Mendes et al., 2020). Thus, since large gene families are typically discarded, a smaller number of orthogroups with ≥100 genes is desired. Hence, we recommend running OrthoFinder with *mcl* values of 3, as it halves the percentage of discarded orthogroups and leads to slightly higher orthogroup scores.

### OrthoFinder’s normalized bit scores correct for gene length biases, but not completely

Inferring orthogroups consists in applying a clustering algorithm (typically Markov clustering) to bit scores obtained from similarity search programs. However, Emms & Kelly (2015) demonstrated that raw bit scores are a biased measure of similarity, because short sequences cannot obtain high bit scores, and long sequences often have higher scores than those obtained for the best hits of short sequences. To address this issue, OrthoFinder implements a gene length normalization that seems to eliminate such bias, as there is no correlation between gene length and bit scores after applying normalization (Emms & Kelly, 2015). Although such lack of correlation suggests that the gene length bias is removed, the effect of gene length on orthogroup content remains to be investigated.

To verify the effectiveness of OrthoFinder’s normalization, we fitted a linear model using log-transformed gene lengths as predictors and log-transformed orthogroup scores as outcomes. The model fit displayed a significant negative association (P <0.001, R^2^ = -0.2), suggesting some tendency of orthogroups with longer genes to have lower scores (Fig. 4A). However, the model fit is not perfect for the data, as only 20% of the variance in orthogroup scores is explained by gene length. Thus, we conclude that the normalization implemented in OrthoFinder sufficiently corrects for gene length biases, but there might still be room for improvement.

**Fig. 4.**
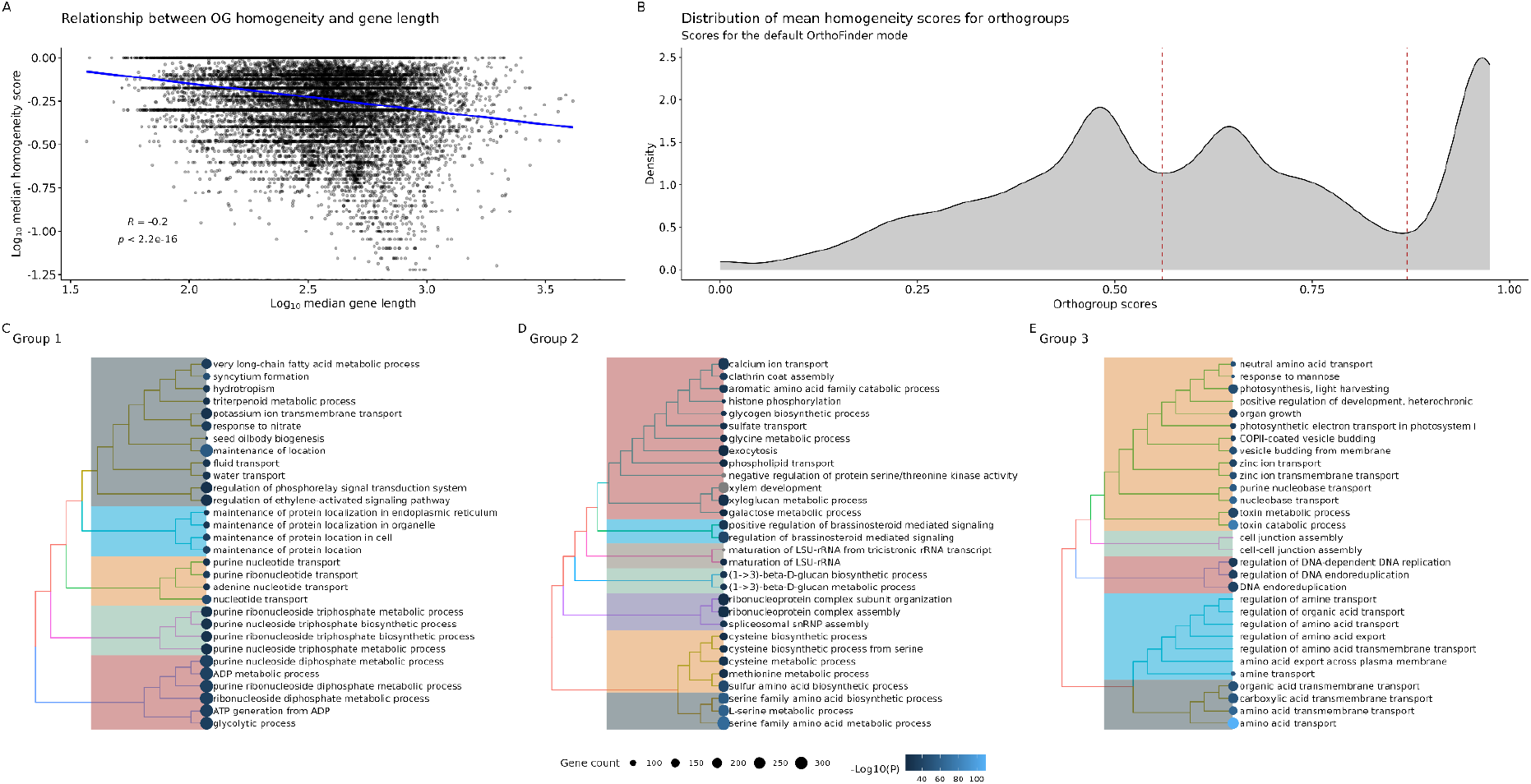
Validation of OrthoFinder’s bit score normalization and functional analyses of orthogroups clusters. **A**. Relationship between orthogroup scores and orthogroup median length. The blue line represents a linear regression fit. The model shows a significant association, but only 20% of the variance in orthogroup scores is explained by orthogroup length. **B**. Distribution of orthogroup scores for an OrthoFinder run with default parameters. Peaks were used to split the distribution in 3 clusters, with boundaries indicated by dashed red lines. **C**. Tree plot of enriched functional terms for each orthogroup cluster. Enrichment analyses were performed with clusterProfiler (T. Wu et al., 2021), and visualized with enrichplot. Terms are grouped in a tree-like structure based on semantic similarities calculated with the function *pairwise_termsim()* from the enrichplot package.

### Functional analyses unveil biological processes associated with rapidly and slowly evolving gene families

Although some combinations of OrthoFinder parameters lead to higher orthogroup scores, we noticed that all distributions have a similar shape. Upon closer inspection with the default OrthoFinder mode (standard DIAMOND, *mcl* = 1.5), we were able to split the distribution of orthogroup scores in three clusters based on peaks (Fig. 4B). Orthogroups in cluster 1 have the lowest scores (from 0 to 0.56), indicating that protein domains are shared by only a small fraction of the genes thereof and, hence, that such orthogroups are gaining and losing domains at faster rates. Orthogroups in cluster 3, on the contrary, have the highest scores (from 0.87 to 1), indicating that protein domains are shared by most (or all) of their members, and suggesting slower evolutionary rates. Finally, orthogroups in cluster 2 have intermediate scores (from 0.56 to 0.87), demonstrating that their evolutionary rates are neither fast nor slow.

To explore the functional profiles of each orthogroup cluster, we performed enrichment analyses of GO terms (biological process only), MapMan bins, and InterPro domains. All genes in orthogroups were used as background. For cluster 1 (lowest orthogroup scores), we found an enrichment of genes associated with ATP production, water and K^+^ transport, seed oil body biogenesis, and response to nitrate and ethylene (Fig. 4C). Orthogroups in cluster 2 were enriched in genes associated with sulfur amino acid metabolism, spliceosome biogenesis, □-1,3-glucan biosynthesis, response to brassinosteroids, xylem development, exocytosis, and calcium and sulfate transport. Finally, orthogroups in cluster 3 were enriched in genes involved in photosynthesis, zinc and amino acid transport, DNA replication, endocytosis, cell-cell junction assembly, and toxin catabolism.

Evolutionary rate variations among gene families have been extensively studied, and they typically distinguish families involved in housekeeping processes from those involved in environmental responses (Guo, 2013; Hahn et al., 2005). For instance, gene families associated with immune response and transcriptional regulation evolve rapidly due to the pressure to adapt to changing environments (Ngou et al., 2022; P. Wang et al., 2018). On the other hand, gene families involved in housekeeping processes, such as DNA replication and ribosome biogenesis, tend to evolve slowly due to the high cost of disruptive mutations in these genes (P. Wang et al., 2018). Thus, our findings are in line with previous observations, as we observed that rapidly evolving families are associated with environmental response (e.g., nitrate and ethylene), while slowly evolving families are associated with more basic cellular processes, such as DNA replication and photosynthesis (Fig. 4C).

### Graph-based assessment of synteny detection with different combinations of parameters

To detect synteny between genomic regions, one must explicitly define the minimum number of genes required to call a syntenic block or segment, and the maximum number of allowed gaps between genes. For example, the MCScanX algorithm (Y. Wang et al., 2012) requires a syntenic block to have a minimum of 5 genes by default (*i*.*e*., -s 5), and a maximum gap of 20 genes is allowed (-m 20). Using a large dataset of mammalian and angiosperm genomes, Zhao and Schranz (2019) demonstrated that a better synteny detection can be achieved by changing default values. Here, we used the R package syntenet (Almeida-Silva et al., 2023) to detect synteny among Fabaceae genomes with 5 combinations of parameters: *a3m25, a5m15, a5m25, a5m35*, and *a7m25*, where ‘a’ stands for the minimum number of anchors to call synteny, and ‘m’ stands for the maximum number of gaps between genes. Anchor pairs were represented as a synteny network, and we assessed both the complete network (including all 9 Fabaceae species) and each species’ network.

For the full network, scaled network scores were very similar, but the parameter combination *a3m25* resulted in the best synteny network, with the largest number of nodes, scale-free topology fit, and overall score (Fig. 5A). Interestingly, the network obtained with the default parameter combination of the MCScanX algorithm, *a5m25*, had the lowest score. When analyzing each species’ network separately, we observed that the best parameter combination depends on the species (Fig. 5B). This finding highlights the importance of performing quality checks and assessments for each data set, as there is no universally optimal combination. However, we noticed some patterns for the Fabaceae data set. The combinations *a7m25* and *a5m15* are typically the worst, leading to zero scores in some species due to clustering coefficients of zero (Fig. 5B). Thus, users who want to test multiple parameter combinations for their own data sets can skip these combinations to save time. The combinations *a3m25, a5m25*, and *a5m35* lead to the best scores in 45%, 33%, and 22% of the species, respectively. Finally, we observed that the parameter combination that leads to the highest score in most species-specific networks (*a3m25*) is also the best combination for the full network (with all species) (Fig. 5B). Although it suggests that this combination tends to be the best in most cases, we advise users to test multiple combinations whenever possible.

**Fig. 5.**
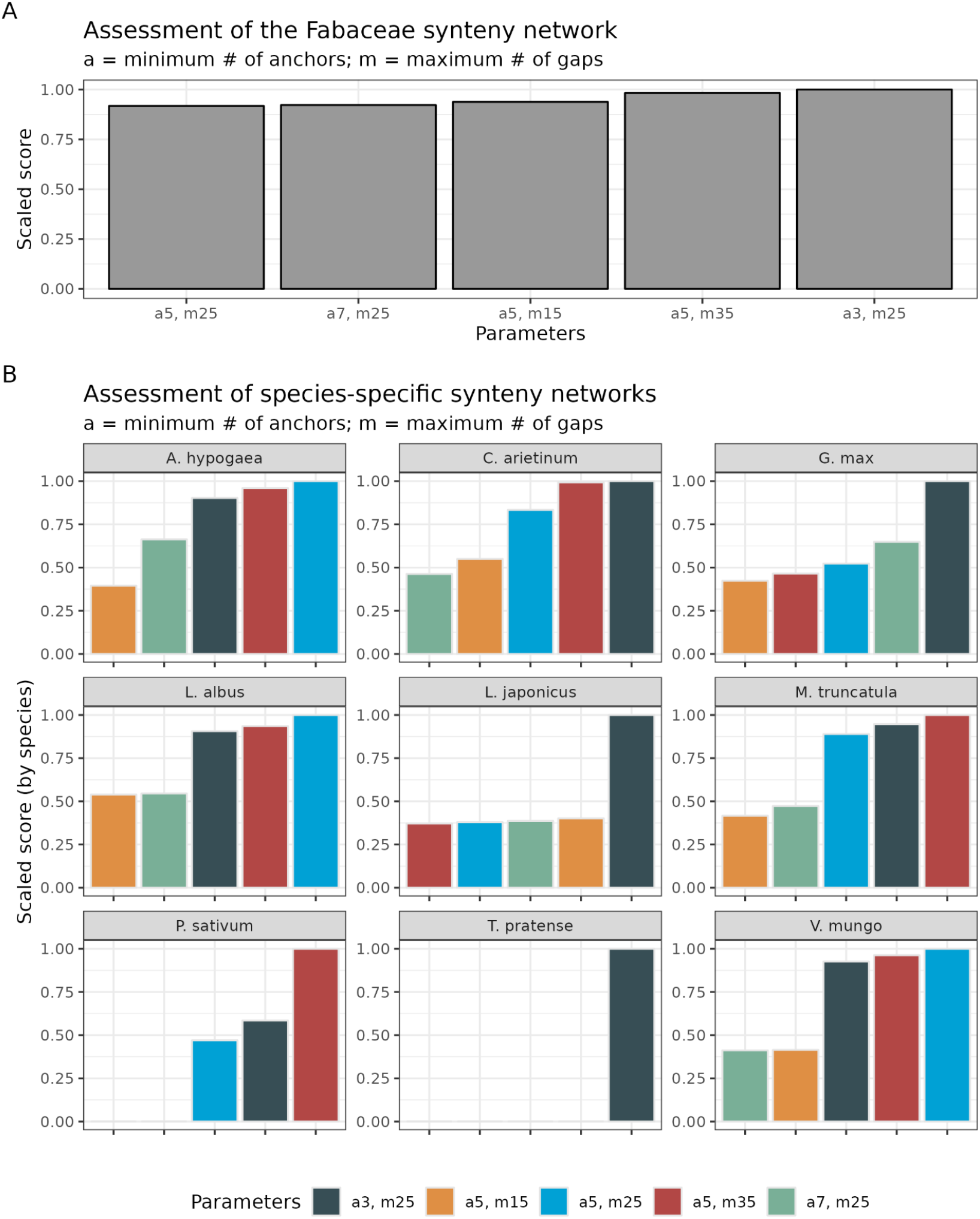
Assessment of synteny detection in Fabaceae species with different combinations of parameters. **A**. Network scores for a synteny network containing all Fabaceae species. **B**. Network scores for species-specific synteny networks. Networks with zero scores are due to clustering coefficients of 0. Colors represent different combinations of parameters.

## Conclusion

*cogeqc* is an R package that can be used to assess the quality of genome assembly and annotation in a phylogenetic context, and to assess the quality of orthogroup inference and synteny detection. The package was designed to be user-friendly, easily integrate with other packages and software tools, and provide summary results as publication-ready plots. Applications to real data sets demonstrated how the package can be used to select optimal parameters in orthogroup inference and synteny analyses, providing users with general guidelines.

## Acknowledgements

YVdP acknowledges funding from the European Research Council (ERC) under the European Union’s Horizon 2020 research and innovation program (No. 833522). YVdP and FA-S acknowledge funding from Ghent University (Methusalem funding, BOF.MET.2021.0005.01).

## Data availability

To ensure full reproducibility, all code and data used in this manuscript are available in a GitHub repository at https://github.com/almeidasilvaf/cogeqc_paper, and the code used in benchmarks are available in Supplementary Texts.

## Notes

### Competing Interest Statement

The authors have declared no competing interest.

## REFERENCES

Alföldi, J., & Lindblad-Toh, K. (2013). Comparative genomics as a tool to understand evolution and disease. Genome Research, 23(7), 1063–1068.

Almeida-Silva, F., Zhao, T., Ullrich, K. K., Schranz, M. E., & Van de Peer, Y. (2023). syntenet: An R/Bioconductor package for the inference and analysis of synteny networks. Bioinformatics, 39(1),btac806.

An, D., Zhou, Y., Li, C., Xiao, Q., Wang, T., Zhang, Y., Wu, Y., Li, Y., Chao, D.-Y., & Messing, J. (2019). Plant evolution and environmental adaptation unveiled by long-read whole-genome sequencing of Spirodela. Proceedings of the National Academy of Sciences, 116(38), 18893–18899.

Barabási, A.-L. (2009).Scale-free networks: A decade and beyond. Science, 325(5939), 412–413.

Barabasi, A.-L., & Oltvai, Z. N. (2004). Network biology: Understanding the cell’s functional organization. Nature Reviews Genetics, 5(2), 101–113.

Buchfink, B., Reuter, K., & Drost, H.-G. (2021). Sensitive protein alignments at tree-of-life scale using DIAMOND. Nature Methods, 18(4), 366–368.

Cheng, F., Liu, S., Wu, J., Fang, L., Sun, S., Liu, B., Li, P., Hua, W., & Wang, X. (2011). BRAD, the genetics and genomics database for Brassica plants. BMC Plant Biology, 11, 1–6.

Csuros, M. (2010). Count: Evolutionary analysis of phylogenetic profiles with parsimony and likelihood. Bioinformatics, 26(15), 1910–1912.

Emms, D. M., & Kelly, S. (2015). OrthoFinder: Solving fundamental biases in whole genome comparisons dramatically improves orthogroup inference accuracy. Genome Biology, 16(1), 1–14.

Emms, D. M., & Kelly, S. (2019). OrthoFinder: Phylogenetic orthology inference for comparative genomics. Genome Biology, 20(1), 1–14.

Feron, R., & Waterhouse, R. M. (2022). Assessing species coverage and assembly quality of rapidly accumulating sequenced genomes. GigaScience, 11.

Goodstein, D. M., Shu, S., Howson, R., Neupane, R., Hayes, R. D., Fazo, J., Mitros, T., Dirks, W., Hellsten, U., & Putnam, N. (2012). Phytozome: A comparative platform for green plant genomics. Nucleic Acids Research, 40(D1), D1178–D1186.

Gough, J. (2005). Convergent evolution of domain architectures (is rare). Bioinformatics, 21(8), 1464–1471.

Guo, Y.-L. (2013). Gene family evolution in green plants with emphasis on the origination and evolution of a rabidopsis thaliana genes. The Plant Journal, 73(6), 941–951.

Hahn, M. W., De Bie, T., Stajich, J. E., Nguyen, C., & Cristianini, N. (2005). Estimating the tempo and mode of gene family evolution from comparative genomic data. Genome Research, 15(8), 1153–1160.

Hernández-Plaza, A., Szklarczyk, D., Botas, J., Cantalapiedra, C. P., Giner-Lamia, J., Mende, D. R., Kirsch, R., Rattei, T., Letunic, I., & Jensen, L. J. (2023). eggNOG 6.0: Enabling comparative genomics across 12 535 organisms. Nucleic Acids Research, 51(D1), D389–D394.

Huelsmann, M., Hecker, N., Springer, M. S., Gatesy, J., Sharma, V., & Hiller, M. (2019). Genes lost during the transition from land to water in cetaceans highlight genomic changes associated with aquatic adaptations. Science Advances, 5(9), eaaw6671.

Konstantinidis, K. T., & Tiedje, J. M. (2005). Genomic insights that advance the species definition for prokaryotes. Proceedings of the National Academy of Sciences, 102(7), 2567–2572.

Koonin, E. V. (2003). Comparative genomics, minimal gene-sets and the last universal common ancestor. Nature Reviews Microbiology, 1(2), 127–136.

Kuznetsov, D., Tegenfeldt, F., Manni, M., Seppey, M., Berkeley, M., Kriventseva, E. V., & Zdobnov, E. M. (2023). OrthoDB v11: Annotation of orthologs in the widest sampling of organismal diversity. Nucleic Acids Research, 51(D1), D445–D451.

Li, L., Liu, Z., Zhou, Z., Zhang, M., Meng, D., Liu, X., Huang, Y., Li, X., Jiang, Z., & Zhong, S. (2021). Comparative genomics provides insights into the genetic diversity and evolution of the DPANN superphylum. Msystems, 6(4), e00602–21.

Liu, D., Hunt, M., & Tsai, I. J. (2018). Inferring synteny between genome assemblies: A systematic evaluation. BMC Bioinformatics, 19(1), 1–13.

Lyons, E., Pedersen, B., Kane, J., Alam, M., Ming, R., Tang, H., Wang, X., Bowers, J., Paterson, A., & Lisch, D. (2008). Finding and comparing syntenic regions among Arabidopsis and the outgroups papaya, poplar, and grape: CoGe with rosids. Plant Physiology, 148(4), 1772–1781.

Marks, R. A., Hotaling, S., Frandsen, P. B., & VanBuren, R. (2021). Representation and participation across 20 years of plant genome sequencing. Nature Plants, 7(12), 1571–1578.

Mendes, F. K., Vanderpool, D., Fulton, B., & Hahn, M. W. (2020). CAFE 5 models variation in evolutionary rates among gene families. Bioinformatics, 36(22–23), 5516–5518.

Michael, T. P. (2014). Plant genome size variation: Bloating and purging DNA. Briefings in Functional Genomics, 13(4), 308–317.

Ngou, B. P. M., Heal, R., Wyler, M., Schmid, M. W., & Jones, J. D. (2022). Concerted expansion and contraction of immune receptor gene repertoires in plant genomes. Nature Plants, 8(10), 1146–1152.

Palfalvi, G., Hackl, T., Terhoeven, N., Shibata, T. F., Nishiyama, T., Ankenbrand, M., Becker, D., Förster, F., Freund, M., & Iosip, A. (2020). Genomes of the venus flytrap and close relatives unveil the roots of plant carnivory. Current Biology, 30(12), 2312–2320.

Penel, S., Arigon, A.-M., Dufayard, J.-F., Sertier, A.-S., Daubin, V., Duret, L., Gouy, M., & Perrière, G. (2009). Databases of homologous gene families for comparative genomics. BMC Bioinformatics, 10(6), 1–13.

Prüfer, K., Racimo, F., Patterson, N., Jay, F., Sankararaman, S., Sawyer, S., Heinze, A., Renaud, G., Sudmant, P. H., & De Filippo, C. (2014). The complete genome sequence of a Neanderthal from the Altai Mountains. Nature, 505(7481), 43–49.

Ravasz, E., Somera, A. L., Mongru, D. A., Oltvai, Z. N., & Barabási, A.-L. (2002). Hierarchical organization of modularity in metabolic networks. Science, 297(5586), 1551–1555.

Simão, F. A., Waterhouse, R. M., Ioannidis, P., Kriventseva, E. V., & Zdobnov, E. M. (2015). aBUSCO: Assessing genome assembly and annotation completeness with single-copy orthologs. Bioinformatics, 31(19), 3210–3212.

Sullivan, G. M., & Feinn, R. (2012). Using effect size—Or why the P value is not enough. Journal of Graduate Medical Education, 4(3), 279–282.

Sun, Y., Shang, L., Zhu, Q.-H., Fan, L., & Guo, L. (2021). Twenty years of plant genome sequencing: Achievements and challenges. Trends in Plant Science.

Van Bel, M., Silvestri, F., Weitz, E. M., Kreft, L., Botzki, A., Coppens, F., & Vandepoele, K. (2022). PLAZA 5.0: Extending the scope and power of comparative and functional genomics in plants. Nucleic Acids Research, 50(D1), 1468–1474.

Van de Peer, Y., Mizrachi, E., & Marchal, K. (2017). The evolutionary significance of polyploidy. Nature Reviews Genetics, 18(7), 411–424. https://doi.org/10.1038/nrg.2017.26

Venancio, T. M., Balaji, S., Iyer, L. M., & Aravind, L. (2009). Reconstructing the ubiquitin network-cross-talk with other systems and identification of novel functions. Genome Biology, 10, 1–18.

Wan, T., Liu, Z., Leitch, I. J., Xin, H., Maggs-Kölling, G., Gong, Y., Li, Z., Marais, E., Liao, Y., Dai, C., Liu, F., Wu, Q., Song, C., Zhou, Y., Huang, W., Jiang, K., Wang, Q., Yang, Y., Zhong, Z., … Wang, Q. (2021). The Welwitschia genome reveals a unique biology underpinning extreme longevity in deserts. Nature Communications, 12(1). https://doi.org/10.1038/s41467-021-24528-4

Wang, P., Moore, B. M., Panchy, N. L., Meng, F., Lehti-Shiu, M. D., & Shiu, S.-H. (2018). Factors influencing gene family size variation among related species in a plant family, Solanaceae. Genome Biology and Evolution, 10(10), 2596–2613.

Wang, P., & Wang, F. (2022). A proposed metric set for evaluation of genome assembly quality. Trends in Genetics.

Wang, Y., Tang, H., Debarry, J. D., Tan, X., Li, J., Wang, X., Lee, T. H., Jin, H., Marler, B., Guo, H., Kissinger, J. C., & Paterson, A. H. (2012). MCScanX: A toolkit for detection and evolutionary analysis of gene synteny and collinearity. Nucleic Acids Research, 40(7), 1–14. https://doi.org/10.1093/nar/gkr1293

Wu, M., Kostyun, J. L., Hahn, M. W., & Moyle, L. C. (2018). Dissecting the basis of novel trait evolution in a radiation with widespread phylogenetic discordance. Molecular Ecology, 27(16), 3301–3316.

Wu, T., Hu, E., Xu, S., Chen, M., Guo, P., Dai, Z., Feng, T., Zhou, L., Tang, W., & Zhan, L. (2021). clusterProfiler 4.0: A universal enrichment tool for interpreting omics data. The Innovation, 2(3), 100141.

Xu, Y., Lei, Y., Su, Z., Zhao, M., Zhang, J., Shen, G., Wang, L., Li, J., Qi, J., & Wu, J. (2021). A chromosome-scale Gastrodia elata genome and large-scale comparative genomic analysis indicate convergent evolution by gene loss in mycoheterotrophic and parasitic plants. The Plant Journal, 108(6), 1609–1623.

Yousaf, A., Liu, J., Ye, S., & Chen, H. (2021). Current Progress in Evolutionary Comparative Genomics of Great Apes. Frontiers in Genetics, 12, 657468.

Yuan, Y., Zhang, Y., Zhang, P., Liu, C., Wang, J., Gao, H., Hoelzel, A. R., Seim, I., Lv, M., & Lin, M. (2021). Comparative genomics provides insights into the aquatic adaptations of mammals. Proceedings of the National Academy of Sciences, 118(37), e2106080118.

Zhao, T., & Schranz, M. E. (2019). Network-based microsynteny analysis identifies major differences and genomic outliers in mammalian and angiosperm genomes. Proceedings of the National Academy of Sciences, 116(6), 2165–2174. https://doi.org/10.1073/pnas.1801757116

Zwaenepoel, A., & Van de Peer, Y. (2020). Model-based detection of whole-genome duplications in a phylogeny. Molecular Biology and Evolution, 37(9), 2734–2746.

