## Supplementary Text S1 for "Assessing the quality of comparative genomics data and results with the *cogeqc* R/Bioconductor package"

### Supplementary Text S1: Assessing the completeness of Chlorophyta genomes

***Fabricio Almeida-Silva*** [1,2] ***and Yves Van de Peer*** [1,2,3,4]

[1]VIB-UGent Center for Plant Systems Biology, Ghent, Belgium
[2]Department of Plant Biotechnology and Bioinformatics, Ghent University, Ghent, Belgium
[3]College of Horticulture, Academy for Advanced Interdisciplinary Studies, Nanjing Agricultural University, Nanjing, China
[4]Center for Microbial Ecology and Genomics, Department of Biochemistry, Genetics and Microbiology, University of Pretoria, Pretoria, South Africa

**2 February 2023**

```
library(here)
library(cogeqc)
library(tidyverse)
library(Herper)

set.seed(123) # for reproducibility
options(timeout = 6000) # to load files from the web
```

### 1 Overview

Here, we will use *cogeqc* to assess the completeness of Chlorophyta genomes available on Pico-PLAZA 3.0 (Van Bel et al. 2018) using Best Universal Single-Copy Orthologs (BUSCOs).

### 2 Managing external dependencies with virtual environments

Here, for convenience, we will install BUSCO in a Conda environment for use with *cogeqc*. For that, we will use the Bioconductor package *Herper*

Below, you can find the code to install miniconda in a directory of your choice (here, "~/Documents") and create a virtual environment containing a BUSCO installation.

```
# Path to where BUSCO will be installed and env name
my_miniconda <- file.path("~/Documents", "miniconda")
env <- "cogeqc_env"

# Create env named `cogeqc_env` with BUSCO in it
install_CondaTools(
    tools = "busco==5.3.0",
    env = env,
    channels = c("conda-forge", "bioconda"),
    pathToMiniConda = my_miniconda
)
```

### 3 Data acquisition

Now, we will load all genomes directly from PLAZA as `DNAStringSet` objects and export them to a single directory of FASTA files, so we can run BUSCO in batch mode.

```
# Links to Chlorophyta genomes from Pico-PLAZA 3.0
base_url <- "ftp://ftp.psb.ugent.be/pub/plaza/plaza_pico_03/Genomes/"
links <- paste0(
    base_url,
    c("mpu.fasta.gz", "mrcc299.fasta.gz", "olu.fasta.gz", "ome.fasta.gz",
      "orcc809.fasta.gz", "ota.fasta.gz", "bprrcc1105.fasta.gz",
      "cre.fasta.gz", "vca.fasta.gz", "cvu.fasta.gz", "acg.fasta.gz",
      "pse3.fasta.gz", "prcc4223.fasta.gz", "cnc64a.fasta.gz",
```

```
        "hsp.fasta.gz", "apr.fasta.gz")
)

# Load all genomes
genomes <- lapply(links, Biostrings::readDNAStringSet)
names(genomes) <- basename(links)

# Write all genomes to a subdirectory of tempdir
genomes_path <- file.path(tempdir(), "genomes")
if(!dir.exists(genomes_path)) { fs::dir_create(genomes_path) }

write <- lapply(seq_along(genomes), function(x) {
    Biostrings::writeXStringSet(
        x = genomes[[x]],
        filepath = file.path(genomes_path, names(genomes)[x])
    )
    return(NULL)
})

dir(genomes_path)
##  [1] "acg.fasta.gz"        "apr.fasta.gz"        "bprrcc1105.fasta.gz"
##  [4] "cnc64a.fasta.gz"     "cre.fasta.gz"        "cvu.fasta.gz"
##  [7] "hsp.fasta.gz"        "mpu.fasta.gz"        "mrcc299.fasta.gz"
## [10] "olu.fasta.gz"        "ome.fasta.gz"        "orcc809.fasta.gz"
## [13] "ota.fasta.gz"        "prcc4223.fasta.gz"   "pse3.fasta.gz"
## [16] "vca.fasta.gz"
```

### 4 Running BUSCO

Now that all genomes are stored as FASTA files in /tmp/RtmpcuLMqJ/genomes, we can assess their completeness with BUSCO.

```
# See all possible lineage datasets
with_CondaEnv(
    env, list_busco_datasets(), my_miniconda
)

# Run BUSCO using chlorophyta_odb10 as the lineage data set
busco <- with_CondaEnv(
    env,
    run_busco(
        sequence = genomes_path,
        outlabel = "chlorophyta_busco",
        mode = "genome",
        lineage = "chlorophyta_odb10",
        outpath = tempdir(),
        download_path = tempdir()
    ),
    my_miniconda
)
```

```r
# Read and parse the output
outdir <- file.path(tempdir(), "chlorophyta_busco")
busco_summary <- read_busco(outdir)
save(
    busco_summary,
    file = here::here("products", "result_files", "busco_summary.rda"),
    compress = "xz"
)
```

The parsed BUSCO output (as returned by `read_busco()`) looks like this:

```r
load(here("products", "result_files", "busco_summary.rda"))
head(busco_summary)
##          Class Frequency           Lineage           File
## 1 Complete_SC      94.1 chlorophyta_odb10    pse3.fasta.gz
## 2 Complete_SC      95.1 chlorophyta_odb10     cre.fasta.gz
## 3 Complete_SC      96.8 chlorophyta_odb10     olu.fasta.gz
## 4 Complete_SC      98.7 chlorophyta_odb10 mrcc299.fasta.gz
## 5 Complete_SC      91.8 chlorophyta_odb10     apr.fasta.gz
## 6 Complete_SC      86.4 chlorophyta_odb10     acg.fasta.gz
```

### 5 Visualizing summary statistics

Finally, let's visualize summary BUSCO stats:

```r
# Manually create tree based on Pico-PLAZA's tree
c_branches <- function(b1, b2) {
    x <- paste0("(", b1, ",", b2, ")")
}

ostreococcus_root <- "((((Ostreococcus_lucimarinus, Ostreococcus_sp_RCC809), Ostreococcus_tauri), Ostreococcu
micromonas <- "(Micromonas_pusilla_strain_CCMP1545, Micromonas_sp_RCC299)"
chlamydomonadales <- "(Volvox_carteri, Chlamydomonas_reinhardtii)"
picochlorum <- "(Picochlorum_sp_SENEW3, Picochlorum_RCC4223)"
chlorellales <- "((Helicosporidium_sp, Auxenochlorella_protothecoides), Chlorella_sp_NC64A)"
trebouxiophyceae <- c_branches(
    "(Coccomyxa_subellipsoidea_C-169, Asterochloris_sp_Cgr/DA1pho_v2)",
    c_branches(picochlorum, chlorellales)
)

chlo_tree <- c_branches(
    c_branches(
        ostreococcus_root, micromonas
    ),
    c_branches(
        chlamydomonadales, trebouxiophyceae
    )
)
chlo_tree <- paste0(chlo_tree, ";")
```

```r
# Read tree as a phylo object and clean species names
chlo_tree <- treeio::read.tree(text = chlo_tree)
chlo_tree$tip.label <- gsub("_", " ", chlo_tree$tip.label)

# Plot species tree and get species order from tree topology
p_tree <- plot_species_tree(chlo_tree, xlim = c(0, 12))
taxa_order <- rev(ggtree::get_taxa_name(p_tree))

# Plot BUSCO summary stats
p_busco <- busco_summary %>%
    mutate(File = str_replace_all(File, "\\.fasta.*", "")) %>%
    mutate(File = str_replace_all(
        File,
        c(
            "pse3" = "Picochlorum sp SENEW3",
            "cre" = "Chlamydomonas reinhardtii",
            "olu" = "Ostreococcus lucimarinus",
            "mrcc299" = "Micromonas sp RCC299",
            "apr" = "Auxenochlorella protothecoides",
            "acg" = "Asterochloris sp Cgr/DA1pho v2",
            "cvu" = "Coccomyxa subellipsoidea C-169",
            "bprrcc1105" = "Bathycoccus prasinos",
            "orcc809" = "Ostreococcus sp RCC809",
            "prcc4223" = "Picochlorum RCC4223",
            "ota" = "Ostreococcus tauri",
            "hsp" = "Helicosporidium sp",
            "mpu" = "Micromonas pusilla strain CCMP1545",
            "vca" = "Volvox carteri",
            "ome" = "Ostreococcus mediterraneus",
            "cnc64a" = "Chlorella sp NC64A"
        )
    )) %>%
    mutate(File = factor(File, taxa_order)) %>%
    plot_busco() +
    theme(axis.text.y = element_blank()) +
    labs(y = "")

# Combining phylogeny with BUSCO plot
combined <- patchwork::wrap_plots(p_tree, p_busco)
combined
```

Except for *Helicosporidium sp.*, Chlorophyta genomes on Pico-PLAZA 3.0 have a high quality, as denoted by their high completeness.

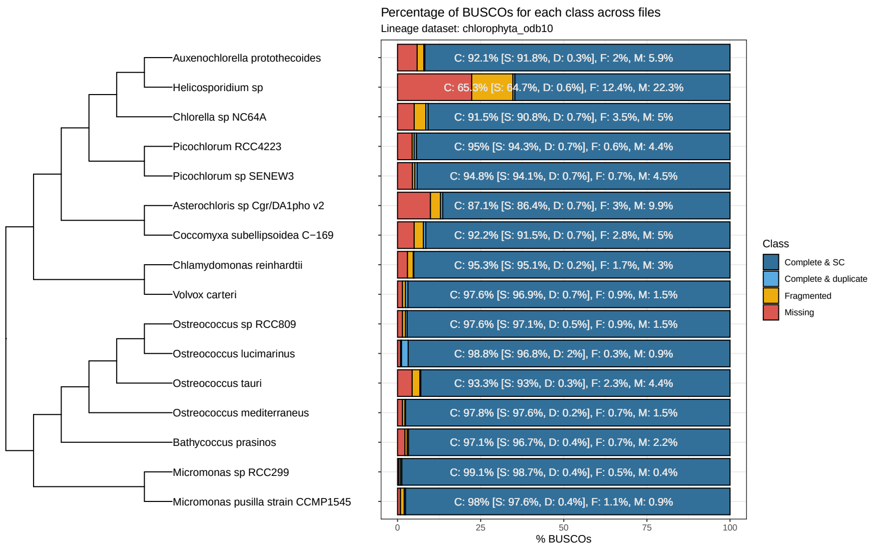

**Figure 1:** BUSCO scores for Chlorophyta genomes on Pico-PLAZA 3.0.

### Session info

This document was created under the following conditions:

```
sessioninfo::session_info()
## - Session info ---------------------------------------------------------------
##  setting  value
##  version  R version 4.2.2 Patched (2022-11-10 r83330)
##  os       Ubuntu 20.04.5 LTS
##  system   x86_64, linux-gnu
##  ui       X11
##  language (EN)
##  collate  en_US.UTF-8
##  ctype    en_US.UTF-8
##  tz       Europe/Brussels
##  date     2023-02-02
##  pandoc   2.19.2 @ /usr/lib/rstudio/resources/app/bin/quarto/bin/tools/ (via rmarkdown)
##
## - Packages -------------------------------------------------------------------
##  package          * version   date (UTC) lib source
##  ape                5.6-2     2022-03-02 [1] CRAN (R 4.2.0)
##  aplot              0.1.8     2022-10-09 [1] CRAN (R 4.2.1)
##  assertthat         0.2.1     2019-03-21 [1] CRAN (R 4.2.0)
##  backports          1.4.1     2021-12-13 [1] CRAN (R 4.2.0)
##  beeswarm           0.4.0     2021-06-01 [1] CRAN (R 4.2.2)
##  BiocGenerics       0.42.0    2022-04-26 [1] Bioconductor
##  BiocManager        1.30.18   2022-05-18 [1] CRAN (R 4.2.0)
##  BiocStyle        * 2.25.0    2022-06-15 [1] Github (Bioconductor/BiocStyle@7150c28)
##  Biostrings         2.64.1    2022-08-18 [1] Bioconductor
##  bitops             1.0-7     2021-04-24 [1] CRAN (R 4.2.0)
```

```
## bookdown           0.29      2022-09-12 [1] CRAN (R 4.2.1)
## broom              1.0.1     2022-08-29 [1] CRAN (R 4.2.1)
## cellranger         1.1.0     2016-07-27 [1] CRAN (R 4.2.0)
## cli                3.4.1     2022-09-23 [1] CRAN (R 4.2.1)
## cogeqc           * 1.3.1     2023-01-24 [1] Bioconductor
## colorspace         2.0-3     2022-02-21 [1] CRAN (R 4.2.0)
## crayon             1.5.2     2022-09-29 [1] CRAN (R 4.2.1)
## DBI                1.1.3     2022-06-18 [1] CRAN (R 4.2.0)
## dbplyr             2.2.1     2022-06-27 [1] CRAN (R 4.2.1)
## digest             0.6.29    2021-12-01 [1] CRAN (R 4.2.0)
## dplyr            * 1.0.10    2022-09-01 [1] CRAN (R 4.2.1)
## ellipsis           0.3.2     2021-04-29 [1] CRAN (R 4.2.0)
## evaluate           0.17      2022-10-07 [1] CRAN (R 4.2.1)
## fansi              1.0.3     2022-03-24 [1] CRAN (R 4.2.0)
## farver             2.1.1     2022-07-06 [1] CRAN (R 4.2.1)
## fastmap            1.1.0     2021-01-25 [1] CRAN (R 4.2.0)
## forcats          * 0.5.2     2022-08-19 [1] CRAN (R 4.2.1)
## fs                 1.5.2     2021-12-08 [1] CRAN (R 4.2.0)
## gargle             1.2.1     2022-09-08 [1] CRAN (R 4.2.1)
## generics           0.1.3     2022-07-05 [1] CRAN (R 4.2.1)
## GenomeInfoDb       1.32.4    2022-09-06 [1] Bioconductor
## GenomeInfoDbData   1.2.8     2022-05-06 [1] Bioconductor
## ggbeeswarm         0.7.1     2022-12-16 [1] CRAN (R 4.2.2)
## ggfun              0.0.8     2022-11-07 [1] CRAN (R 4.2.1)
## ggplot2          * 3.4.0     2022-11-04 [1] CRAN (R 4.2.1)
## ggplotify          0.1.0     2021-09-02 [1] CRAN (R 4.2.0)
## ggtree             3.7.1.001 2022-11-10 [1] Github (YuLab-SMU/ggtree@b7ef83e)
## glue               1.6.2     2022-02-24 [1] CRAN (R 4.2.0)
## googledrive        2.0.0     2021-07-08 [1] CRAN (R 4.2.0)
## googlesheets4      1.0.1     2022-08-13 [1] CRAN (R 4.2.1)
## gridGraphics       0.5-1     2020-12-13 [1] CRAN (R 4.2.0)
## gtable             0.3.1     2022-09-01 [1] CRAN (R 4.2.1)
## haven              2.5.1     2022-08-22 [1] CRAN (R 4.2.1)
## here             * 1.0.1     2020-12-13 [1] CRAN (R 4.2.0)
## Herper           * 1.1.2     2022-05-18 [1] Github (RockefellerUniversity/Herper@5bceeb4)
## hms                1.1.2     2022-08-19 [1] CRAN (R 4.2.1)
## htmltools          0.5.3     2022-07-18 [1] CRAN (R 4.2.1)
## httr               1.4.4     2022-08-17 [1] CRAN (R 4.2.1)
## igraph             1.3.5     2022-09-22 [1] CRAN (R 4.2.1)
## IRanges            2.30.1    2022-08-18 [1] Bioconductor
## jsonlite           1.8.3     2022-10-21 [1] CRAN (R 4.2.1)
## knitr              1.40      2022-08-24 [1] CRAN (R 4.2.1)
## labeling           0.4.2     2020-10-20 [1] CRAN (R 4.2.0)
## lattice            0.20-45   2021-09-22 [1] CRAN (R 4.2.0)
## lazyeval           0.2.2     2019-03-15 [1] CRAN (R 4.2.0)
## lifecycle          1.0.3     2022-10-07 [1] CRAN (R 4.2.1)
## lubridate          1.8.0     2021-10-07 [1] CRAN (R 4.2.0)
## magrittr           2.0.3     2022-03-30 [1] CRAN (R 4.2.0)
## Matrix             1.5-1     2022-09-13 [1] CRAN (R 4.2.1)
## modelr             0.1.9     2022-08-19 [1] CRAN (R 4.2.1)
## munsell            0.5.0     2018-06-12 [1] CRAN (R 4.2.0)
```

```
##  nlme               3.1-160   2022-10-10 [1] CRAN (R 4.2.1)
##  patchwork          1.1.2     2022-08-19 [1] CRAN (R 4.2.1)
##  pillar             1.8.1     2022-08-19 [1] CRAN (R 4.2.1)
##  pkgconfig          2.0.3     2019-09-22 [1] CRAN (R 4.2.0)
##  plyr               1.8.7     2022-03-24 [1] CRAN (R 4.2.0)
##  png                0.1-7     2013-12-03 [1] CRAN (R 4.2.0)
##  purrr            * 0.3.5     2022-10-06 [1] CRAN (R 4.2.1)
##  R6                 2.5.1     2021-08-19 [1] CRAN (R 4.2.0)
##  Rcpp               1.0.9     2022-07-08 [1] CRAN (R 4.2.1)
##  RCurl              1.98-1.9  2022-10-03 [1] CRAN (R 4.2.1)
##  readr            * 2.1.3     2022-10-01 [1] CRAN (R 4.2.1)
##  readxl             1.4.1     2022-08-17 [1] CRAN (R 4.2.1)
##  reprex             2.0.2     2022-08-17 [1] CRAN (R 4.2.1)
##  reshape2           1.4.4     2020-04-09 [1] CRAN (R 4.2.0)
##  reticulate       * 1.26      2022-08-31 [1] CRAN (R 4.2.1)
##  rjson              0.2.21    2022-01-09 [1] CRAN (R 4.2.0)
##  rlang              1.0.6     2022-09-24 [1] CRAN (R 4.2.1)
##  rmarkdown          2.17      2022-10-07 [1] CRAN (R 4.2.1)
##  rprojroot          2.0.3     2022-04-02 [1] CRAN (R 4.2.0)
##  rstudioapi         0.14      2022-08-22 [1] CRAN (R 4.2.1)
##  rvest              1.0.3     2022-08-19 [1] CRAN (R 4.2.1)
##  S4Vectors          0.34.0    2022-04-26 [1] Bioconductor
##  scales             1.2.1     2022-08-20 [1] CRAN (R 4.2.1)
##  sessioninfo        1.2.2     2021-12-06 [1] CRAN (R 4.2.0)
##  stringi            1.7.8     2022-07-11 [1] CRAN (R 4.2.1)
##  stringr          * 1.4.1     2022-08-20 [1] CRAN (R 4.2.1)
##  tibble           * 3.1.8     2022-07-22 [1] CRAN (R 4.2.1)
##  tidyr            * 1.2.1     2022-09-08 [1] CRAN (R 4.2.1)
##  tidyselect         1.2.0     2022-10-10 [1] CRAN (R 4.2.1)
##  tidytree           0.4.1     2022-09-26 [1] CRAN (R 4.2.1)
##  tidyverse        * 1.3.2     2022-07-18 [1] CRAN (R 4.2.1)
##  treeio             1.23.0    2022-11-10 [1] Github (GuangchuangYu/treeio@db85803)
##  tzdb               0.3.0     2022-03-28 [1] CRAN (R 4.2.0)
##  utf8               1.2.2     2021-07-24 [1] CRAN (R 4.2.0)
##  vctrs              0.5.0     2022-10-22 [1] CRAN (R 4.2.1)
##  vipor              0.4.5     2017-03-22 [1] CRAN (R 4.2.1)
##  withr              2.5.0     2022-03-03 [1] CRAN (R 4.2.0)
##  xfun               0.33      2022-09-12 [1] CRAN (R 4.2.1)
##  xml2               1.3.3     2021-11-30 [1] CRAN (R 4.2.0)
##  XVector            0.36.0    2022-04-26 [1] Bioconductor
##  yaml               2.3.5     2022-02-21 [1] CRAN (R 4.2.0)
##  yulab.utils        0.0.5     2022-06-30 [1] CRAN (R 4.2.1)
##  zlibbioc           1.42.0    2022-04-26 [1] Bioconductor
##
##  [1] /home/faalm/R/x86_64-pc-linux-gnu-library/4.2
##  [2] /usr/local/lib/R/site-library
##  [3] /usr/lib/R/site-library
##  [4] /usr/lib/R/library
##
## ------------------------------------------------------------------------------
```

### References

Van Bel, Michiel, Tim Diels, Emmelien Vancaester, Lukasz Kreft, Alexander Botzki, Yves Van de Peer, Frederik Coppens, and Klaas Vandepoele. 2018. "PLAZA 4.0: An Integrative Resource for Functional, Evolutionary and Comparative Plant Genomics." *Nucleic Acids Research* 46 (D1): D1190–96.
