## Supplementary Text S2 for "Assessing the quality of comparative genomics data and results with the *cogeqc* R/Bioconductor package"

### Supplementary Text S2: Assessing orthogroup inference in public databases

***Fabricio Almeida-Silva*** [1,2] ***and Yves Van de Peer*** [1,2,3,4]

[1]VIB-UGent Center for Plant Systems Biology, Ghent, Belgium
[2]Department of Plant Biotechnology and Bioinformatics, Ghent University, Ghent, Belgium
[3]College of Horticulture, Academy for Advanced Interdisciplinary Studies, Nanjing Agricultural University, Nanjing, China
[4]Center for Microbial Ecology and Genomics, Department of Biochemistry, Genetics and Microbiology, University of Pretoria, Pretoria, South Africa

**2 February 2023**

```
library(cogeqc)
library(here)
library(tidyverse)
library(ggpubr)

set.seed(123) # for reproducibility
source(here("code", "utils.R"))
```

### 1 Overview

Here, we will use the protein domain-based approach in *cogeqc* to assess gene families from different sources, namely:

- PLAZA Dicots 5.0 (Van Bel et al. 2022)
- OrthoDB (Kuznetsov et al. 2023)
- eggNOG (Hernandez-Plaza et al. 2023)
- HOGENOM (Penel et al. 2009)

### 2 Calculating orthogroup scores

To make comparison possible, we will *Arabidopsis thaliana* domain annotation as a proxy, as this species is present in all of the aforementioned databases. For that, we will use the function `calculate_H()` from *cogeqc*.

Orthogroups assignments from OrthoDB, eggNOG, InParanoid, PhylomeDB, and HOGENOM will be obtained from UniProt.

#### 2.1 PLAZA Dicots 5.0

Below, we will obtain orthogroups and *A. thaliana*'s domain annotation from PLAZA 5.0, and then we will calculate homogeneity scores for each orthogroup.

```
# Obtain gene families from PLAZA
fams_plaza <- readr::read_tsv(
    paste0(
        "https://ftp.psb.ugent.be/pub/plaza/plaza_public_dicots_05/",
        "GeneFamilies/genefamily_data.HOMFAM.csv.gz"
    ), show_col_types = FALSE, skip = 2
) %>%
    filter(species == "ath") %>%
    as.data.frame()
names(fams_plaza) <- c("Orthogroup", "Species", "Gene")
head(fams_plaza)
##     Orthogroup Species      Gene
## 1 HOM05D000001     ath AT1G02310
## 2 HOM05D000001     ath AT1G03510
## 3 HOM05D000001     ath AT1G03540
## 4 HOM05D000001     ath AT1G04020
## 5 HOM05D000001     ath AT1G04840
```

```
## 6 HOM05D000001     ath AT1G05750

# Obtain domain anotation for A. thaliana
ath_interpro <- readr::read_tsv(
    paste0(
        "https://ftp.psb.ugent.be/pub/plaza/plaza_public_dicots_05/",
        "InterPro/interpro.ath.csv.gz"
    ), show_col_types = FALSE, skip = 8
) %>%
    select(1,3)
names(ath_interpro) <- c("Gene", "Annotation")
head(ath_interpro)
## # A tibble: 6 x 2
##   Gene      Annotation
##   <chr>     <chr>
## 1 AT1G01010 IPR036093
## 2 AT1G01010 IPR003441
## 3 AT1G01010 IPR036093
## 4 AT1G01020 IPR007290
## 5 AT1G01020 IPR007290
## 6 AT1G01030 IPR003340

# Combining everything and calculating homogeneity scores
fam_df_plaza <- merge(fams_plaza, ath_interpro)
head(fam_df_plaza)
##        Gene   Orthogroup Species Annotation
## 1 AT1G01010 HOM05D000010     ath  IPR036093
## 2 AT1G01010 HOM05D000010     ath  IPR003441
## 3 AT1G01010 HOM05D000010     ath  IPR036093
## 4 AT1G01020 HOM05D006082     ath  IPR007290
## 5 AT1G01020 HOM05D006082     ath  IPR007290
## 6 AT1G01030 HOM05D000466     ath  IPR015300

H_summary <- function(ortho_df = NULL) {
    H <- calculate_H(ortho_df)
    mean_H <- round(mean(H$Score), 2)
    median_H <- round(median(H$Score), 2)
    result_list <- list(H = H, mean_score = mean_H, median_score = median_H)
    return(result_list)
}

H_plaza <- H_summary(fam_df_plaza)
head(H_plaza$H)
##     Orthogroup     Score
## 1 HOM05D000001  283.3132
## 2 HOM05D000002  129.9598
## 3 HOM05D000003  889.1268
## 4 HOM05D000004    0.0000
## 5 HOM05D000005 1135.8799
## 6 HOM05D000006 2820.8337
```

#### 2.2 OrthoDB, eggNOG, and HOGENOM

Orthogroup assignments from these databases will be obtained from UniProt (Consortium 2021).

```
# Get list of proteins - from primary transcripts only
ath_proteome <- Biostrings::readAAStringSet(
    paste0(
        "https://ftp.uniprot.org/pub/databases/uniprot/",
        "current_release/knowledgebase/reference_proteomes/Eukaryota/",
        "UP000006548/UP000006548_3702.fasta.gz"
    )
)
ath_proteins <- names(ath_proteome)
ath_proteins <- sapply(strsplit(ath_proteins, split = "\\|"), `[`, 2)

# Extract phylogenomic information for all genes
source(here::here("code", "utils.R"))
fams_uniprot <- extract_ogs_uniprot(ath_proteins)

fams_orthodb <- fams_uniprot[, c("Gene", "OrthoDB")] %>% drop_na()
fams_eggnog <- fams_uniprot[, c("Gene", "eggNOG")] %>% drop_na()
fams_hogenom <- fams_uniprot[, c("Gene", "HOGENOM")] %>% drop_na()

#----Calculate homogeneity scores for each database-----------------------------
# OrthoDB
fams_df_orthodb <- merge(fams_orthodb, ath_interpro)
names(fams_df_orthodb)[2] <- "Orthogroup"
H_orthodb <- H_summary(fams_df_orthodb)

# eggNOG
fams_df_eggnog <- merge(fams_eggnog, ath_interpro)
names(fams_df_eggnog)[2] <- "Orthogroup"
H_eggnog <- H_summary(fams_df_eggnog)

# HOGENOM
fams_df_hogenom <- merge(fams_hogenom, ath_interpro)
names(fams_df_hogenom)[2] <- "Orthogroup"
H_hogenom <- H_summary(fams_df_hogenom)
```

### 3 Comparing homogeneity scores

Finally, let's compare homogeneity scores and visualize their distributions. First, let's combine all data frames of homogeneity scores into a single data frame.

```
H_combined <- bind_rows(
    H_plaza$H %>% mutate(Source = "PLAZA"),
    H_orthodb$H %>% mutate(Source = "OrthoDB"),
    H_eggnog$H %>% mutate(Source = "eggNOG"),
    H_hogenom$H %>% mutate(Source = "HOGENOM")
)
```

```
save(
    H_combined,
    file = here::here("products", "result_files", "H_combined.rda"),
    compress = "xz"
)
```

Now, let's compare the distributions of homogeneity scores for each database to see if there are any differences. For that, we will calculate P-values from a Wilcoxon test with Wicoxon effect sizes (r). The Wilcoxon effect size is calculated as the Z statistic divided by the square root of the sample size.

```
# Scale scores to maximum, so that they range from 0 to 1
H_combined$Score <- H_combined$Score / max(H_combined$Score)
head(H_combined)
##     Orthogroup      Score Source
## 1 HOM05D000001 0.10043599  PLAZA
## 2 HOM05D000002 0.04607143  PLAZA
## 3 HOM05D000003 0.31520000  PLAZA
## 4 HOM05D000004 0.00000000  PLAZA
## 5 HOM05D000005 0.40267523  PLAZA
## 6 HOM05D000006 1.00000000  PLAZA

# Quick exploration of means and medians
H_combined %>%
    group_by(Source) %>%
    summarise(mean = mean(Score), median = median(Score))
## # A tibble: 4 x 3
##   Source   mean median
##   <chr>   <dbl>  <dbl>
## 1 eggNOG  0.565  0.546
## 2 HOGENOM 0.603  0.609
## 3 OrthoDB 0.578  0.567
## 4 PLAZA   0.610  0.6

# Compare homogeneity scores - all vs all
db_wilcox <- compare(H_combined, "Score ~ Source")

db_wilcox |>
    filter_comparison() |>
    knitr::kable(
        caption = "Mann-Whitney U test for differences in orthogroup scores with Wilcoxon effect sizes.",
        digits = 10
    )
```

We can see that there are diffences in mean. In summary:

1. eggNOG orthogroups have lower scores than every other source
2. HOGENOM orthogroups have higher scores than OrthoDB, but lower than PLAZA.
3. PLAZA orthogroup scores are higher than every other database.

**Table 1:** Mann-Whitney U test for differences in orthogroup scores with Wilcoxon effect sizes.

| group1 | group2 | n1 | n2 | padj | effsize | magnitude |
|---|---|---|---|---|---|---|
| eggNOG | HOGENOM | 3092 | 3257 | 0.0e+00 | 0.11102956 | small |
| eggNOG | OrthoDB | 3092 | 3201 | 8.5e-09 | 0.07197679 | small |
| eggNOG | PLAZA | 3092 | 3503 | 0.0e+00 | 0.09434683 | small |
| HOGENOM | OrthoDB | 3257 | 3201 | 0.0e+00 | 0.09071787 | small |
| HOGENOM | PLAZA | 3257 | 3503 | 3.0e-03 | 0.03402611 | small |
| OrthoDB | PLAZA | 3201 | 3503 | 7.0e-10 | 0.07526911 | small |

However, the effect sizes are very small, suggesting that significant differences could be due to large sample sizes, as P-values are highly affected by sample sizes.

Now, let's visualize the distributions with significant differences highlighted. Here, we will only display comparison bars for comparisons with $P < 0.05$ and effect sizes $> 0.1$.

```
# Comparisons to be made
comps <- list(
    c("HOGENOM", "eggNOG")
)

# Change order of levels according to comparison results
H_combined$Source <- factor(
    H_combined$Source, levels = rev(c(
        "PLAZA", "HOGENOM", "OrthoDB", "eggNOG"
    ))
)

# Visualize distributions with significant differences highlighted
distros <- ggviolin(
    H_combined, y = "Score", x = "Source",
    orientation = "horiz", trim = TRUE, add = c("boxplot", "mean"),
    fill = "Source", add.params = list(fill = "white"), palette = "jama"
) +
    ggpubr::stat_compare_means(
        comparisons = comps,
        label = "p.signif",
        method = "wilcox.test"
    ) +
    theme(legend.position = "none") +
    labs(y = "Scaled homogeneity scores", x = "Source of orthogroups",
         title = "Distribution of mean homogeneity scores for orthogroups",
         subtitle = "Scores were calculated based on *A. thaliana* genes") +
    theme(plot.subtitle = ggtext::element_markdown())

distros
```

To conclude, despite some significant differences, all databases perform equally well in their orthogroup definition. The observed differences in means could be due to large sample sizes, as indicated by very low effect sizes, and to the different species composition of the database.

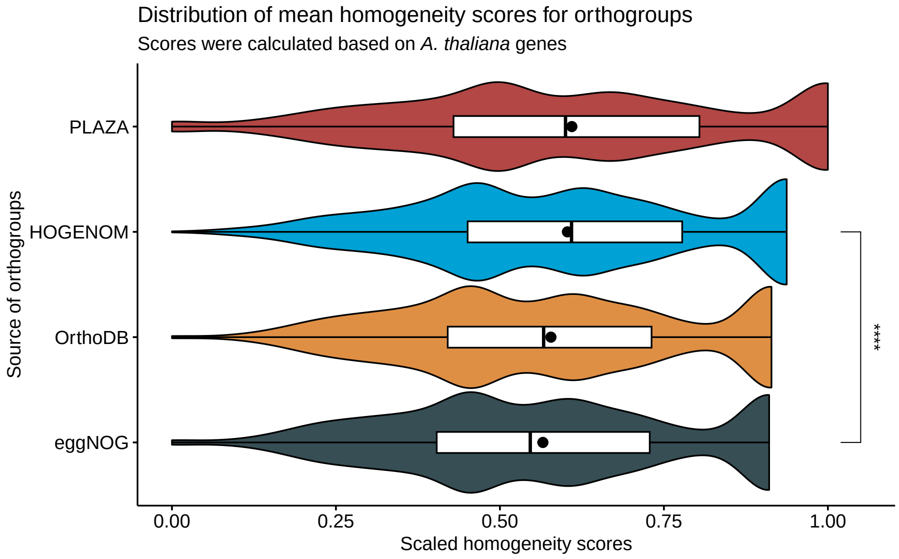

**Figure 1:** Distribution of mean orthogroup scores.

### Session info

This document was created under the following conditions:

```
sessioninfo::session_info()
## - Session info ---------------------------------------------------------------
##  setting  value
##  version  R version 4.2.2 Patched (2022-11-10 r83330)
##  os       Ubuntu 20.04.5 LTS
##  system   x86_64, linux-gnu
##  ui       X11
##  language (EN)
##  collate  en_US.UTF-8
##  ctype    en_US.UTF-8
##  tz       Europe/Brussels
##  date     2023-02-02
##  pandoc   2.19.2 @ /usr/lib/rstudio/resources/app/bin/quarto/bin/tools/ (via rmarkdown)
##
## - Packages -------------------------------------------------------------------
##  package          * version   date (UTC) lib source
##  abind              1.4-5     2016-07-21 [1] CRAN (R 4.2.0)
##  ape                5.6-2     2022-03-02 [1] CRAN (R 4.2.0)
##  aplot              0.1.8     2022-10-09 [1] CRAN (R 4.2.1)
##  assertthat         0.2.1     2019-03-21 [1] CRAN (R 4.2.0)
##  backports          1.4.1     2021-12-13 [1] CRAN (R 4.2.0)
##  beeswarm           0.4.0     2021-06-01 [1] CRAN (R 4.2.2)
##  BiocGenerics       0.42.0    2022-04-26 [1] Bioconductor
##  BiocManager        1.30.18   2022-05-18 [1] CRAN (R 4.2.0)
##  BiocStyle        * 2.25.0    2022-06-15 [1] Github (Bioconductor/BiocStyle@7150c28)
##  Biostrings         2.64.1    2022-08-18 [1] Bioconductor
```

```
##  bit                 4.0.4     2020-08-04 [1] CRAN (R 4.2.0)
##  bit64               4.0.5     2020-08-30 [1] CRAN (R 4.2.0)
##  bitops              1.0-7     2021-04-24 [1] CRAN (R 4.2.0)
##  bookdown            0.29      2022-09-12 [1] CRAN (R 4.2.1)
##  broom               1.0.1     2022-08-29 [1] CRAN (R 4.2.1)
##  car                 3.1-0     2022-06-15 [1] CRAN (R 4.2.0)
##  carData             3.0-5     2022-01-06 [1] CRAN (R 4.2.0)
##  cellranger          1.1.0     2016-07-27 [1] CRAN (R 4.2.0)
##  cli                 3.4.1     2022-09-23 [1] CRAN (R 4.2.1)
##  codetools           0.2-18    2020-11-04 [1] CRAN (R 4.2.0)
##  cogeqc            * 1.3.1     2023-01-24 [1] Bioconductor
##  coin                1.4-2     2021-10-08 [1] CRAN (R 4.2.1)
##  colorspace          2.0-3     2022-02-21 [1] CRAN (R 4.2.0)
##  crayon              1.5.2     2022-09-29 [1] CRAN (R 4.2.1)
##  curl                4.3.3     2022-10-06 [1] CRAN (R 4.2.1)
##  DBI                 1.1.3     2022-06-18 [1] CRAN (R 4.2.0)
##  dbplyr              2.2.1     2022-06-27 [1] CRAN (R 4.2.1)
##  digest              0.6.29    2021-12-01 [1] CRAN (R 4.2.0)
##  dplyr             * 1.0.10    2022-09-01 [1] CRAN (R 4.2.1)
##  ellipsis            0.3.2     2021-04-29 [1] CRAN (R 4.2.0)
##  evaluate            0.17      2022-10-07 [1] CRAN (R 4.2.1)
##  fansi               1.0.3     2022-03-24 [1] CRAN (R 4.2.0)
##  farver              2.1.1     2022-07-06 [1] CRAN (R 4.2.1)
##  fastmap             1.1.0     2021-01-25 [1] CRAN (R 4.2.0)
##  forcats           * 0.5.2     2022-08-19 [1] CRAN (R 4.2.1)
##  fs                  1.5.2     2021-12-08 [1] CRAN (R 4.2.0)
##  gargle              1.2.1     2022-09-08 [1] CRAN (R 4.2.1)
##  generics            0.1.3     2022-07-05 [1] CRAN (R 4.2.1)
##  GenomeInfoDb        1.32.4    2022-09-06 [1] Bioconductor
##  GenomeInfoDbData    1.2.8     2022-05-06 [1] Bioconductor
##  ggbeeswarm          0.7.1     2022-12-16 [1] CRAN (R 4.2.2)
##  ggfun               0.0.8     2022-11-07 [1] CRAN (R 4.2.1)
##  ggplot2           * 3.4.0     2022-11-04 [1] CRAN (R 4.2.1)
##  ggplotify           0.1.0     2021-09-02 [1] CRAN (R 4.2.0)
##  ggpubr            * 0.4.0     2020-06-27 [1] CRAN (R 4.2.0)
##  ggsci               2.9       2018-05-14 [1] CRAN (R 4.2.0)
##  ggsignif            0.6.4     2022-10-13 [1] CRAN (R 4.2.1)
##  ggtext              0.1.2     2022-09-16 [1] CRAN (R 4.2.1)
##  ggtree              3.7.1.001 2022-11-10 [1] Github (YuLab-SMU/ggtree@b7ef83e)
##  glue                1.6.2     2022-02-24 [1] CRAN (R 4.2.0)
##  googledrive         2.0.0     2021-07-08 [1] CRAN (R 4.2.0)
##  googlesheets4       1.0.1     2022-08-13 [1] CRAN (R 4.2.1)
##  gridGraphics        0.5-1     2020-12-13 [1] CRAN (R 4.2.0)
##  gridtext            0.1.5     2022-09-16 [1] CRAN (R 4.2.1)
##  gtable              0.3.1     2022-09-01 [1] CRAN (R 4.2.1)
##  haven               2.5.1     2022-08-22 [1] CRAN (R 4.2.1)
##  here              * 1.0.1     2020-12-13 [1] CRAN (R 4.2.0)
##  hms                 1.1.2     2022-08-19 [1] CRAN (R 4.2.1)
##  htmltools           0.5.3     2022-07-18 [1] CRAN (R 4.2.1)
##  httr                1.4.4     2022-08-17 [1] CRAN (R 4.2.1)
##  igraph              1.3.5     2022-09-22 [1] CRAN (R 4.2.1)
```

```
##  IRanges            2.30.1    2022-08-18 [1] Bioconductor
##  jsonlite           1.8.3     2022-10-21 [1] CRAN (R 4.2.1)
##  knitr              1.40      2022-08-24 [1] CRAN (R 4.2.1)
##  labeling           0.4.2     2020-10-20 [1] CRAN (R 4.2.0)
##  lattice            0.20-45   2021-09-22 [1] CRAN (R 4.2.0)
##  lazyeval           0.2.2     2019-03-15 [1] CRAN (R 4.2.0)
##  libcoin            1.0-9     2021-09-27 [1] CRAN (R 4.2.1)
##  lifecycle          1.0.3     2022-10-07 [1] CRAN (R 4.2.1)
##  lubridate          1.8.0     2021-10-07 [1] CRAN (R 4.2.0)
##  magrittr           2.0.3     2022-03-30 [1] CRAN (R 4.2.0)
##  markdown           1.1       2019-08-07 [1] CRAN (R 4.2.0)
##  MASS               7.3-58.1  2022-08-03 [1] CRAN (R 4.2.1)
##  Matrix             1.5-1     2022-09-13 [1] CRAN (R 4.2.1)
##  matrixStats        0.62.0    2022-04-19 [1] CRAN (R 4.2.0)
##  modelr             0.1.9     2022-08-19 [1] CRAN (R 4.2.1)
##  modeltools         0.2-23    2020-03-05 [1] CRAN (R 4.2.1)
##  multcomp           1.4-20    2022-08-07 [1] CRAN (R 4.2.1)
##  munsell            0.5.0     2018-06-12 [1] CRAN (R 4.2.0)
##  mvtnorm            1.1-3     2021-10-08 [1] CRAN (R 4.2.0)
##  nlme               3.1-160   2022-10-10 [1] CRAN (R 4.2.1)
##  patchwork          1.1.2     2022-08-19 [1] CRAN (R 4.2.1)
##  pillar             1.8.1     2022-08-19 [1] CRAN (R 4.2.1)
##  pkgconfig          2.0.3     2019-09-22 [1] CRAN (R 4.2.0)
##  plyr               1.8.7     2022-03-24 [1] CRAN (R 4.2.0)
##  purrr            * 0.3.5     2022-10-06 [1] CRAN (R 4.2.1)
##  R6                 2.5.1     2021-08-19 [1] CRAN (R 4.2.0)
##  Rcpp               1.0.9     2022-07-08 [1] CRAN (R 4.2.1)
##  RCurl              1.98-1.9  2022-10-03 [1] CRAN (R 4.2.1)
##  readr            * 2.1.3     2022-10-01 [1] CRAN (R 4.2.1)
##  readxl             1.4.1     2022-08-17 [1] CRAN (R 4.2.1)
##  reprex             2.0.2     2022-08-17 [1] CRAN (R 4.2.1)
##  reshape2           1.4.4     2020-04-09 [1] CRAN (R 4.2.0)
##  rlang              1.0.6     2022-09-24 [1] CRAN (R 4.2.1)
##  rmarkdown          2.17      2022-10-07 [1] CRAN (R 4.2.1)
##  rprojroot          2.0.3     2022-04-02 [1] CRAN (R 4.2.0)
##  rstatix            0.7.0     2021-02-13 [1] CRAN (R 4.2.1)
##  rstudioapi         0.14      2022-08-22 [1] CRAN (R 4.2.1)
##  rvest              1.0.3     2022-08-19 [1] CRAN (R 4.2.1)
##  S4Vectors          0.34.0    2022-04-26 [1] Bioconductor
##  sandwich           3.0-2     2022-06-15 [1] CRAN (R 4.2.1)
##  scales             1.2.1     2022-08-20 [1] CRAN (R 4.2.1)
##  sessioninfo        1.2.2     2021-12-06 [1] CRAN (R 4.2.0)
##  stringi            1.7.8     2022-07-11 [1] CRAN (R 4.2.1)
##  stringr          * 1.4.1     2022-08-20 [1] CRAN (R 4.2.1)
##  survival           3.4-0     2022-08-09 [1] CRAN (R 4.2.1)
##  TH.data            1.1-1     2022-04-26 [1] CRAN (R 4.2.1)
##  tibble           * 3.1.8     2022-07-22 [1] CRAN (R 4.2.1)
##  tidyr            * 1.2.1     2022-09-08 [1] CRAN (R 4.2.1)
##  tidyselect         1.2.0     2022-10-10 [1] CRAN (R 4.2.1)
##  tidytree           0.4.1     2022-09-26 [1] CRAN (R 4.2.1)
##  tidyverse        * 1.3.2     2022-07-18 [1] CRAN (R 4.2.1)
```

```
##  treeio             1.23.0    2022-11-10 [1] Github (GuangchuangYu/treeio@db85803)
##  tzdb               0.3.0     2022-03-28 [1] CRAN (R 4.2.0)
##  utf8               1.2.2     2021-07-24 [1] CRAN (R 4.2.0)
##  vctrs              0.5.0     2022-10-22 [1] CRAN (R 4.2.1)
##  vipor              0.4.5     2017-03-22 [1] CRAN (R 4.2.1)
##  vroom              1.6.0     2022-09-30 [1] CRAN (R 4.2.1)
##  withr              2.5.0     2022-03-03 [1] CRAN (R 4.2.0)
##  xfun               0.33      2022-09-12 [1] CRAN (R 4.2.1)
##  xml2               1.3.3     2021-11-30 [1] CRAN (R 4.2.0)
##  XVector            0.36.0    2022-04-26 [1] Bioconductor
##  yaml               2.3.5     2022-02-21 [1] CRAN (R 4.2.0)
##  yulab.utils        0.0.5     2022-06-30 [1] CRAN (R 4.2.1)
##  zlibbioc           1.42.0    2022-04-26 [1] Bioconductor
##  zoo                1.8-11    2022-09-17 [1] CRAN (R 4.2.1)
##
##  [1] /home/faalm/R/x86_64-pc-linux-gnu-library/4.2
##  [2] /usr/local/lib/R/site-library
##  [3] /usr/lib/R/site-library
##  [4] /usr/lib/R/library
##
## ------------------------------------------------------------------------------
```

### References

Consortium, The UniProt. 2021. “UniProt: The Universal Protein Knowledgebase in 2021.” *Nucleic Acids Research* 49 (D1): D480–89.

Hernandez-Plaza, Ana, Damian Szklarczyk, Jorge Botas, Carlos P Cantalapiedra, Joaquin Giner-Lamia, Daniel R Mende, Rebecca Kirsch, et al. 2023. “eggNOG 6.0: Enabling Comparative Genomics Across 12 535 Organisms.” *Nucleic Acids Research* 51 (D1): D389–94.

Kuznetsov, Dmitry, Fredrik Tegenfeldt, Mose Manni, Mathieu Seppey, Matthew Berkeley, Evgenia V Kriventseva, and Evgeny M Zdobnov. 2023. “OrthoDB V11: Annotation of Orthologs in the Widest Sampling of Organismal Diversity.” *Nucleic Acids Research* 51 (D1): D445–51.

Penel, Simon, Anne-Muriel Arigon, Jean-François Dufayard, Anne-Sophie Sertier, Vincent Daubin, Laurent Duret, Manolo Gouy, and Guy Perrière. 2009. “Databases of Homologous Gene Families for Comparative Genomics.” In *BMC Bioinformatics*, 10:1–13. 6. BioMed Central.

Van Bel, Michiel, Francesca Silvestri, Eric M Weitz, Lukasz Kreft, Alexander Botzki, Frederik Coppens, and Klaas Vandepoele. 2022. “PLAZA 5.0: Extending the Scope and Power of Comparative and Functional Genomics in Plants.” *Nucleic Acids Research* 50 (D1): D1468–74.
