## Supplementary Text S3 for "Assessing the quality of comparative genomics data and results with the *cogeqc* R/Bioconductor package"

### Supplementary Text S3: Assessing orthogroup inference for Brassicaceae genomes

***Fabricio Almeida-Silva*** [1,2] ***and Yves Van de Peer*** [1,2,3,4]

[1]VIB-UGent Center for Plant Systems Biology, Ghent, Belgium
[2]Department of Plant Biotechnology and Bioinformatics, Ghent University, Ghent, Belgium
[3]College of Horticulture, Academy for Advanced Interdisciplinary Studies, Nanjing Agricultural University, Nanjing, China
[4]Center for Microbial Ecology and Genomics, Department of Biochemistry, Genetics and Microbiology, University of Pretoria, Pretoria, South Africa

**6 February 2023**

```r
library(here)
library(cogeqc)
library(tidyverse)
library(ggpubr)
library(rstatix)
library(clusterProfiler)
library(enrichplot)
library(patchwork)
library(dplyr)

source(here("code", "utils.R"))
```

### 1 Overview

Here, we will compare the protein domain-based approach in `cogeqc` to assess the impact of multiple combinations of parameters in OrthoFinder (Emms and Kelly 2019) in the accuracy of orthogroup inference. The data set used here will be a collection of Brassicaceae genomes. The parameters we will change are:

1. Program (`-S` option)

- DIAMOND
- DIAMOND ultrasensitive

2. MCL inflation parameter (`-I`)

- 1
- 1.5 (default)
- 2
- 3

### 2 Orthogroup inference

To start, we will load the proteome data and export each proteome as a FASTA file in the `data` directory, so we can pass it to OrthoFinder.

```r
# Load proteomes
load(here("data", "brassicaceae_proteomes.rda"))

# Write files to data/
lapply(seq_along(brassicaceae_proteomes), function(x) {
    outfile <- here("data", paste0(names(brassicaceae_proteomes)[x], ".fasta"))
    Biostrings::writeXStringSet(
        brassicaceae_proteomes[[x]], outfile
    )
})
```

Now, we can run OrthoFinder for each combination of parameters. Here, we created 2 different bash scripts for each DIAMOND mode. They are:

- `of_diamond.sh`: code to run DIAMOND (default mode) for different inflation parameters;

- `of_diamond_ultra.sh`: code to run DIAMOND in ultrasensitive mode for different inflation parameters

The 2 files can be run with:

```
bash of_diamond.sh
bash of_diamond_ultra.sh
```

The *Orthogroups.tsv* files were all moved to the directory `products/result_files`.

#### 2.1 Exploratory analysis of orthogroup inference results

Now that we have the *Orthogroups.tsv* files from OrthoFinder, let's load them.

```
# Extract tar.xz file
tarfile <- here("products", "result_files", "Orthogroups.tar.xz")
outdir <- tempdir()

system2("tar", args = c("-xf", tarfile, "--directory", outdir))

# Get path to OrthoFinder output
og_files <- list.files(
    path = outdir,
    pattern = "Orthogroups.*", full.names = TRUE
)

# Read and parse files
ogs <- lapply(og_files, function(x) {
    og <- read_orthogroups(x)
    og <- og %>%
        mutate(Species = stringr::str_replace_all(Species, "\\.", "")) %>%
        mutate(Gene = str_replace_all(
            Gene, c(
                "\\.[0-9]$" = "",
                "\\.[0-9]\\.p$" = "",
                "\\.t[0-9]$" = "",
                "\\.g$" = ""
            )
        ))
    return(og)
})
og_names <- gsub("\\.tsv", "", basename(og_files))
og_names <- gsub("Orthogroups_", "", og_names)

names(ogs) <- og_names
```

Let's explore OG sizes for each combination of parameters and filter orthogroups by size to remove orthogroups that are artificially large.

```
# Visualize OG sizes
og_sizes_plot <- patchwork::wrap_plots(
    plot_og_sizes(ogs$default_1) + ggtitle("Default, mcl = 1"),
    plot_og_sizes(ogs$default_1_5) + ggtitle("Default, mcl = 1.5") +
```

```
        theme(axis.text.y = element_blank()),
    plot_og_sizes(ogs$default_2) + ggtitle("Default, mcl = 2") +
        theme(axis.text.y = element_blank()),
    plot_og_sizes(ogs$default_3) + ggtitle("Default, mcl = 3") +
        theme(axis.text.y = element_blank()),
    plot_og_sizes(ogs$ultra_1) + ggtitle("Ultra, mcl = 1") +
        theme(axis.text.y = element_blank()),
    plot_og_sizes(ogs$ultra_1_5) + ggtitle("Ultra, mcl = 1.5") +
        theme(axis.text.y = element_blank()),
    plot_og_sizes(ogs$ultra_2) + ggtitle("Ultra, mcl = 2") +
        theme(axis.text.y = element_blank()),
    plot_og_sizes(ogs$ultra_3) + ggtitle("Ultra, mcl = 3") +
        theme(axis.text.y = element_blank()),
    nrow = 1, ncol = 8
)

og_sizes_plot
```

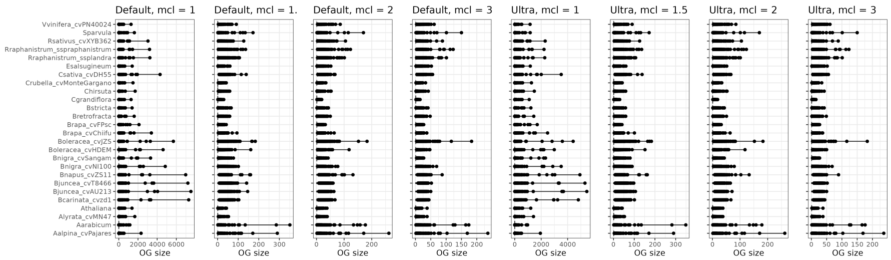

**Figure 1:** Orthogroup sizes for each run.

Expectedly, OrthoFinder runs with mcl inflation parameters of 1 lead to very large orthogroups, including some orthogroups with thousands of genes.

Now, let's explore the percentage of orthogroups with >200, >100, and >50 genes in each OrthoFinder run.

```
# Calculate OG sizes for each run
og_sizes <- lapply(ogs, function(x) {
    sizes <- as.matrix(table(x$Orthogroup, x$Species))
    total <- rowSums(sizes)

    sizes_df <- data.frame(unclass(sizes))
    sizes_df$Total <- total
    return(sizes_df)
})

# What is the percentage of OGs with >=100 genes? And with >50 genes?
percentage_size <- function(size_df, n = 100) {
    return(sum(size_df$Total > n) / nrow(size_df) * 100)
}

percentages <- data.frame(
```

```
    Mode = names(og_sizes),
    P200 = unlist(lapply(og_sizes, percentage_size, n = 200)),
    P100 = unlist(lapply(og_sizes, percentage_size, n = 100)),
    P50 = unlist(lapply(og_sizes, percentage_size, n = 50)),
    OGs = unlist(lapply(og_sizes, nrow))
)

# Reorder rows from lowest to highest mcl inflation
orders <- c(
    "default_1", "default_1_5", "default_2", "default_3",
    "ultra_1", "ultra_1_5", "ultra_2", "ultra_3"
)
percentages <- percentages[orders, ]

# Visual exploration
percentage_plot <- percentages %>%
    tidyr::pivot_longer(cols = !Mode) %>%
    mutate(name = str_replace_all(
        name,
        c(
            "OGs" = "Number of OGs",
            "P200" = "% OGs with >200 genes",
            "P100" = "% OGs with >100 genes",
            "P50" = "% OGs with >50 genes"
        )
    )) %>%
    ggplot(., aes(y = Mode, x = value)) +
    geom_col(aes(fill = Mode), show.legend = "none") +
    scale_fill_manual(
        values = c("ultra_3" = "#08519C", "ultra_2" = "#3182BD",
                   "ultra_1_5" = "#6BAED6", "ultra_1" = "#BDD7E7",
                   "default_3" = "#006D2C", "default_2" = "#31A354",
                   "default_1_5" = "#74C476", "default_1" = "#BAE4B3")
    ) +
    facet_wrap(~name, ncol = 4, scales = "free_x") +
    theme_bw() +
    labs(
        x = "", y = "OrthoFinder mode",
        title = "Relationship between the number of orthogroups and orthogroup size per OrthoFinder mode"
    )

percentage_plot
```

It is very clear that increasing the mcl inflation increases the number of orthogroups, but decreases the percentage of OGs with more than 100 and 50 genes.

Finally, let's remove OGs with >=200 genes to remove noise.

```
# Filter OGs
ogs_filtered <- lapply(seq_along(ogs), function(x) {

    # Which OGs less than 200 genes?
```

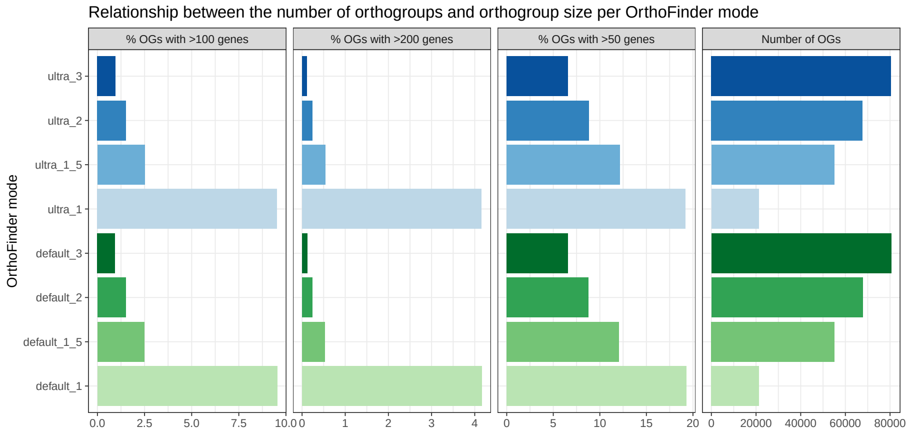

**Figure 2:** Percentage of orthogroups with >50, >100, and >200 genes for each run.

```
    og_keep <- rownames(og_sizes[[x]][og_sizes[[x]]$Total < 200, ])

    fogs <- ogs[[x]][ogs[[x]]$Orthogroup %in% og_keep, ]
    return(fogs)
})
names(ogs_filtered) <- names(ogs)
```

### 3 Orthogroup assessment

Now, let's get InterPro domain annotation for the following species to assess orthogroups:

- *A. thaliana*
- *A. arabicum*
- *A. lyrata*
- *B. carinata*
- *C. rubella*
- *C. hirsuta*
- *S. parvula*

```
# Define function to read functional annotation from PLAZA 5.0
read_annotation <- function(url, cols = c(1, 3)) {
    annot <- readr::read_tsv(url, show_col_types = FALSE, skip = 8) %>%
        select(cols)
    names(annot)[1:2] <- c("Gene", "Annotation")
    return(annot)
}
```

```
# Get Interpro annotation
base <- "https://ftp.psb.ugent.be/pub/plaza/plaza_public_dicots_05/InterPro/"
interpro <- list(
    Athaliana = read_annotation(paste0(base, "interpro.ath.csv.gz")),
    Aarabicum = read_annotation(paste0(base, "interpro.aar.csv.gz")),
```

```
    Alyrata_cvMN47 = read_annotation(paste0(base, "interpro.aly.csv.gz")),
    Bcarinata_cvzd1 = read_annotation(paste0(base, "interpro.bca.csv.gz")),
    Crubella_cvMonteGargano = read_annotation(paste0(base, "interpro.cru.csv.gz")),
    Chirsuta = read_annotation(paste0(base, "interpro.chi.csv.gz")),
    Sparvula = read_annotation(paste0(base, "interpro.spa.csv.gz"))
)
interpro <- lapply(interpro, as.data.frame)

# Calculate homogeneity scores
species_annotation <- names(interpro)
og_assessment <- lapply(seq_along(ogs_filtered), function(x) {

    message("Working on mode ", names(ogs_filtered)[x])
    orthogroups <- ogs_filtered[[x]]
    orthogroups <- orthogroups[orthogroups$Species %in% species_annotation, ]

    res <- assess_orthogroups(orthogroups, interpro)
    res$Mode <- factor(
        names(ogs_filtered)[x],
        levels = c(
            "ultra_3", "ultra_2", "ultra_1_5", "ultra_1",
            "default_3", "default_2", "default_1_5", "default_1"
        )
    )
    return(res)
})
og_assessment <- Reduce(rbind, og_assessment)

# Save homogeneity stats
save(
    og_assessment, compress = "xz",
    file = here("products", "result_files", "og_assessment_brassicaceae.rda")
)
```

### 4 Comparing and visualizing homogeneity statistics

Here, we will compare and visualize how the homogeneity scores are affected by:

- different species choice
- different mcl inflation values
- different DIAMOND modes (default and ultra)

Quick exploration of median and mean homogeneity:

```
load(here("products", "result_files", "og_assessment_brassicaceae.rda"))

# Scale value to the maximum so that values range from 0 to 1
og_assessment$Median_score <- og_assessment$Median_score /
    max(og_assessment$Median_score)

# Mean
```

```
mean_og <- og_assessment %>%
    group_by(Mode) %>%
    summarise(mean = mean(Median_score))

# Median
median_og <- og_assessment %>%
    group_by(Mode) %>%
    summarise(median = median(Median_score))

mean_and_median_og <- inner_join(mean_og, median_og) |>
    dplyr::rename(Mean = mean, Median = median)

knitr::kable(mean_and_median_og, caption = "Mean and median OG scores.", digits = 3)
```

**Table 1:** Mean and median OG scores.

| Mode | Mean | Median |
|---|---|---|
| ultra_3 | 0.640 | 0.640 |
| ultra_2 | 0.631 | 0.635 |
| ultra_1_5 | 0.620 | 0.628 |
| ultra_1 | 0.425 | 0.424 |
| default_3 | 0.639 | 0.640 |
| default_2 | 0.631 | 0.635 |
| default_1_5 | 0.620 | 0.628 |
| default_1 | 0.425 | 0.423 |

#### 4.1 Global distributions

Here, we will compare and visualize all distros considering different DIAMOND modes and mcl inflation values. To start, let's perform Wilcoxon tests for all combinations of modes and obtain effect sizes.

```
# Relevel 'Mode' factor
og_assessment$Mode <- factor(
    og_assessment$Mode,
    levels = c(
        "ultra_3", "ultra_2", "ultra_1_5", "ultra_1",
        "default_3", "default_2", "default_1_5", "default_1"
    )
)

# Comparing all vs all
comp_global <- compare(og_assessment, "Median_score ~ Mode")
comp_global |>
    filter_comparison() |>
    knitr::kable(
        caption = "Mann-Whitney U test for differences in orthogroup scores with Wilcoxon effect sizes.",
        digits = 10
    )
```

**Table 2:** Mann-Whitney U test for differences in orthogroup scores with Wilcoxon effect sizes.

| group1 | group2 | n1 | n2 | padj | effsize | magnitude |
|---|---|---|---|---|---|---|
| ultra_3 | ultra_2 | 19738 | 18575 | 0.00e+00 | 0.04120347 | small |
| ultra_3 | ultra_1_5 | 19738 | 16898 | 0.00e+00 | 0.06125964 | small |
| ultra_3 | ultra_1 | 19738 | 5534 | 0.00e+00 | 0.34087185 | moderate |
| ultra_3 | default_3 | 19738 | 19765 | 5.00e-10 | 0.03113134 | small |
| ultra_3 | default_2 | 19738 | 18633 | 0.00e+00 | 0.04197169 | small |
| ultra_3 | default_1_5 | 19738 | 16975 | 0.00e+00 | 0.06233198 | small |
| ultra_3 | default_1 | 19738 | 5587 | 0.00e+00 | 0.34258513 | moderate |
| ultra_2 | ultra_1_5 | 18575 | 16898 | 0.00e+00 | 0.04340346 | small |
| ultra_2 | ultra_1 | 18575 | 5534 | 0.00e+00 | 0.33536590 | moderate |
| ultra_2 | default_3 | 18575 | 19765 | 0.00e+00 | 0.04018176 | small |
| ultra_2 | default_2 | 18575 | 18633 | 2.55e-08 | 0.02855053 | small |
| ultra_2 | default_1_5 | 18575 | 16975 | 0.00e+00 | 0.04451653 | small |
| ultra_2 | default_1 | 18575 | 5587 | 0.00e+00 | 0.33742024 | moderate |
| ultra_1_5 | ultra_1 | 16898 | 5534 | 0.00e+00 | 0.32401575 | moderate |
| ultra_1_5 | default_3 | 16898 | 19765 | 0.00e+00 | 0.05737302 | small |
| ultra_1_5 | default_2 | 16898 | 18633 | 0.00e+00 | 0.04259062 | small |
| ultra_1_5 | default_1_5 | 16898 | 16975 | 1.72e-06 | 0.02554732 | small |
| ultra_1_5 | default_1 | 16898 | 5587 | 0.00e+00 | 0.32541386 | moderate |
| ultra_1 | default_3 | 5534 | 19765 | 0.00e+00 | 0.34014551 | moderate |
| ultra_1 | default_2 | 5534 | 18633 | 0.00e+00 | 0.33456485 | moderate |
| ultra_1 | default_1_5 | 5534 | 16975 | 0.00e+00 | 0.32260079 | moderate |
| ultra_1 | default_1 | 5534 | 5587 | 2.70e-02 | 0.01928759 | small |
| default_3 | default_2 | 19765 | 18633 | 0.00e+00 | 0.04084430 | small |
| default_3 | default_1_5 | 19765 | 16975 | 0.00e+00 | 0.06124558 | small |
| default_3 | default_1 | 19765 | 5587 | 0.00e+00 | 0.34147823 | moderate |
| default_2 | default_1_5 | 18633 | 16975 | 0.00e+00 | 0.04366051 | small |
| default_2 | default_1 | 18633 | 5587 | 0.00e+00 | 0.33616829 | moderate |
| default_1_5 | default_1 | 16975 | 5587 | 0.00e+00 | 0.32420710 | moderate |

As we can see, using mcl = 1 leads to much smaller homogeneity scores as compared to every other mcl value. For mcl values >=1.5, there are differences, but they are likely due to large sample sizes, as indicated by small effect sizes.

The default OrthoFinder mode (default DIAMOND, mcl = 1.5) leads to higher homogeneity as compared to runs using mcl = 1, both in default and ultrasensitive DIAMOND modes. The difference between the default mode and runs with higher mcl values are negligible.

Now, let's visualize the distributions and compare the default OrthoFinder mode with every other mode, highlighting significant differences ($P < 0.05$) with effect size > 0.1.

```
# Visualize
global_comps <- list(
    c("default_1_5", "ultra_1"),
    c("default_1_5", "default_1")
)

p_distros_global <- ggviolin(
```

```
    og_assessment, y = "Median_score", x = "Mode",
    orientation = "horiz", trim = TRUE,
    add = c("boxplot", "mean"),
    fill = "Mode", add.params = list(fill = "white")
) +
    scale_fill_manual(
        values = c("ultra_3" = "#08519C", "ultra_2" = "#3182BD",
                   "ultra_1_5" = "#6BAED6", "ultra_1" = "#BDD7E7",
                   "default_3" = "#006D2C", "default_2" = "#31A354",
                   "default_1_5" = "#74C476", "default_1" = "#BAE4B3")
    ) +
    stat_compare_means(
        comparisons = global_comps, label = "p.signif",
        method = "wilcox.test"
    ) +
    theme(legend.position = "none") +
    labs(y = "Scaled homogeneity scores", x = "OrthoFinder modes",
         title = "Distribution of mean homogeneity scores for orthogroups") +
    theme(plot.subtitle = ggtext::element_markdown())

p_distros_global
```

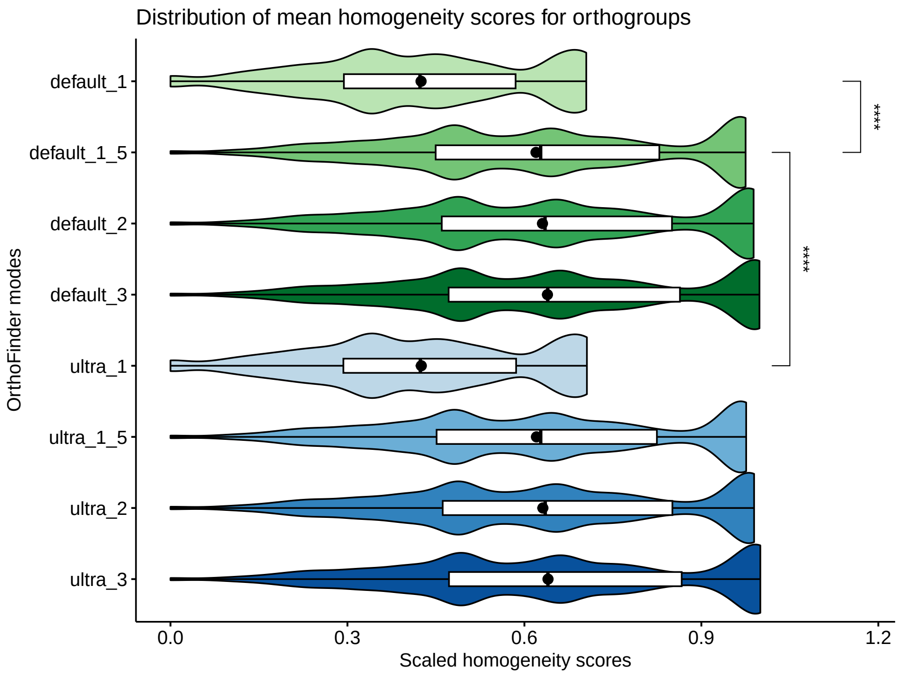

**Figure 3:** Distribution of mean orthogroup scores for each OrthoFinder run.

#### 4.2 The effect of species choice

Here, we will compare the distributions of orthogroups scores using each species individually to see if the species choice has an impact on the conclusions.

```
og_species_long <- Reduce(rbind, lapply(2:8, function(x) {

    var <- names(og_assessment)[x]
    species_name <- gsub("_.*", "", var)

    long_df <- og_assessment[, c("Orthogroups", var, "Mode")]
    names(long_df) <- c("OGs", "Score", "Mode")
    long_df$Score <- long_df$Score / max(long_df$Score, na.rm = TRUE)
    long_df$Species <- species_name

    return(long_df)
}))

og_species_long <- og_species_long[!is.na(og_species_long$Score), ]
og_species_long <- og_species_long |>
    mutate(
        Species = str_replace_all(
            Species,
            c(
                "Aarabicum" = "A. arabicum",
                "Alyrata" = "A. lyrata",
                "Athaliana" = "A. thaliana",
                "Bcarinata" = "B. carinata",
                "Chirsuta" = "C. hirsuta",
                "Crubella" = "C. rubella",
                "Sparvula" = "S. parvula"
            )
        )
    )

p_distros_by_species <- ggviolin(
    og_species_long,
    y = "Score", x = "Mode",
    orientation = "horiz", trim = TRUE,
    add = c("boxplot", "mean"), facet.by = "Species", nrow = 1,
    fill = "Mode", add.params = list(fill = "white")
) +
    scale_fill_manual(
        values = c(
            "ultra_3" = "#08519C", "ultra_2" = "#3182BD",
            "ultra_1_5" = "#6BAED6", "ultra_1" = "#BDD7E7",
            "default_3" = "#006D2C", "default_2" = "#31A354",
            "default_1_5" = "#74C476", "default_1" = "#BAE4B3"
        )
    ) +
    theme(legend.position = "none") +
    labs(
        y = "Scaled homogeneity scores", x = "OrthoFinder modes",
        title = "Distribution of OG scores for each species"
    ) +
```

```
        scale_x_discrete(
            labels = c(
                "default_1" = "Default, 1",
                "default_1_5" = "Default, 1.5",
                "default_2" = "Default, 2",
                "default_3" = "Default, 3",
                "ultra_1" = "Ultra, 1",
                "ultra_1_5" = "Ultra, 1.5",
                "ultra_2" = "Ultra, 2",
                "ultra_3" = "Ultra, 3"
            )
        ) +
        theme(axis.text.x = element_text(angle = 60, vjust = 0.5))

p_distros_by_species
```

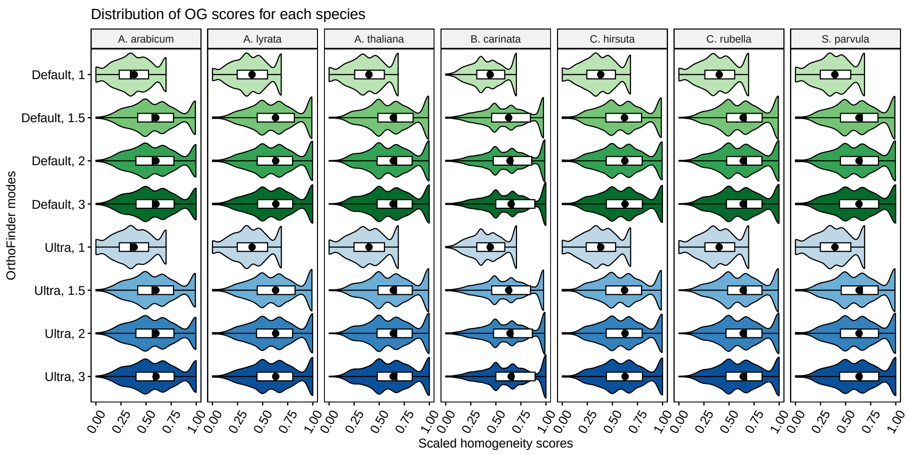

**Figure 4:** Distribution of orthogroup scores for each OrthoFinder run calculated for each species separately.

We conclude that the species choice does not affect the comparisons of orthogroup scores among OrthoFinder runs.

#### 4.3 The effect of mcl inflation parameters

Here, we will explore the impact of changing mcl inflation parameters in the homogeneity of orthogroups.

```
# Process data to include information on DIAMOND mode and mcl
og_modes <- og_assessment %>%
    mutate(diamond = str_replace_all(Mode, "_.*", "")) %>%
    mutate(mcl = str_replace_all(Mode, c("default_" = "", "ultra_" = ""))) %>%
    mutate(mcl = str_replace_all(mcl, "_", ".")) %>%
    mutate(mcl = as.numeric(mcl))

# Obtain P-values from Wilcoxon tests and effect sizes
comp_mcl_default <- og_modes %>%
    filter(diamond == "default") %>%
```

```
    compare(., "Median_score ~ mcl")

comp_mcl_default |>
    filter_comparison() |>
    knitr::kable(
        caption = "Mann-Whitney U test for differences in orthogroup scores between runs with different mcl p
        digits = 10
    )
```

**Table 3:** Mann-Whitney U test for differences in orthogroup scores between runs with different mcl parameters and standard DIAMOND mode. Effect sizes represent Wilcoxon effect sizes.

| group1 | group2 | n1 | n2 | padj | effsize | magnitude |
|---|---|---|---|---|---|---|
| 1 | 1.5 | 5587 | 16975 | 0 | 0.32420710 | moderate |
| 1 | 2 | 5587 | 18633 | 0 | 0.33616829 | moderate |
| 1 | 3 | 5587 | 19765 | 0 | 0.34147823 | moderate |
| 1.5 | 2 | 16975 | 18633 | 0 | 0.04366051 | small |
| 1.5 | 3 | 16975 | 19765 | 0 | 0.06124558 | small |
| 2 | 3 | 18633 | 19765 | 0 | 0.04084430 | small |

```
comp_mcl_ultra <- og_modes %>%
    filter(diamond == "ultra") %>%
    compare(., "Median_score ~ mcl")

comp_mcl_ultra |>
    filter_comparison() |>
    knitr::kable(
        caption = "Mann-Whitney U test for differences in orthogroup scores between runs with different mcl p
        digits = 10
    )
```

**Table 4:** Mann-Whitney U test for differences in orthogroup scores between runs with different mcl parameters and ultra-sensitive DIAMOND mode. Effect sizes represent Wilcoxon effect sizes.

| group1 | group2 | n1 | n2 | padj | effsize | magnitude |
|---|---|---|---|---|---|---|
| 1 | 1.5 | 5534 | 16898 | 0 | 0.32401575 | moderate |
| 1 | 2 | 5534 | 18575 | 0 | 0.33536590 | moderate |
| 1 | 3 | 5534 | 19738 | 0 | 0.34087185 | moderate |
| 1.5 | 2 | 16898 | 18575 | 0 | 0.04340346 | small |
| 1.5 | 3 | 16898 | 19738 | 0 | 0.06125964 | small |
| 2 | 3 | 18575 | 19738 | 0 | 0.04120347 | small |

In line with what we demonstrated in the global distributions, the Wilcoxon tests show that mcl $= 1$ leads to much lower homogeneity scores than all other mcl values, regardless of the DIAMOND mode. Additionally, increasing mcl values leads to increased homogeneity scores (i.e., homogeneity scores follow the order of mcl $3 > 2 > 1.5 > 1$), but differences among mcl values $>=$1.5 are negligible, as indicated by small effect sizes. Thus, low P-values could be due to large sample sizes.

Now, let's visualize the distributions.

```
# List of comparisons to be made
mcl_comp <- list(
    c("1", "1.5"), c("1", "2"), c("1", "3"), c("1.5", "3")
)

# Plot
p_distros_mcl <- og_assessment %>%
    mutate(diamond = str_replace_all(Mode, "_.*", "")) %>%
    mutate(mcl = str_replace_all(Mode, c("default_" = "", "ultra_" = ""))) %>%
    mutate(mcl = str_replace_all(mcl, "_", ".")) %>%
    mutate(mcl = as.numeric(mcl)) %>%
    ggviolin(., x = "mcl", y = "Median_score", trim = TRUE,
             add = c("boxplot", "mean"), facet.by = "diamond",
             fill = "Mode", add.params = list(fill = "white")) +
    theme(legend.position = "none") +
    scale_fill_manual(
        values = c("ultra_3" = "#08519C", "ultra_2" = "#3182BD",
                   "ultra_1_5" = "#6BAED6", "ultra_1" = "#BDD7E7",
                   "default_3" = "#006D2C", "default_2" = "#31A354",
                   "default_1_5" = "#74C476", "default_1" = "#BAE4B3")
    ) +
    stat_compare_means(
        comparisons = mcl_comp, label = "p.signif",
        method = "wilcox.test"
    ) +
    labs(
        y = "Scaled homogeneity scores", x = "MCL inflation parameters",
        title = "Effect of MCL inflation values on orthogroup inference",
        subtitle = "Panels represent DIAMOND sensitivity modes"
    )

p_distros_mcl
```

#### 4.4 The effect of DIAMOND mode (default vs ultra)

Here, we will investigate whether changing the DIAMOND mode (default vs ultrasensitive) in OrthoFinder affects orthogroup homogeneity.

```
# Compare median scores
mcl1 <- og_modes %>%
    filter(mcl == 1) %>%
    compare(., "Median_score ~ diamond") |>
    filter_comparison()

mcl1_5 <- og_modes %>%
    filter(mcl == 1.5) %>%
    compare(., "Median_score ~ diamond") |>
    filter_comparison()

mcl2 <- og_modes %>%
```

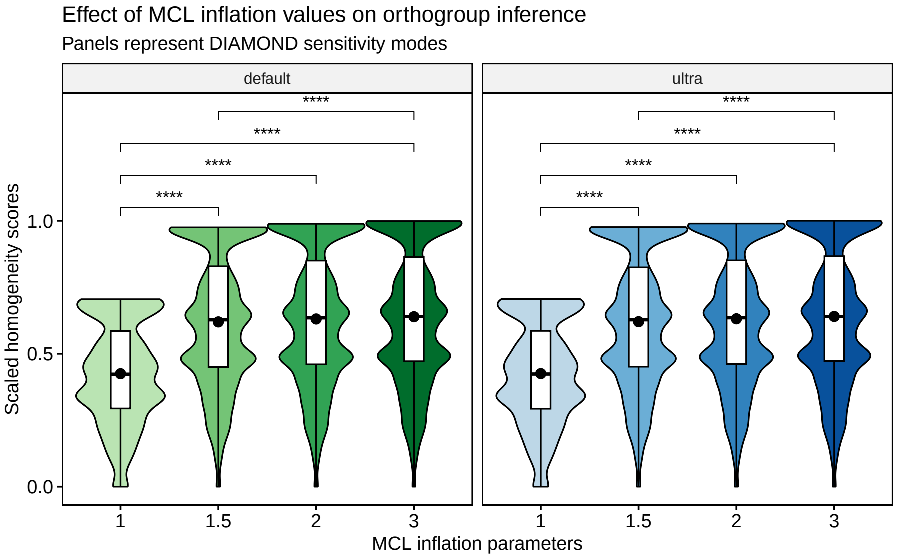

**Figure 5:** Effect of MCL inflation values on orthogroup scores.

```
    filter(mcl == 2) %>%
    compare(., "Median_score ~ diamond") |>
    filter_comparison()

mcl3 <- og_modes %>%
    filter(mcl == 3) %>%
    compare(., "Median_score ~ diamond") |>
    filter_comparison()

bind_rows(
    mcl1 |> mutate(mcl = 1),
    mcl1_5 |> mutate(mcl = 1.5),
    mcl2 |> mutate(mcl = 2),
    mcl3 |> mutate(mcl = 3)
) |>
    knitr::kable(
        caption = "Mann-Whitney U test for differences in orthogroup scores between runs with different DIAMO
        digits = 10
    )
```

**Table 5:** Mann-Whitney U test for differences in orthogroup scores between runs with different DIAMOND modes for each mcl value. Effect sizes represent Wilcoxon effect sizes.

| group1 | group2 | n1 | n2 | padj | effsize | magnitude | mcl |
|---|---|---|---|---|---|---|---|
| default | ultra | 5587 | 5534 | 2.10e-02 | 0.01928759 | small | 1.0 |
| default | ultra | 16975 | 16898 | 1.29e-06 | 0.02554732 | small | 1.5 |
| default | ultra | 18633 | 18575 | 1.82e-08 | 0.02855053 | small | 2.0 |
| default | ultra | 19765 | 19738 | 3.00e-10 | 0.03113134 | small | 3.0 |

Again, we can see that there are significant P-values, but very small effect sizes, indicating no difference resulting from the DIAMOND mode. Thus, users can run the default mode of DIAMOND, which is way faster, without any loss of biological signal for orthogroup inference.

Let's visualize the distributions.

```
# Plot
p_distros_diamond <- og_modes %>%
    ggviolin(., x = "diamond", y = "Median_score", trim = TRUE,
             add = c("boxplot", "mean"), facet.by = "mcl", ncol = 4,
             fill = "Mode", add.params = list(fill = "white")) +
    theme(legend.position = "none") +
    scale_fill_manual(
        values = c("ultra_3" = "#08519C", "ultra_2" = "#3182BD",
                   "ultra_1_5" = "#6BAED6", "ultra_1" = "#BDD7E7",
                   "default_3" = "#006D2C", "default_2" = "#31A354",
                   "default_1_5" = "#74C476", "default_1" = "#BAE4B3")
    ) +
    labs(y = "Scaled homogeneity scores", x = "DIAMOND mode",
         title = "Effect of DIAMOND sensitivity mode on orthogroup inference",
         subtitle = "Panels represent MCL inflation parameters") +
    theme(plot.subtitle = ggtext::element_markdown())

p_distros_diamond
```

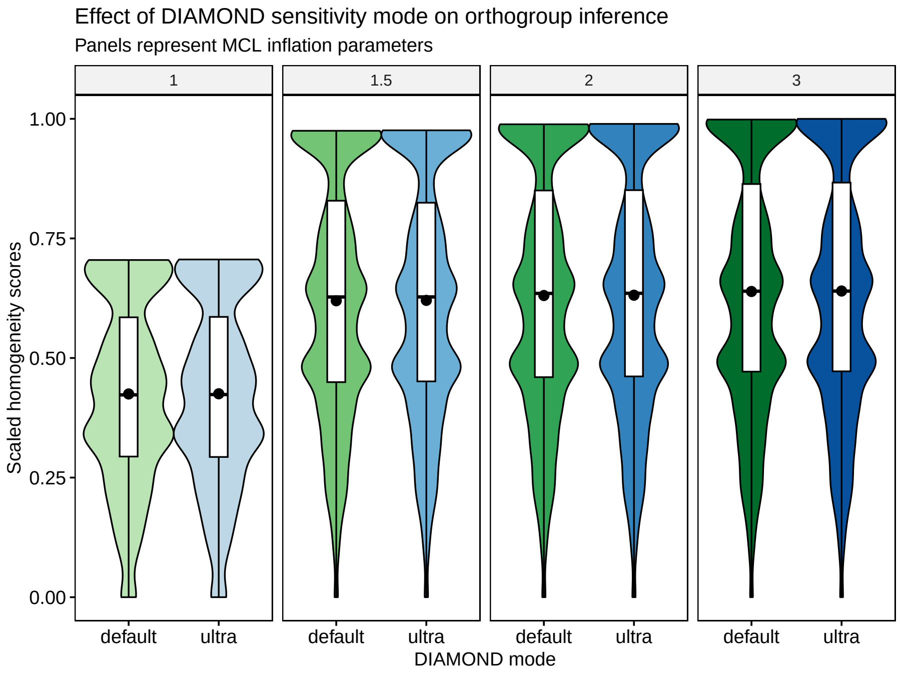

**Figure 6:** Effect of DIAMOND mode on orthogroup scores.

### 5 Functional analysis of homogeneous and heterogeneous gene families

By looking at the global distributions of homogeneity scores, we can see that all distributions have a similar shape. This pattern suggests that some gene families tend to be more homogeneous (scores close to 1), while others tend to include domains that are not shared by all members. The latter can be, for instance, rapidly evolving families that gain or lose domains at faster rates.

To explore what these groups of families contain, we will perform a functional enrichment analysis each group. First of anything, let's plot the distribution for the default OrthoFinder mode and highlight the groups.

```
# Plot distro with groups
p_distros_groups <- og_assessment %>%
    filter(Mode == "default_1_5") %>%
    ggplot(aes(x = Median_score)) +
    geom_density(fill = "grey80", color = "black") +
    ggpubr::theme_pubr() +
    labs(
        y = "Density", x = "Orthogroup scores",
        title = "Distribution of mean homogeneity scores for orthogroups",
        subtitle = "Scores for the default OrthoFinder mode"
    ) +
    geom_vline(xintercept = 0.56, color = "firebrick", linetype = 2) +
    geom_vline(xintercept = 0.87, color = "firebrick", linetype = 2)

p_distros_groups
```

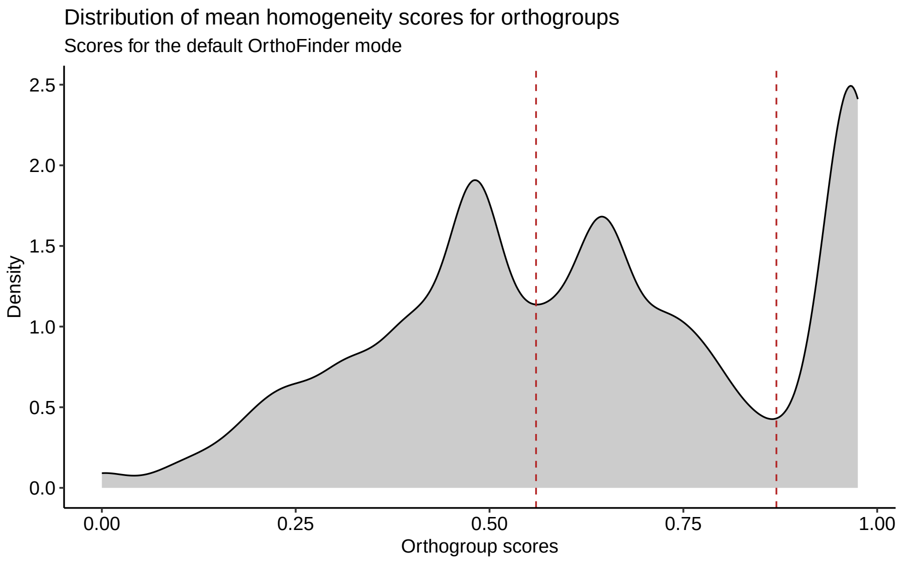

**Figure 7:** Distribution of mean homogeneity scores for orthogroups

Now, let's get vectors of genes in orthogroups from each of the groups highlighted in the figure above.

```
species <- c(
    "Athaliana", "Aarabicum", "Alyrata_cvMN47", "Bcarinata_cvzd1",
    "Crubella_cvMonteGargano", "Chirsuta", "Sparvula"
)

# Get genes and orthogroups (default mode)
genes_ogs <- ogs_filtered$default_1_5

# Keep only species for which we have functional annotation info
genes_ogs <- genes_ogs[genes_ogs$Species %in% species, c(1, 3)]

# Get background genes (all genes in OGs)
background <- genes_ogs$Gene

# Find orthogroups for each group
## G1: 0 - 0.56
g1 <- og_assessment %>%
    filter(Mode == "default_1_5") %>%
    mutate(Median_score = Median_score / max(Median_score)) %>%
    filter(Median_score <= 0.56) %>%
    select(Orthogroups) %>%
    inner_join(., genes_ogs, by = c("Orthogroups" = "Orthogroup")) %>%
    pull(Gene)

## G2: 0.56 - 0.87
g2 <- og_assessment %>%
    filter(Mode == "default_1_5") %>%
    mutate(Median_score = Median_score / max(Median_score)) %>%
    filter(Median_score > 0.56 & Median_score <= 0.87) %>%
    select(Orthogroups) %>%
    inner_join(., genes_ogs, by = c("Orthogroups" = "Orthogroup")) %>%
    pull(Gene)

## G3: 0.87 - 1
g3 <- og_assessment %>%
    filter(Mode == "default_1_5") %>%
    mutate(Median_score = Median_score / max(Median_score)) %>%
    filter(Median_score > 0.87) %>%
    select(Orthogroups) %>%
    inner_join(., genes_ogs, by = c("Orthogroups" = "Orthogroup")) %>%
    pull(Gene)
```

Next, we need to get functional annotation from PLAZA.

```
options(timeout = 6000)
plaza_species <- c("ath", "aar", "aly", "bca", "cru", "chi", "spa")

# GO annotation
bgo <- "https://ftp.psb.ugent.be/pub/plaza/plaza_public_dicots_05/GO/"
go <- lapply(plaza_species, function(x) {
```

```
    y <- read_annotation(paste0(bgo, "go.", x, ".csv.gz"), c(1, 3, 8))
    term2gene <- y[, c(2, 1)] %>% distinct(., .keep_all = TRUE)
    term2name <- y[, c(2, 3)] %>% distinct(., .keep_all = TRUE)
    res <- list(
        TERM2GENE = as.data.frame(term2gene),
        TERM2NAME = as.data.frame(term2name)
    )
    return(res)
})
go_gene <- Reduce(rbind, lapply(go, function(x) return(x$TERM2GENE)))
go_des <- Reduce(rbind, lapply(go, function(x) return(x$TERM2NAME)))

## Remove non-BP terms
ath_bp <- file.path(tempdir(), "ath_bp.rds")
download.file(
    "https://jokergoo.github.io/rGREAT_genesets/genesets/bp_athaliana_eg_gene_go_genesets.rds",
    destfile = ath_bp
)
gobp <- readRDS(ath_bp)
gobp <- names(gobp)
go_gene <- go_gene[go_gene$Annotation %in% gobp, ]
go_des <- go_des[go_des$Annotation %in% gobp, ]
rm(gobp)

# MapMan annotation
bmm <- "https://ftp.psb.ugent.be/pub/plaza/plaza_public_dicots_05/MapMan/"
mm <- lapply(plaza_species, function(x) {
    y <- read_annotation(paste0(bmm, "mapman.", x, ".csv.gz"), c(3:5))
    term2gene <- y[, c(2, 1)] %>% distinct(., .keep_all = TRUE)
    term2name <- y[, c(2, 3)] %>% distinct(., .keep_all = TRUE)
    res <- list(
        TERM2GENE = as.data.frame(term2gene),
        TERM2NAME = as.data.frame(term2name)
    )
    return(res)
})
mm_gene <- Reduce(rbind, lapply(mm, function(x) return(x$TERM2GENE)))
mm_des <- Reduce(rbind, lapply(mm, function(x) return(x$TERM2NAME))) %>%
    mutate(desc = str_replace_all(desc, ".*\\.", ""))

# InterPro
bi <- "https://ftp.psb.ugent.be/pub/plaza/plaza_public_dicots_05/InterPro/"
ip <- lapply(plaza_species, function(x) {
    y <- read_annotation(paste0(bi, "interpro.", x, ".csv.gz"), c(1, 3, 4))
    term2gene <- y[, c(2, 1)] %>% distinct(., .keep_all = TRUE)
    term2name <- y[, c(2, 3)] %>% distinct(., .keep_all = TRUE)
    res <- list(
        TERM2GENE = as.data.frame(term2gene),
        TERM2NAME = as.data.frame(term2name)
    )
    return(res)
```

```
})
ip_gene <- Reduce(rbind, lapply(ip, function(x) return(x$TERM2GENE)))
ip_des <- Reduce(rbind, lapply(ip, function(x) return(x$TERM2NAME)))
```

Now, we can finally perform the enrichment analyses.

```
# Perform enrichment analyses
library(clusterProfiler)

tgene <- list(
    GO = go_gene,
    MapMan = mm_gene,
    InterPro = ip_gene
)
tname <- list(
    GO = go_des,
    MapMan = mm_des,
    InterPro = ip_des
)

## G1
g1_sea <- Reduce(rbind, lapply(seq_along(tgene), function(x) {
    return(as.data.frame(enricher(
        g1, universe = background,
        TERM2GENE = tgene[[x]], TERM2NAME = tname[[x]]
    ))[, 1:6])
}))

## G2
g2_sea <- Reduce(rbind, lapply(seq_along(tgene), function(x) {
    return(as.data.frame(enricher(
        g2, universe = background,
        TERM2GENE = tgene[[x]], TERM2NAME = tname[[x]]
    ))[, 1:6])
}))

## G3
g3_sea <- Reduce(rbind, lapply(seq_along(tgene), function(x) {
    return(as.data.frame(enricher(
        g3, universe = background,
        TERM2GENE = tgene[[x]], TERM2NAME = tname[[x]]
    ))[, 1:6])
}))

# Combine SEA results in a single data frame and export it as a .tsv file
## Combine data frames
sea_res <- rbind(
    g1_sea %>% mutate(group = "G1"),
    g2_sea %>% mutate(group = "G2"),
    g3_sea %>% mutate(group = "G3")
)
```

```r
## Export .tsv
write_tsv(
    sea_res,
    file = here("products", "tables", "enrichment_bygroup.tsv")
)
```

The complete enrichment results are stored in the table `enrichment_bygroup.tsv`. To make visualization and interpretation easier, we will perform semantic similarity analysis to group redundant terms and get a global view of processes associated with each cluster.

Here, we will only use GO terms from the category "Biological Process".

```r
# Semantic similarity analysis for GO-BP terms
## G1
g1_summary <- pairwise_termsim(enricher(
    g1, universe = background,
    TERM2GENE = go_gene, TERM2NAME = go_des
))

## G2
g2_summary <- pairwise_termsim(enricher(
    g2, universe = background,
    TERM2GENE = go_gene, TERM2NAME = go_des
))

## G3
g3_summary <- pairwise_termsim(enricher(
    g3, universe = background,
    TERM2GENE = go_gene, TERM2NAME = go_des
))

# Save objects
save(
    g1_summary, compress = "xz",
    file = here("products", "result_files", "g1_summary.rda")
)

save(
    g2_summary, compress = "xz",
    file = here("products", "result_files", "g2_summary.rda")
)

save(
    g3_summary, compress = "xz",
    file = here("products", "result_files", "g3_summary.rda")
)
```

Now, let's plot the results.

```r
# Tree plot
p_tree_g1 <- treeplot(g1_summary, nWords = 0) +
    ggsci::scale_fill_jama() +
```

```
    ggtitle("Group 1")
p_tree_g1$layers[[4]] <- NULL

p_tree_g2 <- treeplot(g2_summary, nCluster = 7, nWords = 0) +
    ggsci::scale_fill_jama() +
    ggtitle("Group 2")
p_tree_g2$layers[[4]] <- NULL

p_tree_g3 <- treeplot(g3_summary, nWords = 0) +
    ggsci::scale_fill_jama() +
    ggtitle("Group 3")
p_tree_g3$layers[[4]] <- NULL

# Replace P.adj with -log10(P.adj)
p_tree_g1$data$color <- -log10(p_tree_g1$data$color)
p_tree_g2$data$color <- -log10(p_tree_g2$data$color)
p_tree_g3$data$color <- -log10(p_tree_g3$data$color)

# Combine plots in one, with shared legends
rcol <- range(
    c(
        p_tree_g1$data$color, p_tree_g2$data$color, p_tree_g3$data$color
    ),
    na.rm = TRUE
)
rsize <- range(
    c(
        p_tree_g1$data$count, p_tree_g2$data$count, p_tree_g2$data$count
    ),
    na.rm = TRUE
)

wrap_plots(p_tree_g1, p_tree_g2, p_tree_g3) +
    plot_layout(guides = "collect") &
    scale_color_continuous(name = "-Log10(P)", limits = signif(rcol, 2)) &
    scale_size_continuous(name = "Gene count", limits = rsize) &
    theme(legend.position = "bottom")
```

```
# Dot plot
p_dot_g1 <- dotplot(g1_summary, showCategory = 20) + ggtitle("Group 1")
p_dot_g2 <- dotplot(g2_summary, showCategory = 20) + ggtitle("Group 2")
p_dot_g3 <- dotplot(g3_summary, showCategory = 20) + ggtitle("Group 3")

# Replace P.adj with -log10(P.adj)
p_dot_g1$data$p.adjust <- -log10(p_dot_g1$data$p.adjust)
p_dot_g2$data$p.adjust <- -log10(p_dot_g2$data$p.adjust)
p_dot_g3$data$p.adjust <- -log10(p_dot_g3$data$p.adjust)
```

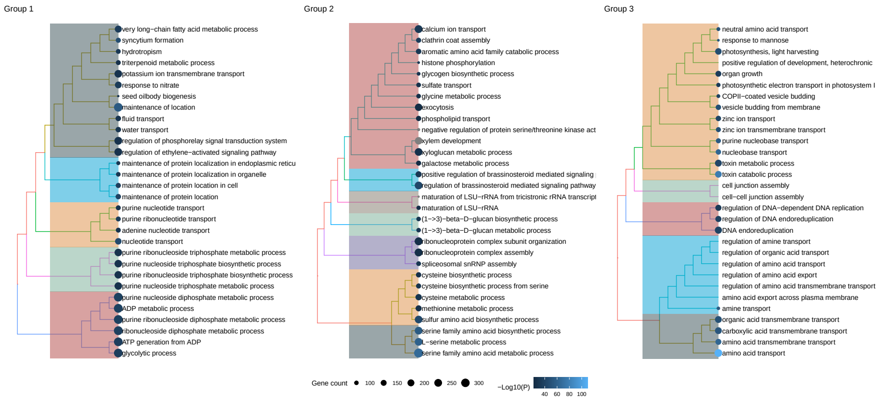

**Figure 8:** Tree plot of functional terms associated with each orthogroup cluster.

```
# Combine plots in one, keep shared legend
rcol <- range(
    c(
        p_dot_g1$data$p.adjust, p_dot_g2$data$p.adjust,
        p_dot_g3$data$p.adjust
    ),
    na.rm = TRUE
)
rsize <- range(
    c(
        p_dot_g1$data$Count, p_dot_g2$data$Count, p_dot_g3$data$Count
    ),
    na.rm = TRUE
)

wrap_plots(p_dot_g1, p_dot_g2, p_dot_g3) +
    plot_layout(guides = "collect") &
    scale_color_continuous(name = "-Log10(P)", limits = signif(rcol, 2)) &
    scale_size_continuous(name = "Gene count", limits = rsize) &
    theme(legend.position = "bottom")
```

The plots show that genes associated to particular biological processes tend to be clustered in the same orthogroup (group 3, scores closer to 1), while genes associated to other biological processes tend to be more dispersed across orthogroups (groups 1 and 2, scores closer to 1), possibly because they are evolving faster and, hence, have lower sequence similarity among themselves. In details, these genes and processes are:

- **Group 1:** ATP production, water and K+ transport, seed oilbody biogenesis, and response to nitrate and ethylene.
- **Group 2:** sulfur amino acid metabolsm, spliceosome biogenesis, beta-1,3-glucan biosynthesis, response to brassinosteroids, xylem development, exocytosis, and calcium and sulfate transport.
- **Group 3:** photosynthesis, zinc and amino acid transport, DNA replication, endocytosis, cell-cell junction assembly, and toxin catabolism.

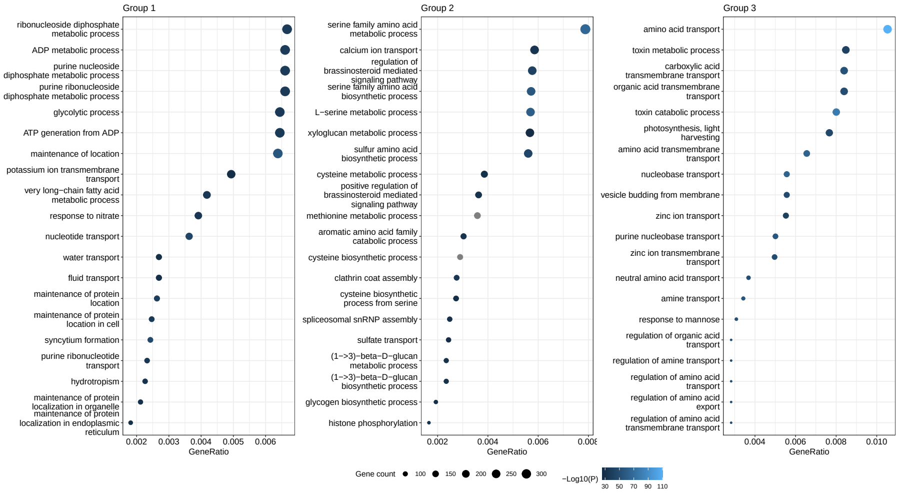

**Figure 9:** Dotplot of functional terms associated with each orthogroup cluster.

### 6 Is there an association between OG score and OG gene length?

Emms and Kelly (2015) have demonstrated a gene length bias that influences the accuracy of orthogroup detection. This is because short sequences cannot produce large bit scores or low e-values, and long sequences produce many hits with scores better than those for the best hits of short sequences (Emms and Kelly 2015). OrthoFinder implements a score transform that claims to eliminate such bias. But does it remove the bias completely?

To answer this question, we will use homogeneity scores for the default OrthoFinder run (default DIAMOND mode, mcl = 1.5).

First of all, let's calculate the mean and median gene length for each orthogroup.

```
# Combine proteomes into a single AAStringSet object and clean gene names
names(brassicaceae_proteomes) <- NULL
proteomes <- do.call(c, brassicaceae_proteomes)
rm(brassicaceae_proteomes)

names(proteomes) <- gsub("\\\t.*", "", names(proteomes))
names(proteomes) <- gsub(" .*", "", names(proteomes))
names(proteomes) <- gsub("\\.[0-9]$", "", names(proteomes))
names(proteomes) <- gsub("\\.[0-9]\\.p$", "", names(proteomes))
names(proteomes) <- gsub("\\.t[0-9]$", "", names(proteomes))
names(proteomes) <- gsub("\\.g$", "", names(proteomes))

# Load only orthogroups from the default OrthoFinder run
og <- read_orthogroups(file.path(tempdir(), "Orthogroups_default_1_5.tsv")) %>%
    mutate(Gene = str_replace_all(
        Gene, c(
            "\\\t.*" = "",
```

```
                "\\.[0-9]$" = "",
                "\\.[0-9]\\.p$" = "",
                "\\.t[0-9]$" = "",
                "\\.g$" = ""
            )
        )) %>%
        dplyr::select(Orthogroup, Gene)

# Calculate mean gene lengths for each orthogroup
gene_lengths <- data.frame(
    Gene = names(proteomes),
    Length = Biostrings::width(proteomes)
)

og_gene_lengths <- og %>%
    inner_join(., gene_lengths) %>%
    group_by(Orthogroup) %>%
    summarise(
        mean_length = mean(Length),
        median_length = median(Length)
    )

# Add homogeneity scores to data frame of mean gene length per orthogroup
og_length_and_scores <- og_assessment %>%
    dplyr::filter(Mode == "default_1_5") %>%
    dplyr::select(Orthogroups, Mean_H, Median_H) %>%
    inner_join(., og_gene_lengths, by = c("Orthogroups" = "Orthogroup"))

# Save data
save(
    og_length_and_scores,
    file = here("products", "result_files", "og_length_and_scores.rda"),
    compress = "xz"
)
```

Now, let's plot the data.

```
p_association_length_homogeneity <- og_length_and_scores %>%
    mutate(
        logH = log10(Median_H),
        logLength = log10(median_length)
    ) %>%
    ggscatter(
        ., x = "logLength", y = "logH", alpha = 0.3,
        color = "black", size = 1,
        add = "reg.line", add.params = list(color = "blue", fill = "lightgray"),
        conf.int = TRUE,
        cor.coef = TRUE,
        cor.coeff.args = list(
            method = "spearman", label.x = 1.7, label.y = -1, label.sep = "\n"
        )
```

```
    ) +
    labs(
        title = "Relationship between OG homogeneity score and gene length",
        x = expression(Log[10]~"median gene length"),
        y = expression(Log[10]~"median homogeneity score")
    )

p_association_length_homogeneity
```

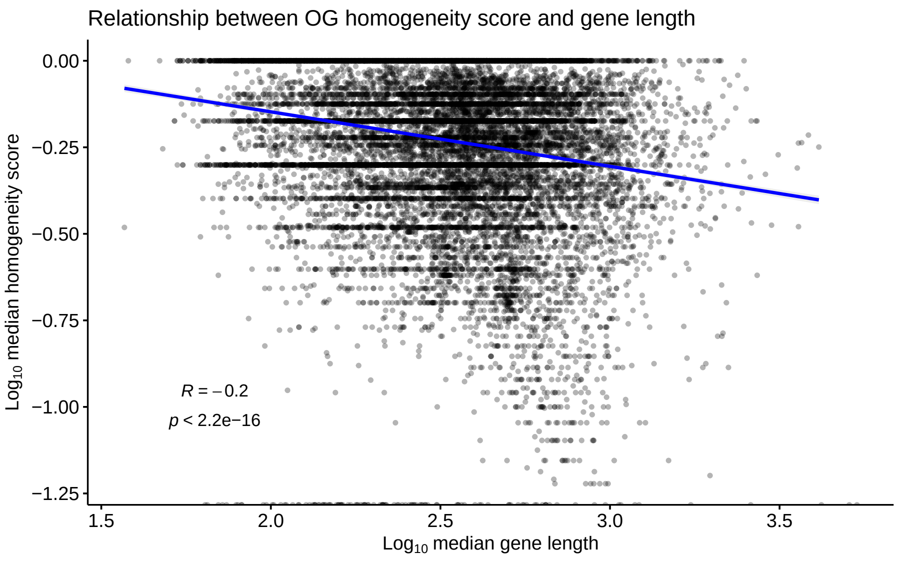

**Figure 10:** Relationship between sequence length and orthogroup scores.

There is a significant, negative association between homogeneity and OG gene length, but the strength of association is small ($R^2 = -0.2$). Although weak, this negative relationship shows that OrthoFinder can reduce the gene length bias, but not completely.

### Session info

This document was created under the following conditions:

```
sessioninfo::session_info()
## - Session info ---------------------------------------------------------------
##  setting  value
##  version  R version 4.2.2 Patched (2022-11-10 r83330)
##  os       Ubuntu 20.04.5 LTS
##  system   x86_64, linux-gnu
##  ui       X11
##  language (EN)
##  collate  en_US.UTF-8
##  ctype    en_US.UTF-8
##  tz       Europe/Brussels
##  date     2023-02-06
```

```
##  pandoc   2.19.2 @ /usr/lib/rstudio/resources/app/bin/quarto/bin/tools/ (via rmarkdown)
##
## - Packages -------------------------------------------------------------------
##  package          * version   date (UTC) lib source
##  abind              1.4-5     2016-07-21 [1] CRAN (R 4.2.0)
##  AnnotationDbi      1.58.0    2022-04-26 [1] Bioconductor
##  ape                5.6-2     2022-03-02 [1] CRAN (R 4.2.0)
##  aplot              0.1.8     2022-10-09 [1] CRAN (R 4.2.1)
##  assertthat         0.2.1     2019-03-21 [1] CRAN (R 4.2.0)
##  backports          1.4.1     2021-12-13 [1] CRAN (R 4.2.0)
##  beeswarm           0.4.0     2021-06-01 [1] CRAN (R 4.2.2)
##  Biobase            2.56.0    2022-04-26 [1] Bioconductor
##  BiocGenerics       0.42.0    2022-04-26 [1] Bioconductor
##  BiocManager        1.30.18   2022-05-18 [1] CRAN (R 4.2.0)
##  BiocParallel       1.30.4    2022-10-11 [1] Bioconductor
##  BiocStyle        * 2.25.0    2022-06-15 [1] Github (Bioconductor/BiocStyle@7150c28)
##  Biostrings         2.64.1    2022-08-18 [1] Bioconductor
##  bit                4.0.4     2020-08-04 [1] CRAN (R 4.2.0)
##  bit64              4.0.5     2020-08-30 [1] CRAN (R 4.2.0)
##  bitops             1.0-7     2021-04-24 [1] CRAN (R 4.2.0)
##  blob               1.2.3     2022-04-10 [1] CRAN (R 4.2.0)
##  bookdown           0.29      2022-09-12 [1] CRAN (R 4.2.1)
##  broom              1.0.1     2022-08-29 [1] CRAN (R 4.2.1)
##  cachem             1.0.6     2021-08-19 [1] CRAN (R 4.2.0)
##  car                3.1-0     2022-06-15 [1] CRAN (R 4.2.0)
##  carData            3.0-5     2022-01-06 [1] CRAN (R 4.2.0)
##  cellranger         1.1.0     2016-07-27 [1] CRAN (R 4.2.0)
##  cli                3.4.1     2022-09-23 [1] CRAN (R 4.2.1)
##  clusterProfiler  * 4.4.4     2022-06-21 [1] Bioconductor
##  codetools          0.2-18    2020-11-04 [1] CRAN (R 4.2.0)
##  cogeqc           * 1.3.1     2023-01-24 [1] Bioconductor
##  coin               1.4-2     2021-10-08 [1] CRAN (R 4.2.1)
##  colorspace         2.0-3     2022-02-21 [1] CRAN (R 4.2.0)
##  crayon             1.5.2     2022-09-29 [1] CRAN (R 4.2.1)
##  data.table         1.14.2    2021-09-27 [1] CRAN (R 4.2.0)
##  DBI                1.1.3     2022-06-18 [1] CRAN (R 4.2.0)
##  dbplyr             2.2.1     2022-06-27 [1] CRAN (R 4.2.1)
##  digest             0.6.29    2021-12-01 [1] CRAN (R 4.2.0)
##  DO.db              2.9       2022-06-28 [1] Bioconductor
##  DOSE               3.22.1    2022-08-30 [1] Bioconductor
##  downloader         0.4       2015-07-09 [1] CRAN (R 4.2.0)
##  dplyr            * 1.0.10    2022-09-01 [1] CRAN (R 4.2.1)
##  ellipsis           0.3.2     2021-04-29 [1] CRAN (R 4.2.0)
##  enrichplot       * 1.17.3    2022-10-26 [1] Github (YuLab-SMU/enrichplot@c33d502)
##  evaluate           0.17      2022-10-07 [1] CRAN (R 4.2.1)
##  fansi              1.0.3     2022-03-24 [1] CRAN (R 4.2.0)
##  farver             2.1.1     2022-07-06 [1] CRAN (R 4.2.1)
##  fastmap            1.1.0     2021-01-25 [1] CRAN (R 4.2.0)
##  fastmatch          1.1-3     2021-07-23 [1] CRAN (R 4.2.0)
##  fgsea              1.22.0    2022-04-26 [1] Bioconductor
##  forcats          * 0.5.2     2022-08-19 [1] CRAN (R 4.2.1)
```

```
##  fs                  1.5.2     2021-12-08 [1] CRAN (R 4.2.0)
##  gargle              1.2.1     2022-09-08 [1] CRAN (R 4.2.1)
##  generics            0.1.3     2022-07-05 [1] CRAN (R 4.2.1)
##  GenomeInfoDb        1.32.4    2022-09-06 [1] Bioconductor
##  GenomeInfoDbData    1.2.8     2022-05-06 [1] Bioconductor
##  ggbeeswarm          0.7.1     2022-12-16 [1] CRAN (R 4.2.2)
##  ggforce             0.4.1     2022-10-04 [1] CRAN (R 4.2.1)
##  ggfun               0.0.8     2022-11-07 [1] CRAN (R 4.2.1)
##  ggnewscale          0.4.8     2022-10-06 [1] CRAN (R 4.2.1)
##  ggplot2           * 3.4.0     2022-11-04 [1] CRAN (R 4.2.1)
##  ggplotify           0.1.0     2021-09-02 [1] CRAN (R 4.2.0)
##  ggpubr            * 0.4.0     2020-06-27 [1] CRAN (R 4.2.0)
##  ggraph              2.1.0     2022-10-09 [1] CRAN (R 4.2.1)
##  ggrepel             0.9.1     2021-01-15 [1] CRAN (R 4.2.0)
##  ggsci               2.9       2018-05-14 [1] CRAN (R 4.2.0)
##  ggsignif            0.6.4     2022-10-13 [1] CRAN (R 4.2.1)
##  ggtext              0.1.2     2022-09-16 [1] CRAN (R 4.2.1)
##  ggtree              3.7.1.001 2022-11-10 [1] Github (YuLab-SMU/ggtree@b7ef83e)
##  glue                1.6.2     2022-02-24 [1] CRAN (R 4.2.0)
##  GO.db               3.15.0    2022-05-06 [1] Bioconductor
##  googledrive         2.0.0     2021-07-08 [1] CRAN (R 4.2.0)
##  googlesheets4       1.0.1     2022-08-13 [1] CRAN (R 4.2.1)
##  GOSemSim            2.22.0    2022-04-26 [1] Bioconductor
##  graphlayouts        0.8.2     2022-09-29 [1] CRAN (R 4.2.1)
##  gridExtra           2.3       2017-09-09 [1] CRAN (R 4.2.0)
##  gridGraphics        0.5-1     2020-12-13 [1] CRAN (R 4.2.0)
##  gridtext            0.1.5     2022-09-16 [1] CRAN (R 4.2.1)
##  gtable              0.3.1     2022-09-01 [1] CRAN (R 4.2.1)
##  haven               2.5.1     2022-08-22 [1] CRAN (R 4.2.1)
##  here              * 1.0.1     2020-12-13 [1] CRAN (R 4.2.0)
##  hms                 1.1.2     2022-08-19 [1] CRAN (R 4.2.1)
##  htmltools           0.5.3     2022-07-18 [1] CRAN (R 4.2.1)
##  httr                1.4.4     2022-08-17 [1] CRAN (R 4.2.1)
##  igraph              1.3.5     2022-09-22 [1] CRAN (R 4.2.1)
##  IRanges             2.30.1    2022-08-18 [1] Bioconductor
##  jsonlite            1.8.3     2022-10-21 [1] CRAN (R 4.2.1)
##  KEGGREST            1.36.3    2022-07-12 [1] Bioconductor
##  knitr               1.40      2022-08-24 [1] CRAN (R 4.2.1)
##  labeling            0.4.2     2020-10-20 [1] CRAN (R 4.2.0)
##  lattice             0.20-45   2021-09-22 [1] CRAN (R 4.2.0)
##  lazyeval            0.2.2     2019-03-15 [1] CRAN (R 4.2.0)
##  libcoin             1.0-9     2021-09-27 [1] CRAN (R 4.2.1)
##  lifecycle           1.0.3     2022-10-07 [1] CRAN (R 4.2.1)
##  lubridate           1.8.0     2021-10-07 [1] CRAN (R 4.2.0)
##  magrittr            2.0.3     2022-03-30 [1] CRAN (R 4.2.0)
##  markdown            1.1       2019-08-07 [1] CRAN (R 4.2.0)
##  MASS                7.3-58.1  2022-08-03 [1] CRAN (R 4.2.1)
##  Matrix              1.5-1     2022-09-13 [1] CRAN (R 4.2.1)
##  matrixStats         0.62.0    2022-04-19 [1] CRAN (R 4.2.0)
##  memoise             2.0.1     2021-11-26 [1] CRAN (R 4.2.0)
##  mgcv                1.8-40    2022-03-29 [1] CRAN (R 4.2.0)
```

```
## modelr          0.1.9    2022-08-19 [1] CRAN (R 4.2.1)
## modeltools      0.2-23   2020-03-05 [1] CRAN (R 4.2.1)
## multcomp        1.4-20   2022-08-07 [1] CRAN (R 4.2.1)
## munsell         0.5.0    2018-06-12 [1] CRAN (R 4.2.0)
## mvtnorm         1.1-3    2021-10-08 [1] CRAN (R 4.2.0)
## nlme            3.1-160  2022-10-10 [1] CRAN (R 4.2.1)
## patchwork     * 1.1.2    2022-08-19 [1] CRAN (R 4.2.1)
## pillar          1.8.1    2022-08-19 [1] CRAN (R 4.2.1)
## pkgconfig       2.0.3    2019-09-22 [1] CRAN (R 4.2.0)
## plyr            1.8.7    2022-03-24 [1] CRAN (R 4.2.0)
## png             0.1-7    2013-12-03 [1] CRAN (R 4.2.0)
## polyclip        1.10-0   2019-03-14 [1] CRAN (R 4.2.0)
## purrr         * 0.3.5    2022-10-06 [1] CRAN (R 4.2.1)
## qvalue          2.28.0   2022-04-26 [1] Bioconductor
## R6              2.5.1    2021-08-19 [1] CRAN (R 4.2.0)
## RColorBrewer    1.1-3    2022-04-03 [1] CRAN (R 4.2.0)
## Rcpp            1.0.9    2022-07-08 [1] CRAN (R 4.2.1)
## RCurl           1.98-1.9 2022-10-03 [1] CRAN (R 4.2.1)
## readr         * 2.1.3    2022-10-01 [1] CRAN (R 4.2.1)
## readxl          1.4.1    2022-08-17 [1] CRAN (R 4.2.1)
## reprex          2.0.2    2022-08-17 [1] CRAN (R 4.2.1)
## reshape2        1.4.4    2020-04-09 [1] CRAN (R 4.2.0)
## rlang           1.0.6    2022-09-24 [1] CRAN (R 4.2.1)
## rmarkdown       2.17     2022-10-07 [1] CRAN (R 4.2.1)
## rprojroot       2.0.3    2022-04-02 [1] CRAN (R 4.2.0)
## RSQLite         2.2.18   2022-10-04 [1] CRAN (R 4.2.1)
## rstatix       * 0.7.0    2021-02-13 [1] CRAN (R 4.2.1)
## rstudioapi      0.14     2022-08-22 [1] CRAN (R 4.2.1)
## rvest           1.0.3    2022-08-19 [1] CRAN (R 4.2.1)
## S4Vectors       0.34.0   2022-04-26 [1] Bioconductor
## sandwich        3.0-2    2022-06-15 [1] CRAN (R 4.2.1)
## scales          1.2.1    2022-08-20 [1] CRAN (R 4.2.1)
## scatterpie      0.1.8    2022-09-03 [1] CRAN (R 4.2.1)
## sessioninfo     1.2.2    2021-12-06 [1] CRAN (R 4.2.0)
## shadowtext      0.1.2    2022-04-22 [1] CRAN (R 4.2.1)
## stringi         1.7.8    2022-07-11 [1] CRAN (R 4.2.1)
## stringr       * 1.4.1    2022-08-20 [1] CRAN (R 4.2.1)
## survival        3.4-0    2022-08-09 [1] CRAN (R 4.2.1)
## TH.data         1.1-1    2022-04-26 [1] CRAN (R 4.2.1)
## tibble        * 3.1.8    2022-07-22 [1] CRAN (R 4.2.1)
## tidygraph       1.2.2    2022-08-22 [1] CRAN (R 4.2.1)
## tidyr         * 1.2.1    2022-09-08 [1] CRAN (R 4.2.1)
## tidyselect      1.2.0    2022-10-10 [1] CRAN (R 4.2.1)
## tidytree        0.4.1    2022-09-26 [1] CRAN (R 4.2.1)
## tidyverse     * 1.3.2    2022-07-18 [1] CRAN (R 4.2.1)
## treeio          1.23.0   2022-11-10 [1] Github (GuangchuangYu/treeio@db85803)
## tweenr          2.0.2    2022-09-06 [1] CRAN (R 4.2.1)
## tzdb            0.3.0    2022-03-28 [1] CRAN (R 4.2.0)
## utf8            1.2.2    2021-07-24 [1] CRAN (R 4.2.0)
## vctrs           0.5.0    2022-10-22 [1] CRAN (R 4.2.1)
## vipor           0.4.5    2017-03-22 [1] CRAN (R 4.2.1)
```

```
##  viridis           0.6.2     2021-10-13 [1] CRAN (R 4.2.0)
##  viridisLite       0.4.1     2022-08-22 [1] CRAN (R 4.2.1)
##  withr             2.5.0     2022-03-03 [1] CRAN (R 4.2.0)
##  xfun              0.33      2022-09-12 [1] CRAN (R 4.2.1)
##  xml2              1.3.3     2021-11-30 [1] CRAN (R 4.2.0)
##  XVector           0.36.0    2022-04-26 [1] Bioconductor
##  yaml              2.3.5     2022-02-21 [1] CRAN (R 4.2.0)
##  yulab.utils       0.0.5     2022-06-30 [1] CRAN (R 4.2.1)
##  zlibbioc          1.42.0    2022-04-26 [1] Bioconductor
##  zoo               1.8-11    2022-09-17 [1] CRAN (R 4.2.1)
##
##  [1] /home/faalm/R/x86_64-pc-linux-gnu-library/4.2
##  [2] /usr/local/lib/R/site-library
##  [3] /usr/lib/R/site-library
##  [4] /usr/lib/R/library
##
## ------------------------------------------------------------------------------
```

### References

Emms, David M, and Steven Kelly. 2015. "OrthoFinder: Solving Fundamental Biases in Whole Genome Comparisons Dramatically Improves Orthogroup Inference Accuracy." *Genome Biology* 16 (1): 1–14.

———. 2019. "OrthoFinder: Phylogenetic Orthology Inference for Comparative Genomics." *Genome Biology* 20 (1): 1–14.
