## Supplementary Text S4 for "Assessing the quality of comparative genomics data and results with the *cogeqc* R/Bioconductor package"

### Supplementary Text S4: Assessing synteny detection in Fabaceae

***Fabricio Almeida-Silva*** [1,2] ***and Yves Van de Peer*** [1,2,3,4]

[1]VIB-UGent Center for Plant Systems Biology, Ghent, Belgium
[2]Department of Plant Biotechnology and Bioinformatics, Ghent University, Ghent, Belgium
[3]College of Horticulture, Academy for Advanced Interdisciplinary Studies, Nanjing Agricultural University, Nanjing, China
[4]Center for Microbial Ecology and Genomics, Department of Biochemistry, Genetics and Microbiology, University of Pretoria, Pretoria, South Africa

**6 February 2023**

```r
library(cogeqc)
library(here)
library(tidyverse)
library(syntenet)
library(tidytext)
```

### 1 Overview

Here, we will assess synteny detection using a network-based approach. The anchor pairs from synteny identification will be interpreted as edges of an unweighted undirected graph (i.e., a synteny network), and the best synteny detection will be identified based on the graphs' clustering coefficients and node number.

We will demonstrate our network-based synteny assessment using genomic data on Fabaceae species available on PLAZA 5.0 (Van Bel et al. 2022).

### 2 Data acquisition

In this section, we will download whole-genome protein sequences and gene annotation from PLAZA 5.0, and then we will preprocess the data with `syntenet::process_input()`.

```r
species <- c("mtr", "tpr", "psa", "car", "lja", "gma", "vmu", "lal", "arhy")
```

```r
base_url <- "https://ftp.psb.ugent.be/pub/plaza/plaza_public_dicots_05/"

# Get proteomes
seq_url <- paste0(
    base_url, "Fasta/proteome.selected_transcript.",
    species, ".fasta.gz"
)

## Import files and clean gene IDs
seq <- lapply(seq_url, function(x) {
    s <- Biostrings::readAAStringSet(x)
    names(s) <- gsub(".* | ", "", names(s))
    return(s)
})
names(seq) <- species

# Get gene annotation
annot_url <- paste0(
    base_url, "GFF/", species, "/annotation.selected_transcript.exon_features.",
    species, ".gff3.gz"
)

## Import files and keep only relevant fields
annot <- lapply(annot_url, function(x) {
    a <- rtracklayer::import(x)
```

```
    a <- a[, c("type", "gene_id")]
    a <- a[a$type == "gene"]
    return(a)
})
names(annot) <- species

# Process data
pdata <- process_input(seq, annot)

# Remove unprocessed data to clean the working environment
rm(annot)
rm(seq)
```

### 3 Network-based synteny assessment

We will infer synteny networks using the Bioconductor package *syntenet*. This package detects synteny using the MCScanX algorithm (Wang et al. 2012), which can produce different results based on 2 main parameters:

1. **anchors:** minimum required number of genes to call a syntenic block. Default: 5.
2. **max_gaps:** number of upstream and downstream genes to search for anchors. Default: 25.

We will infer synteny networks with 5 combinations of parameters, similarly to Zhao and Schranz (2019), using two approaches:

1. A single Fabaceae synteny network;
2. Species-specific synteny networks for each Fabaceae species.

To start with, let's define the combinations of parameters we will use.

```
# Define combinations of parameters: anchors (a), max_gaps (m)
synteny_params <- list(
    c(3, 25),
    c(5, 15),
    c(5, 25),
    c(5, 35),
    c(7, 25)
)
```

#### 3.1 Assessing the Fabaceae synteny network

First, we will perform similarity searches with DIAMOND.

```
# Define wrapper function to run DIAMOND with different top_hits
out <- file.path(tempdir(), "diamond_all")
d5 <- run_diamond(seq = seq, top_hits = 5, outdir = out)
```

With the DIAMOND list, we can detect synteny.

```r
# Define helper function to detect synteny with multiple combinations of params
synteny_wrapper <- function(diamond, annotation, params) {

    syn <- lapply(params, function(x) {

        anchors <- x[1]
        max_gaps <- x[2]
        outdir <- file.path(tempdir(), paste0("syn_a", anchors, "_m", max_gaps))

        s <- infer_syntenet(
            blast_list = diamond,
            annotation = pdata$annotation,
            outdir = outdir,
            anchors = anchors,
            max_gaps = max_gaps
        )
        return(s)
    })
    return(syn)
}

# Detect synteny
syn_fabaceae <- synteny_wrapper(d5, pdata$annotation, synteny_params)
names(syn_fabaceae) <- unlist(
    lapply(synteny_params, function(x) paste0("a", x[1], "_m", x[2]))
)
```

Now, let's use the network-based synteny assessment to see which combination of parameters is the best.

```r
# Assess networks
fabaceae_scores <- assess_synnet_list(syn_fabaceae)

# Look at scores, ranked from highest to lowest
fabaceae_scores %>%
    arrange(-Score) |>
    knitr::kable(
        caption = "Scores for each synteny network."
    )
```

**Table 1:** Scores for each synteny network.

| CC | Node_count | Rsquared | Score | Network |
|---:|---:|---:|---:|:---|
| 0.8253002 | 237723 | 0.6227916 | 122187.2 | a3_m25 |
| 0.8290880 | 235290 | 0.6156847 | 120105.4 | a5_m35 |
| 0.8392223 | 226657 | 0.6026291 | 114629.5 | a5_m15 |
| 0.8412602 | 224325 | 0.5972865 | 112717.3 | a7_m25 |
| 0.8347725 | 231820 | 0.5795957 | 112161.6 | a5_m25 |

As we can see, the combination of parameters `a = 3; m = 25` is the best for this data set.

Finally, let's visualize scores. To make visualization better, we will scale scores by the maximum value, so that values range from 0 to 1.

```
# Plot scores
synteny_scores_fabaceae <- fabaceae_scores %>%
    arrange(Score) %>%
    mutate(Score = Score / max(Score)) %>%
    mutate(Parameters = str_replace_all(Network, "_", ", ")) %>%
    mutate(Parameters = factor(Parameters, levels = unique(Parameters))) %>%
    ggplot(., aes(x = Parameters, y = Score)) +
    geom_col(fill = "grey60", color = "black") +
    theme_bw() +
    labs(
        title = "Assessment of the Fabaceae synteny network",
        subtitle = "a = minimum # of anchors; m = maximum # of gaps",
        y = "Scaled score"
    )

synteny_scores_fabaceae
```

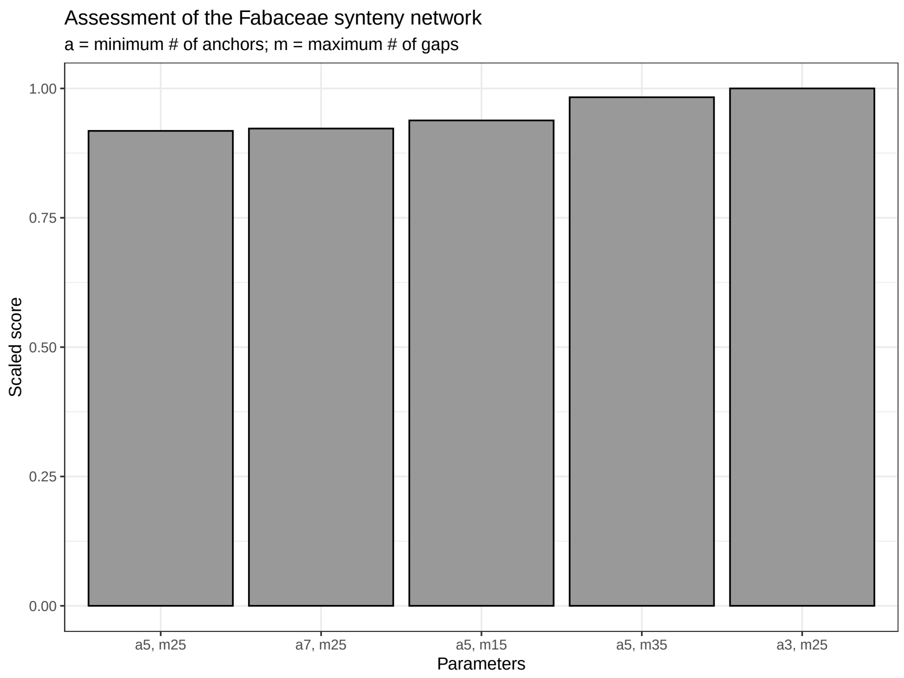

**Figure 1:** Scaled scores for each synteny network.

#### 3.2 Assessing species-specific synteny networks

In this section, we will infer species-specific synteny networks and assess each of them with our network-based approach.

This time, as we already have synteny networks for the whole Fabaceae family, we don't need to infer them again; we will simply subset edges of the network that contain nodes from the same species.

```r
# Create species-specific networks
species_ids <- substr(species, start = 1, stop = 3)

species_networks <- lapply(species_ids, function(x) {

    nets <- lapply(syn_fabaceae, function(y) {
        edges <- y[startsWith(y$Anchor1, x) & startsWith(y$Anchor2, x), ]
        return(edges)
    })
    return(nets)
})
names(species_networks) <- species_ids

# Exploring data
names(species_networks)
## [1] "mtr" "tpr" "psa" "car" "lja" "gma" "vmu" "lal" "arh"
names(species_networks$mtr)
## [1] "a3_m25" "a5_m15" "a5_m25" "a5_m35" "a7_m25"

# Rename `species_networks` to keep full name
names(species_networks) <- c(
    "M. truncatula", "T. pratense", "P. sativum", "C. arietinum",
    "L. japonicus", "G. max", "V. mungo", "L. albus", "A. hypogaea"
)
```

For each species, we will assess the networks inferred with different combinations of parameters.

```r
# Assess species-specific networks
scores_species_nets <- lapply(seq_along(species_networks), function(x) {

    species <- names(species_networks)[x]
    scores <- assess_synnet_list(species_networks[[species]])
    scores$Score[is.nan(scores$Score)] <- 0
    scores <- scores[order(scores$Score, decreasing = TRUE), ]
    scores$Species <- species
    scores$Score <- scores$Score / max(scores$Score)
    return(scores)
})
scores_species_nets <- Reduce(rbind, scores_species_nets)

# Plot data
synteny_scores_species <- scores_species_nets %>%
    mutate(
        Parameters = as.factor(str_replace_all(Network, "_", ", ")),
        Species = as.factor(Species)
    ) %>%
    mutate(Network = reorder_within(Parameters, Score, Species)) %>%
    ggplot(., aes(x = Network, y = Score, fill = Parameters)) +
    geom_bar(stat = "identity", color = "grey90") +
    facet_wrap(~Species, ncol = 3, scales = "free") +
```

```
    scale_x_reordered() +
    ggsci::scale_fill_jama() +
    theme_bw() +
    theme(axis.text.x = element_blank()) +
    labs(
        title = "Assessment of species-specific synteny networks",
        subtitle = "a = minimum # of anchors; m = maximum # of gaps",
        y = "Scaled score (by species)", x = ""
    )

synteny_scores_species
```

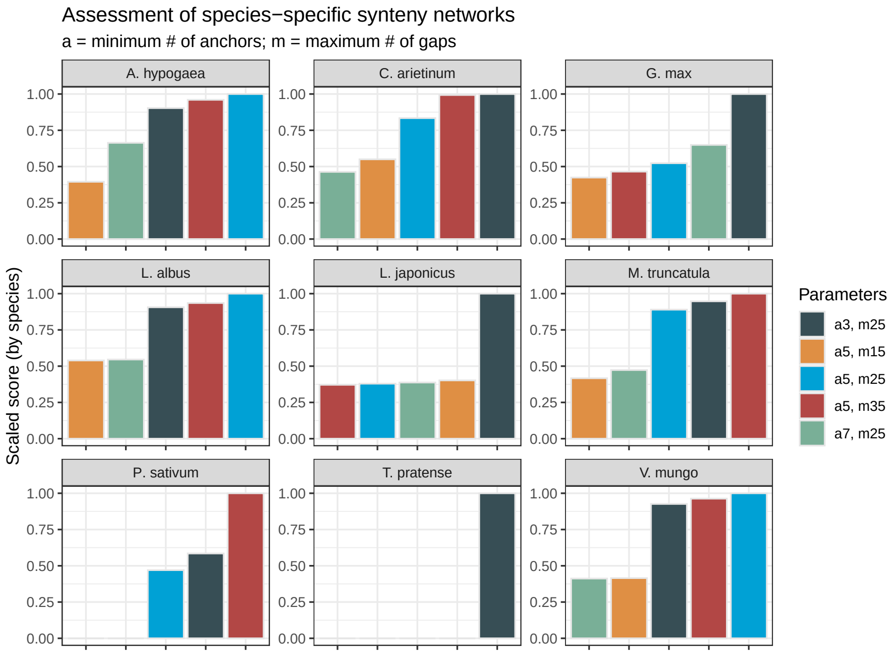

**Figure 2:** Scores for species-specific synteny networks.

The figure demonstrates that the best combination of parameters depends on the species, so there is no "universally" best combination. However, some patterns emerge. The combinations `a = 7; m= 25` and `a = 5; m = 15` are typically the worst. In some cases, they even lead to zero scores due to clustering coefficients of zero. Thus, if users want to test multiple combinations of parameters for their own data set, they should only test the combinations `a = 3; m = 25`, `a = 5; m = 25`, and `a = 5; m = 35`, which lead to the best score in 45%, 33%, and 22% of the species-specific networks, respectively. Interestingly, the combination that leads to the best score in most networks (`a = 3; m = 25`) is also the best when considering the whole Fabaceae synteny network (see previous section).

### Session info

This document was created under the following conditions:

```
## - Session info ---------------------------------------------------------------
##  setting  value
##  version  R version 4.2.2 Patched (2022-11-10 r83330)
##  os       Ubuntu 20.04.5 LTS
##  system   x86_64, linux-gnu
##  ui       X11
##  language (EN)
##  collate  en_US.UTF-8
##  ctype    en_US.UTF-8
##  tz       Europe/Brussels
##  date     2023-02-06
##  pandoc   2.19.2 @ /usr/lib/rstudio/resources/app/bin/quarto/bin/tools/ (via rmarkdown)
##
## - Packages -------------------------------------------------------------------
##  package              * version   date (UTC) lib source
##  ape                    5.6-2     2022-03-02 [1] CRAN (R 4.2.0)
##  aplot                  0.1.8     2022-10-09 [1] CRAN (R 4.2.1)
##  assertthat             0.2.1     2019-03-21 [1] CRAN (R 4.2.0)
##  backports              1.4.1     2021-12-13 [1] CRAN (R 4.2.0)
##  beeswarm               0.4.0     2021-06-01 [1] CRAN (R 4.2.2)
##  Biobase                2.56.0    2022-04-26 [1] Bioconductor
##  BiocGenerics           0.42.0    2022-04-26 [1] Bioconductor
##  BiocIO                 1.6.0     2022-04-26 [1] Bioconductor
##  BiocManager            1.30.18   2022-05-18 [1] CRAN (R 4.2.0)
##  BiocParallel           1.30.4    2022-10-11 [1] Bioconductor
##  BiocStyle            * 2.25.0    2022-06-15 [1] Github (Bioconductor/BiocStyle@7150c28)
##  Biostrings             2.64.1    2022-08-18 [1] Bioconductor
##  bitops                 1.0-7     2021-04-24 [1] CRAN (R 4.2.0)
##  bookdown               0.29      2022-09-12 [1] CRAN (R 4.2.1)
##  broom                  1.0.1     2022-08-29 [1] CRAN (R 4.2.1)
##  cellranger             1.1.0     2016-07-27 [1] CRAN (R 4.2.0)
##  cli                    3.4.1     2022-09-23 [1] CRAN (R 4.2.1)
##  coda                   0.19-4    2020-09-30 [1] CRAN (R 4.2.0)
##  codetools              0.2-18    2020-11-04 [1] CRAN (R 4.2.0)
##  cogeqc               * 1.3.1     2023-01-24 [1] Bioconductor
##  colorspace             2.0-3     2022-02-21 [1] CRAN (R 4.2.0)
##  crayon                 1.5.2     2022-09-29 [1] CRAN (R 4.2.1)
##  DBI                    1.1.3     2022-06-18 [1] CRAN (R 4.2.0)
##  dbplyr                 2.2.1     2022-06-27 [1] CRAN (R 4.2.1)
##  DelayedArray           0.22.0    2022-04-26 [1] Bioconductor
##  digest                 0.6.29    2021-12-01 [1] CRAN (R 4.2.0)
##  dplyr                * 1.0.10    2022-09-01 [1] CRAN (R 4.2.1)
##  ellipsis               0.3.2     2021-04-29 [1] CRAN (R 4.2.0)
##  evaluate               0.17      2022-10-07 [1] CRAN (R 4.2.1)
##  fansi                  1.0.3     2022-03-24 [1] CRAN (R 4.2.0)
##  farver                 2.1.1     2022-07-06 [1] CRAN (R 4.2.1)
##  fastmap                1.1.0     2021-01-25 [1] CRAN (R 4.2.0)
##  forcats              * 0.5.2     2022-08-19 [1] CRAN (R 4.2.1)
##  fs                     1.5.2     2021-12-08 [1] CRAN (R 4.2.0)
##  gargle                 1.2.1     2022-09-08 [1] CRAN (R 4.2.1)
##  generics               0.1.3     2022-07-05 [1] CRAN (R 4.2.1)
```

```
##  GenomeInfoDb            1.32.4    2022-09-06 [1] Bioconductor
##  GenomeInfoDbData        1.2.8     2022-05-06 [1] Bioconductor
##  GenomicAlignments       1.32.1    2022-07-24 [1] Bioconductor
##  GenomicRanges           1.48.0    2022-04-26 [1] Bioconductor
##  ggbeeswarm              0.7.1     2022-12-16 [1] CRAN (R 4.2.2)
##  ggfun                   0.0.8     2022-11-07 [1] CRAN (R 4.2.1)
##  ggnetwork               0.5.10    2021-07-06 [1] CRAN (R 4.2.0)
##  ggplot2               * 3.4.0     2022-11-04 [1] CRAN (R 4.2.1)
##  ggplotify               0.1.0     2021-09-02 [1] CRAN (R 4.2.0)
##  ggsci                   2.9       2018-05-14 [1] CRAN (R 4.2.0)
##  ggtree                  3.7.1.001 2022-11-10 [1] Github (YuLab-SMU/ggtree@b7ef83e)
##  glue                    1.6.2     2022-02-24 [1] CRAN (R 4.2.0)
##  googledrive             2.0.0     2021-07-08 [1] CRAN (R 4.2.0)
##  googlesheets4           1.0.1     2022-08-13 [1] CRAN (R 4.2.1)
##  gridGraphics            0.5-1     2020-12-13 [1] CRAN (R 4.2.0)
##  gtable                  0.3.1     2022-09-01 [1] CRAN (R 4.2.1)
##  haven                   2.5.1     2022-08-22 [1] CRAN (R 4.2.1)
##  here                  * 1.0.1     2020-12-13 [1] CRAN (R 4.2.0)
##  hms                     1.1.2     2022-08-19 [1] CRAN (R 4.2.1)
##  htmltools               0.5.3     2022-07-18 [1] CRAN (R 4.2.1)
##  htmlwidgets             1.5.4     2021-09-08 [1] CRAN (R 4.2.0)
##  httr                    1.4.4     2022-08-17 [1] CRAN (R 4.2.1)
##  igraph                  1.3.5     2022-09-22 [1] CRAN (R 4.2.1)
##  intergraph              2.0-2     2016-12-05 [1] CRAN (R 4.2.0)
##  IRanges                 2.30.1    2022-08-18 [1] Bioconductor
##  janeaustenr             1.0.0     2022-08-26 [1] CRAN (R 4.2.1)
##  jsonlite                1.8.3     2022-10-21 [1] CRAN (R 4.2.1)
##  knitr                   1.40      2022-08-24 [1] CRAN (R 4.2.1)
##  labeling                0.4.2     2020-10-20 [1] CRAN (R 4.2.0)
##  lattice                 0.20-45   2021-09-22 [1] CRAN (R 4.2.0)
##  lazyeval                0.2.2     2019-03-15 [1] CRAN (R 4.2.0)
##  lifecycle               1.0.3     2022-10-07 [1] CRAN (R 4.2.1)
##  lubridate               1.8.0     2021-10-07 [1] CRAN (R 4.2.0)
##  magrittr                2.0.3     2022-03-30 [1] CRAN (R 4.2.0)
##  Matrix                  1.5-1     2022-09-13 [1] CRAN (R 4.2.1)
##  MatrixGenerics          1.8.1     2022-06-26 [1] Bioconductor
##  matrixStats             0.62.0    2022-04-19 [1] CRAN (R 4.2.0)
##  modelr                  0.1.9     2022-08-19 [1] CRAN (R 4.2.1)
##  munsell                 0.5.0     2018-06-12 [1] CRAN (R 4.2.0)
##  network                 1.18.0    2022-10-06 [1] CRAN (R 4.2.1)
##  networkD3               0.4       2017-03-18 [1] CRAN (R 4.2.0)
##  nlme                    3.1-160   2022-10-10 [1] CRAN (R 4.2.1)
##  patchwork               1.1.2     2022-08-19 [1] CRAN (R 4.2.1)
##  pheatmap                1.0.12    2019-01-04 [1] CRAN (R 4.2.0)
##  pillar                  1.8.1     2022-08-19 [1] CRAN (R 4.2.1)
##  pkgconfig               2.0.3     2019-09-22 [1] CRAN (R 4.2.0)
##  plyr                    1.8.7     2022-03-24 [1] CRAN (R 4.2.0)
##  purrr                 * 0.3.5     2022-10-06 [1] CRAN (R 4.2.1)
##  R6                      2.5.1     2021-08-19 [1] CRAN (R 4.2.0)
##  RColorBrewer            1.1-3     2022-04-03 [1] CRAN (R 4.2.0)
##  Rcpp                    1.0.9     2022-07-08 [1] CRAN (R 4.2.1)
```

```
## RCurl                    1.98-1.9  2022-10-03 [1] CRAN (R 4.2.1)
## readr                  * 2.1.3     2022-10-01 [1] CRAN (R 4.2.1)
## readxl                   1.4.1     2022-08-17 [1] CRAN (R 4.2.1)
## reprex                   2.0.2     2022-08-17 [1] CRAN (R 4.2.1)
## reshape2                 1.4.4     2020-04-09 [1] CRAN (R 4.2.0)
## restfulr                 0.0.15    2022-06-16 [1] CRAN (R 4.2.0)
## rjson                    0.2.21    2022-01-09 [1] CRAN (R 4.2.0)
## rlang                    1.0.6     2022-09-24 [1] CRAN (R 4.2.1)
## rmarkdown                2.17      2022-10-07 [1] CRAN (R 4.2.1)
## rprojroot                2.0.3     2022-04-02 [1] CRAN (R 4.2.0)
## Rsamtools                2.12.0    2022-04-26 [1] Bioconductor
## rstudioapi               0.14      2022-08-22 [1] CRAN (R 4.2.1)
## rtracklayer              1.57.0    2022-06-16 [1] Github (lawremi/rtracklayer@2bb0b40)
## rvest                    1.0.3     2022-08-19 [1] CRAN (R 4.2.1)
## S4Vectors                0.34.0    2022-04-26 [1] Bioconductor
## scales                   1.2.1     2022-08-20 [1] CRAN (R 4.2.1)
## sessioninfo              1.2.2     2021-12-06 [1] CRAN (R 4.2.0)
## SnowballC                0.7.0     2020-04-01 [1] CRAN (R 4.2.0)
## statnet.common           4.7.0     2022-09-08 [1] CRAN (R 4.2.1)
## stringi                  1.7.8     2022-07-11 [1] CRAN (R 4.2.1)
## stringr                * 1.4.1     2022-08-20 [1] CRAN (R 4.2.1)
## SummarizedExperiment     1.26.1    2022-04-29 [1] Bioconductor
## syntenet               * 1.1.3     2023-02-01 [1] Bioconductor
## tibble                 * 3.1.8     2022-07-22 [1] CRAN (R 4.2.1)
## tidyr                  * 1.2.1     2022-09-08 [1] CRAN (R 4.2.1)
## tidyselect               1.2.0     2022-10-10 [1] CRAN (R 4.2.1)
## tidytext               * 0.3.4     2022-08-20 [1] CRAN (R 4.2.1)
## tidytree                 0.4.1     2022-09-26 [1] CRAN (R 4.2.1)
## tidyverse              * 1.3.2     2022-07-18 [1] CRAN (R 4.2.1)
## tokenizers               0.2.3     2022-09-23 [1] CRAN (R 4.2.1)
## treeio                   1.23.0    2022-11-10 [1] Github (GuangchuangYu/treeio@db85803)
## tzdb                     0.3.0     2022-03-28 [1] CRAN (R 4.2.0)
## utf8                     1.2.2     2021-07-24 [1] CRAN (R 4.2.0)
## vctrs                    0.5.0     2022-10-22 [1] CRAN (R 4.2.1)
## vipor                    0.4.5     2017-03-22 [1] CRAN (R 4.2.1)
## withr                    2.5.0     2022-03-03 [1] CRAN (R 4.2.0)
## xfun                     0.33      2022-09-12 [1] CRAN (R 4.2.1)
## XML                      3.99-0.11 2022-10-03 [1] CRAN (R 4.2.1)
## xml2                     1.3.3     2021-11-30 [1] CRAN (R 4.2.0)
## XVector                  0.36.0    2022-04-26 [1] Bioconductor
## yaml                     2.3.5     2022-02-21 [1] CRAN (R 4.2.0)
## yulab.utils              0.0.5     2022-06-30 [1] CRAN (R 4.2.1)
## zlibbioc                 1.42.0    2022-04-26 [1] Bioconductor
##
## [1] /home/faalm/R/x86_64-pc-linux-gnu-library/4.2
## [2] /usr/local/lib/R/site-library
## [3] /usr/lib/R/site-library
## [4] /usr/lib/R/library
##
## ------------------------------------------------------------------------------
```

### References

Van Bel, Michiel, Francesca Silvestri, Eric M Weitz, Lukasz Kreft, Alexander Botzki, Frederik Coppens, and Klaas Vandepoele. 2022. "PLAZA 5.0: Extending the Scope and Power of Comparative and Functional Genomics in Plants." *Nucleic Acids Research* 50 (D1): D1468–74.

Wang, Yupeng, Haibao Tang, Jeremy D DeBarry, Xu Tan, Jingping Li, Xiyin Wang, Tae-ho Lee, et al. 2012. "MCScanX: A Toolkit for Detection and Evolutionary Analysis of Gene Synteny and Collinearity." *Nucleic Acids Research* 40 (7): e49–49.

Zhao, Tao, and M Eric Schranz. 2019. "Network-Based Microsynteny Analysis Identifies Major Differences and Genomic Outliers in Mammalian and Angiosperm Genomes." *Proceedings of the National Academy of Sciences* 116 (6): 2165–74.
